# Predictive phage therapy for *Escherichia coli* urinary tract infections: cocktail selection for therapy based on machine learning models

**DOI:** 10.1101/2023.11.23.568453

**Authors:** Marianne Keith, Alba Park de la Torriente, Antonia Chalka, Adriana Vallejo-Trujillo, Sean P. McAteer, Gavin K. Paterson, Alison S. Low, David L. Gally

## Abstract

This study supports the development of predictive bacteriophage (phage) therapy: the concept of phage cocktail selection to treat a bacterial infection based on machine learning models (MLM). For this purpose, MLM were trained on thousands of measured interactions between a panel of phage and sequenced bacterial isolates. The concept was applied to *Escherichia coli* (*E. coli*) associated with urinary tract infections. This is an important common infection in humans and companion animals from which multi-drug resistant (MDR) bloodstream infections can originate. The global threat of MDR infection has reinvigorated international efforts into alternatives to antibiotics including phage therapy. *E. coli* exhibit extensive genome-level variation due to horizontal gene transfer via phage and plasmids. Associated with this, phage selection for *E. coli* is difficult as individual isolates can exhibit considerable variation in phage susceptibility due to differences in factors important to phage infection including phage receptor profiles and resistance mechanisms. The activity of 31 phage were measured on 314 isolates with growth curves in artificial urine. Random Forest models were built for each phage from bacterial genome features and the more generalist phage, acting on over 20% of the bacterial population, exhibited F1 scores of >0.6 and could be used to predict phage cocktails effective against previously untested strains. The study demonstrates the potential of predictive models which integrate bacterial genomics with phage activity datasets allowing their use on data derived from direct sequencing of clinical samples to inform rapid and effective phage therapy.

**Significance Statement:** With the growing challenge of antimicrobial resistance there is an urgency for alternative treatments for common bacterial diseases including urinary tract infections (UTIs). *Escherichia coli* is the main causative agent of UTIs in both humans and companion animals with multidrug resistant strains such as the globally disseminated ST131 becoming more common. Bacteriophage (phage) are natural predators of bacteria and potentially an alternative therapy. However, a major barrier for phage therapy is the specificity of phage on target bacteria and therefore difficulty efficiently selecting the appropriate phage. Here, we demonstrate a genomics driven approach using machine learning prediction models combined with phage activity clustering to select phage cocktails based only on the genome sequence of the infecting bacterial strain.

## Introduction

Uropathogenic *Escherichia coli* (UPEC) are among the most common causes of urinary tract infections (UTIs) in both humans and companion animals. It is reported that up to 80% of UTIs in humans and 35-69% of UTIs in small animal pets are caused by UPEC (1, 2). UTIs are widespread, affecting an estimated 150 million people a year worldwide (3) and 40% of women will develop an UTI during their lifetime (4). It is also reported that 14% of dogs will suffer with a UTI over their lifetime (5). UPEC employ several virulence factors that give them the capacity to colonise, invade and survive outside of the intestine (3). They can transfer from the intestinal environment and ascend and colonise the urinary tract often causing uncomplicated transient infections that selfresolve. However, the likelihood of antibiotic treatment increases if the infection persists and/or the symptoms are more severe. Bladder (cystitis) infections can sometimes ascend to the kidney(s) (pyelonephritis) and potentially into the bloodstream resulting in bacteraemia. Up to 50% of human sepsis cases are associated with *Escherichia coli* (*E. coli*) originating from a urinary tract infection (6, 7). Conversely and much less commonly, kidney infections and UTIs can result from a bloodstream infection. Recurrent and chronic infection increases the exposure of strains to multiple antibiotics and therefore the potential emergence of multidrug resistant bacteria (MDR) (3, 8, 9) and treatment failure. Trimethoprim-sulfamethoxazole, fluoroquinolones and cephalosporins are common first-line antibiotics used to treat UTIs in humans. Antibiotics from these classes are also used to treat pets with UTIs (5, 10). Antibiotic resistance is a growing problem globally and has serious consequences, including increased healthcare costs and the spread of potentially lifethreatening infections. High-risk globally disseminated MDR human UPEC strain types such as ST131 and ST410 are now being reported in veterinary settings (11). Alternative non-antibiotic treatments for UTIs are urgently required to help prevent further development of antibiotic-resistant bacteria and offer treatment options for MDR infections. A recent review highlights potential alternatives which include bacteriophage (phage) therapy but acknowledges further advances in the understanding of phage biology is required before phage therapy could be routinely used (12)

The use of phage, viruses that specifically kill bacteria, is now gaining traction as an alternative to antibiotics for the treatment of bacterial infections. Phage were first described over 100 years ago by Felix d’Herelle (13) and used to treat bacterial infections in this era (14, 15). However, the advent and routine use of antibiotics by the 1940s meant phage therapy development stalled globally. As a result of the rising threat of AMR, there has been a renaissance in phage research with major advances in the characterisation of bacterial resistance mechanisms (16, 17) and knowledge of the counter-offensive strategies evolved by phage (18, 19). Phage as a therapy have many advantages; they are naturally occurring, usually highly specific, non-toxic, effective on antibiotic resistant bacteria and self-dosing. However, the specificity is also a disadvantage as ideally a therapeutic phage needs to be matched to an infecting strain, traditionally by manual screening using agar overlay assays. Phage are sought that are ‘generalist’ meaning they are active on a reasonable proportion of an infecting species, but this is a much greater challenge for some species over others depending on their genome plasticity. This includes *E. coli*, as integrated prophage play a major role in their diversification and often introduce defence mechanisms against other phage to promote their own survival.

To overcome phage specificity issues and potential resistance development, phage therapy has often been developed using cocktails of phage, usually between 2 and 10 phage (20). While phage interference can occur especially if too many phage are included, such cocktails are more likely to include phage that can predate on the infecting strain and conceptually there is a reduced likelihood that the bacteria can develop resistance if the phage use different infection pathways (21–23). Even with cocktails, the incredible diversity of *E. coli* associated with UTIs in human and companion animals make phage selection for effective treatment a major challenge. Recent advances in genomics and machine learning analysis should allow the development of ‘smarter’, personalised medicine approaches within the field of phage therapy. Machine learning is a computational technique capable of analysing large and complex datasets (24) which has already been applied for predicting viral hosts (25–27). However, previous studies have focused on species-level host differentiation, whereas phage therapy will require a greater level of resolution for host attribution, down to individual bacterial isolates. To that end, we propose that machine learning models custom-built around predicting infection efficacy of phage on bacterial isolates of a species can be used to design personalised phage cocktails for patients, aiming to achieve improved efficacy over a generic cocktail.

To help address a knowledge gap of bacterial-phage interactions in clinically relevant conditions and move towards a predictive approach to phage therapy, we have generated a large data set of phage-UPEC interactions in an artificial urine medium. This has been used to train machine learning models for predictive phage therapy but also gives a wealth of information to inform the selection of phage, including activity groups and distribution of resistance mechanisms. Coupled to the direct sequencing of infected urine samples (28), the study shows how therapeutic phage cocktail selection can be quickly achieved using a bacterial genomics approach and appropriate interaction datasets. These cocktails can be constructed at a generic level, best on the shelf preparation or in a bespoke manner to an individual infection.

## Results

### Local epidemiology of UPEC strains for phage treatment

For this study, focused on urinary tract infections, 314 UPEC strains from both canine (n=203) and human (n=111) patients in the Edinburgh area were whole genome sequenced (Illumina, MicrobesNG). This was required as input data to train the machine learning models but also enabled an understanding of local strain diversity. In addition, it facilitated the selection of representative strains for phage enrichment to expand our phage collection from wastewater samples. Figure 1A shows the phylogenetic tree alongside their source host for the 314 strains as well as a set of 10 validation strains used later in the study. The sequences were uploaded to Enterobase allowing assessment of phylotype, sequence type (ST) and O-antigen. Some phage use components of the lipopolysaccharide (LPS) including the O-antigen portion as host receptors and other phage activity may cluster with phylogeny so mapping this information against phage activity could be important for rational cocktail design even without predictive models.

**Figure 1.**
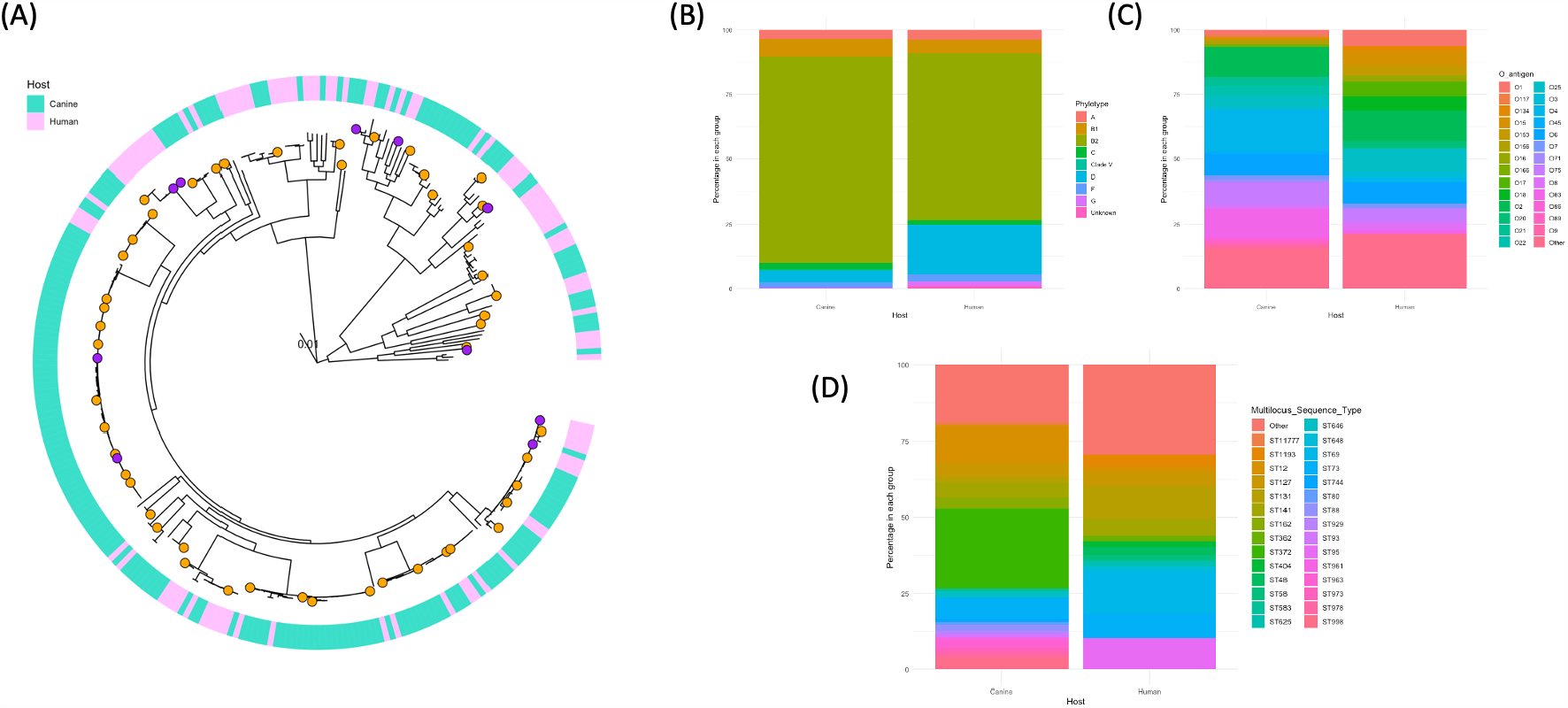
(A) Phylogentic tree showing the relationship of the UPEC strains from the collection used in this study (314 strains in training set and 10 strains from validation set). The host source (canine or human) is shown in the outer ring. Orange dots show the 50 representative strains used to test the general and bespoke cocktails and the purple dots show the 10 ‘unseen’ strains used to validate the machine learning models. (B, C, D) Charts showing the percentage of the canine and human strains (n=314) belonging to the different phylotypes (B), O-Antigen type (C) and sequence type (ST) as defined by Enterobase.

As expected, most of the *E. coli* strains in the UPEC collection belonged to phylotype B2 and D, the phylotypes most associated with extraintestinal disease in humans and domestic animals (Figure 1B) (29, 30). The human strains had a higher proportion associated with phylotype D than the canine strains (18.9% of strains compared with 4.4%). The differences in predominant ST and to a lesser extent O-antigen types between the human and the canine derived isolates (Figure 1C and 1D) indicate that different types of *E. coli* strains are associated with UTIs in these two hosts locally and this is likely to impact phage specificity. ST131, ST69, ST95 and ST73 are globally disseminated pathogenic *E. coli* lineages that are associated with UTI and bloodstream infections (31) and these are the most abundant ST groups represented in the human UPEC set. ST73 is also an abundant ST group in the canine strains (6.9%) although ST372 and ST12 represent almost 38% of the canine isolates.

### Generation of a host-phage interaction data set

To develop machine learning models and better understand genotype to phage activity phenotype, an interaction data set is required for training and analysis. While this potentially could be mined from online databases, the methods used would be varied and there would be a lack of negative data. To overcome this, and to test for activity in conditions more closely reflecting those *in vivo*, we generated an interaction dataset by growing 314 bacterial UPEC strains in an artificial urine (AU) medium in 96 well plate growth assays and challenging them with a set of 31 phage. Ultimately phage selected to treat UTIs need to be active in the host environment and to assess phage activity against UPEC the ideal growth medium would be urine but repeated urine collection would introduce too much variability. Our preliminary work for this study therefore included development of an AU medium based on published studies (*SI Appendix*, Figure S1A). The AU approximated canine pooled urine in terms of growth dynamics and phage activity and also demonstrated that the activity of certain phage could be lost when the bacteria were cultured in urine and AU compared to Lysogeny/Luria Broth (*SI Appendix*, Figure S1B).

Phage were initially selected from our phage library by screening of activity against a small subset of UPEC strains using LB plate assays. Around 60 phage were then assayed in liquid AU against 38 bacterial isolates selected to represent the diversity of the wider UPEC collection. From this data, a final set of 31 phage were selected to generate interaction data for the remaining 284 bacterial isolates (314 in total), >9000 interactions, each measured in triplicate against a no phage control. The final phage set were primarily selected as those showing the broadest host range and diverse activity profiles but some narrower range phage and phage with very related activity were also included. An interaction score was determined for each strain-phage combination using the ratio of the area under the curve (AUC) for phage treatment over a no-phage control culture (*SI Appendix*, Table S1). A score of 100 indicated no activity while scores under 100 were indicative of phage interaction and reduced bacterial growth. Some interactions scored higher than 100 in the interaction assays i.e. where the bacteria grew better in the presence of the phage preparation than the no phage control. The scores were plotted as a heatmap to visualize patterns of phage activity across the strain collection (Figure 2) with any interaction that scored over 100 being given the score of 100 for the purposes of the heatmap illustration. With a score of 60 or less indicating clear phage activity, only two phage in our test set showed activity on more than 40% of strains in the bacterial collection, Chap1 (54%) and Nea2 (41%) (Fig 2 & 3A). 4% of strains (n=14) were completely resistant to all 31 phage tested (all phage interactions scored higher than 90). Subsequent to the generation of this data set, these strains were used to enrich for further phage which will be used to broaden the host range of the phage collection.

**Figure 2.**
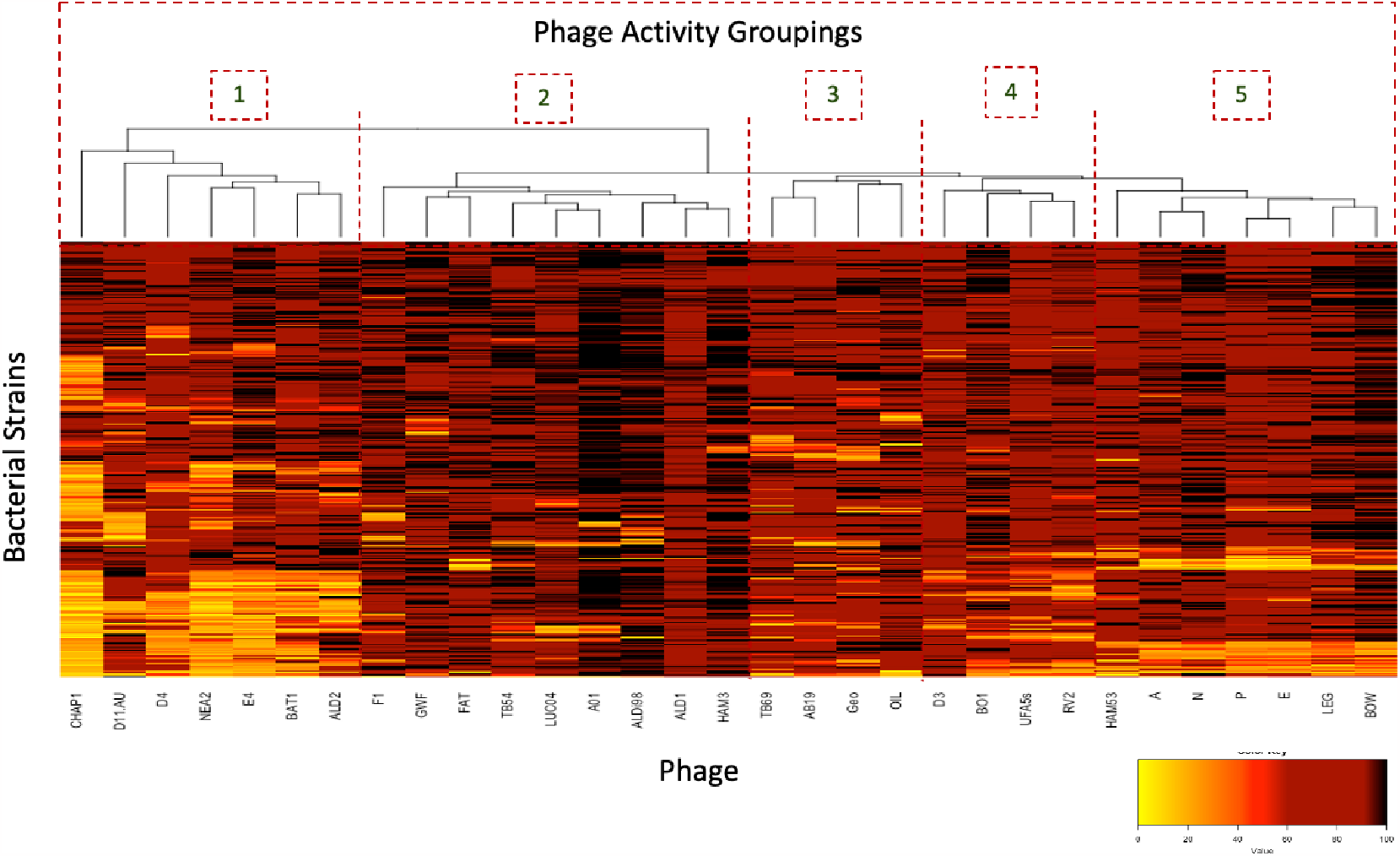
Heatmap of bacterial (n=314) phage (n=31) interactions based on score from growth assay in artificial urine. Low scores (yellow) represent high phage activity and good inhibition of bacterial growth. High scores (dark red) represent limited phage activity. Using heatmap2 in gplots (v3.1.3,), pairwise comparisons of the data interactions allowed the phage to be clustered into five primary activity groups to aid with cocktail selection.

### Machine learning predictions of phage activity

The principal aim of this work was to generate an interaction database that could be used, with features extracted from bacterial whole genome sequences, to train machine learning models to predict phage activity based on an *E. coli* sequence. This is a ‘proof of principal’ study with 31 phage to gain insight into the capacity of the method to predict phage activity, and design effective cocktails, without further manual screening.

The main genomic features used for testing were predicted genes generated by a pangenomic analysis and defined as protein variants (PVs). Separate models were created for each phage, resulting in a total of 31 models. Models were initially evaluated on RMSE scores, which represent the difference between the observed and predicted scores (available in *SI Appendix*, Table S2). The individual models (Figure 3B & *SI Appendix* Figure S3) had average RSME values between 8 and 23. The interaction datasets for some of the phage were very imbalanced (Figure 3A), which required a more stringent investigation of model bias. To reflect their use in a clinical setting, observed and predicted interaction scores were classified either side of a threshold score of 60 (below 60 was a positive (active) interaction and above 60 a negative (inactive) interaction). A confusion matrix was generated for each phage model, and the models were further evaluated on classifier-based metrics; F1 (Figure 3B), recall and precision (*SI Appendix*, Figure S2). F1 was chosen as the best performance metric, taking class imbalance into account, and there was a clear trend that more generalist phage yielded more reliable models (Figure 3B).

**Figure 3.**
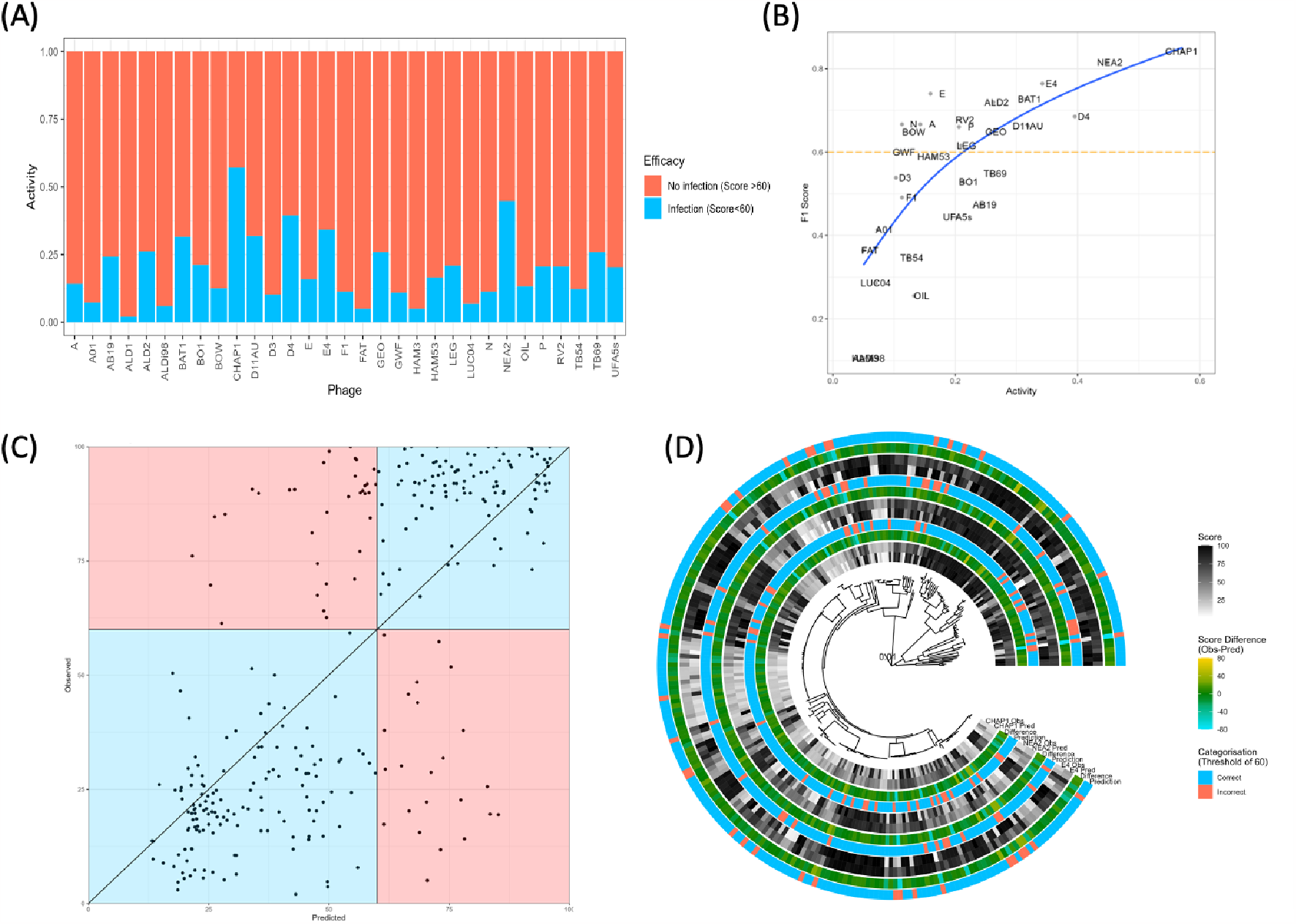
(A) The infectivity of the 31 phage across the *E. coli* dataset. Infections were determined based on a score of less than 60. (B) F1 scores vs activity (ratio of infected isolates) for PV phage models. The dashed line delineates a threshold of 0.6 for the F1 score as a metric for a ‘reliable’ model. (C) Scatter plot of the predicted vs observed interaction scores for phage CHAP1. The diagonal line represents a perfect prediction (observed=predicted). The chart is divided into 4 colour-coded quadrants to represent correct/incorrect predictions (True Negatives and True Positives as blue, False Negatives and False Positives as red). (D) Phylogenetic SNP tree of the isolates used for model training. Exemplar phage models for CHAP1, NEA2 and E4 are shown. For each model, the observed and predicted interaction scores are shown in black and white, followed by the difference between the two scores as a colour gradient, followed by whether each isolate would be categorised (score <60 the cut-off value for activity) in the same way by both the observed and predicted scores.

The spread of predicted vs observed scores for the most generalist phage Chap1 (F1=0.83) is shown in Figure 3C. Similar figures for the other 30 phage models can be found in the supplemental data (*SI Appendix*, Figure S3). The information was also plotted across the phylogeny of the strains, based on core SNP variation, and activity as well as significant positive or negative deviation of the predicted score from the measured score was generally distributed across the phylogeny and not limited to one sub-cluster (Figure 3D and *SI Appendix*, Figure S4).

### Assessment of different approaches to select phage cocktails

Clinically, phage are often administered as a cocktail of multiple phage. The advantage of using a phage cocktail is two-fold; (i) the construction of a general cocktail that could be used ‘off the shelf’ to treat an infection where the inclusion of a range of phage should increase the chance at least one is effective against the infection; (ii) a bespoke cocktail where the included phage are matched to the infection and so should all be effective and the inclusion of multiple active phage should help overcome the development of bacterial resistance to the phage treatment. In both these cases it is useful to include phage with diverse activity and therefore an increased probability they use different mechanisms to infect the host bacteria. Here, the phage were clustered by activity on the bacterial strains generating five primary activity groupings. For this, a matrix with the bacteriumphage interaction scores was used to produce a heatmap (gplots version 3.1.3) and dendrogram based on pairwise comparisons (Figure 2). Cocktails were generated based on using one phage from each activity group. We selected Chap1, F1, TB69, RV2 and Phage P for a general cocktail, which were the broadest range phage from each activity group (Figure 2 and 4A, *SI Appendix*, Table S4). From the dataset of single phage interactions, this cocktail was expected to be effective on 64% (201/314) of strains in the training data set as at least one phage in the cocktail has activity against the strain. This was validated on a subset of 50 strains from the UPEC collection, selected to represent the diversity across the collection (orange dots on Figure 1A). As expected, this general cocktail was effective on 64% of the strains tested (Figure 4A, *SI Apendix*, Table S5). With most of the strains tested, the general cocktail limited the strains to a similar level to the most active single phage in the cocktail on that strain. There was little evidence of synergistic effects from the general cocktail where the score for the cocktail was less than any of the individual phage. In fact, the opposite could be seen where the cocktail score was higher than the best individual phage indicating some antagonist activity.

**Figure 4.**
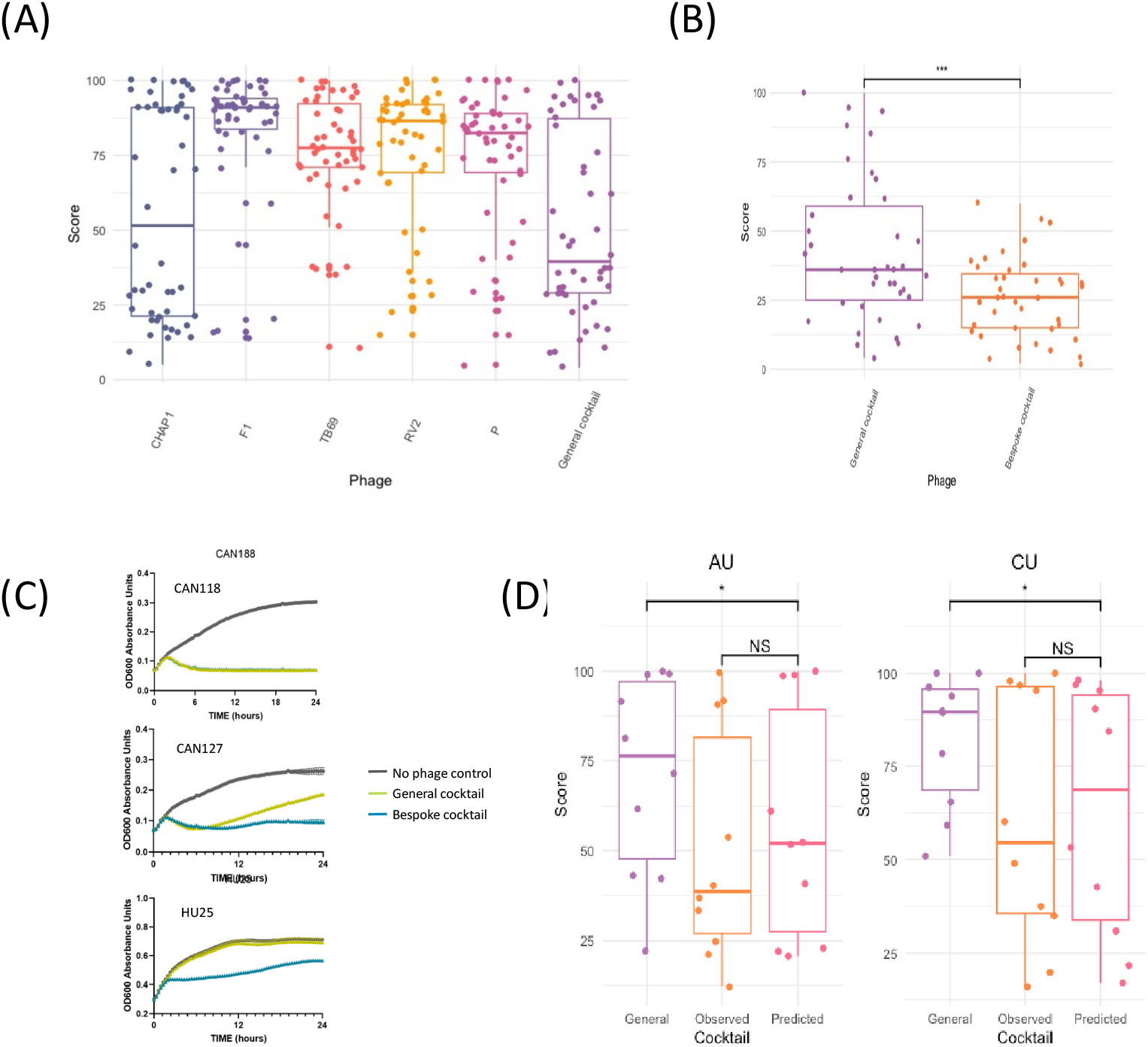
Phage cocktail analysis (A) Using a representative set of 50 UPEC strains from the training set of *E. coli* the General Cocktail was tested. The box plots show the distribution of scores for the individual phage and for the cocktail. (B) For the 38 strains (out of the set of 50) that had more than 2 effective phage (from different activity groups) a bespoke cocktail was created and scores recorded (*SI Appendix*, Table S4 & S5). A statistical analysis (paired t-test) comparing the general to the bespoke cocktail for these 38 strains shows that the bespoke cocktail is more effective (p= 2.081e-05). (C) Representative growth data that was used to generate the activity scores comparing the general and bespoke cocktails for three strains where there is no difference between the cocktails (CAN118), where there is bacterial resistance evolving for the general cocktail (CAN127) and where the bespoke cocktail is effective over the general cocktail (HU25). (D) A set of 10 strains that had not been in the training set were used to validate the prediction models. Using defined rules bespoke cocktails were created using the predicted scores and separately the observed scores and compared alongside the general cocktail. This was done in AU and canine urine (CU) and the scores plotted (representative data from one experiment, repeat data available in *SI Appendix*, Table S6). Analysis of these results (paired t-test) showed the predicted cocktail was more effective than the general cocktail (p=0.03502 (AU), p=0.03614 (CU)). There was not a statistical difference in effectiveness between the observed or predicted cocktail (p=0.1746 (AU), p=0.4288 (CU)).

Bespoke cocktails could also be designed using the phage interaction dataset by choosing a selection of diverse phage active against a strain, a feature that would help combat the development of bacterial phage resistance. Only a small proportion of strains from the test set of 50 were susceptible to more than two phage in the general cocktail (5 strains, *SI Appendix*, Table S5). A bespoke cocktail selected for each strain would mean that all phage included in the cocktail should some activity against the strain. A few strains from the set of 50 could not have a bespoke cocktail of 5 phage designed due to a limitation of overall phage activity, but assessment was possible for 38 strains. The bespoke cocktails were significantly more effective when compared to the general cocktail with a lower median score (Figure 4B, p=2.081e-05). As expected, analysis of the growth curves from these interactions generally showed much less bacterial regrowth in the bespoke cocktails compared to the general cocktail (example shown in Figure 4C) resulting in lower scores.

A key question is whether the predictive models have the potential to inform phage selection of cocktails and how selection using this method compares to the general cocktail. To assess this and to look at phage activity on strains outside the original 314 in the training dataset, 10 newly isolated clinical UPEC strains (3 canine, 7 human), were genome sequenced and these sequences run through the predictive models to give predicted phage interaction scores (*SI Appendix*, Table S6). From the predicted scores, cocktails were constructed using specific rules (materials and methods) so they could be compared to cocktails created using the same rules from observed phage activity data (that was subsequently carried out for validation). Phage cocktail composition for each strain can be found in supplementary material with selection accounting for F1 scores (*SI Appendix*, Table S6). The full set of cocktails (general, bespoke observed and bespoke predicted) were tested against the strains in AU, and as a final validation, in canine urine (CU). The scores from each of these cocktails for duplicate experiments are recorded in supplementary data (*SI Appendix*, Table S6) and the distribution of scores shown in Figure 4D. Three of the strains were insensitive to all the phage tested and none of the cocktails worked well on these strains. The bespoke predicted cocktail performed at a level equivalent to the observed bespoke cocktail and was significantly more effective than the general cocktail in both AU and canine urine (Figure 4D).

## Discussion

This history of successful phage therapy over the last century has often occurred when a more bespoke approach has been used, i.e. where phage are identified that act on the infecting bacteria. While there are some exceptions to this, including ‘jumbo’ phage targeting a wide range of *Salmonella* serovars for a recent advance in food safety (32), the high-profile news stories and the long-standing work at the Eliava Institute are based on identifying the infecting organism and selecting appropriate phage from extensive phage collections. This highlights the potential of phage therapy but a bespoke approach is much harder to do at scale. The selection of appropriate antibiotics is strengthened with knowledge of the antibiotic resistances of the infecting strain, and similarly a diagnostic platform that provides information about phage effectiveness would greatly advance the development of phage treatment. The premise for this study was that genomic information about the infecting strain should be the basis of phage cocktail selection. Towards this, phage activity models were generated from bacterial sequence data combined with phage interaction data allowing prediction of phage that are active on ‘unseen’ strains, i.e. those not used in the model training sets. Phage predicted as active from the models can then be combined into cocktails for treatment.

*E. coli* UTIs are an appropriate challenge for phage selection as horizontal gene transfer mediated by phage and plasmids has resulted in an ever-increasing pangenome of >100,000 genes relative to a ‘core’ genome of only a few hundred genes. Prophage integration also introduces resistance mechanisms to phage infection and this accounts for major variation in phage susceptibility along with receptor and metabolic diversity. As a consequence of the presence of these genomic determinants, phage activity on a strain should be identifiable from its genome sequence. To date, most of the predictive effort has been focused on possible hosts for phage based on phage genome sequencing, reviewed in (24), in particular prediction of bacteriophage sequences from metagenomic studies and predictive calling of possible bacterial hosts. These have mainly used nucleotide information including k-mer biases, although one study has taken into account CRISPR sequences (33). These studies are identifying phage interactions at the level of phylum or genus and have not been focused on defining phage that would be active on clinically infecting strains of a species based on their sequence data. The idea of predictive models based on the different steps in infection has been proposed (34) and our approach presented here, while relatively agnostic in that it uses differential PVs (predicted genes) as features, may well include receptors and resistance mechanisms. However, this is hard to dissect as the majority of features used are grouped genes with no clear annotation. Future pipelines for phage selection could apply both associative features as well as those based on functional knowledge. Either way, good quality training sets with phenotypic interaction data will still be critical.

For the majority of methods using genome features for prediction (reviewed for source attribution in (35)) there will be significant phylogenetic influence. This will mean that features are used with statistically significant associations with phage activity but which do not have a functional relationship to phage susceptibility. This is not an issue for the models as long as they are used within a similar population structure to the training dataset but there would be reduced accuracy if the models are applied to strains from different structures. Machine learning models are also prone to replicating existing biases in their training datasets. This was demonstrated in the predictions scored of phage with very imbalanced datasets (e.g. A01), in which the model was trained on mainly high interaction scores indicating no infection and a very small subset of low scores, indicating infection. The resulting model struggled to differentiate isolates that the phage would actually infect and would be prone to generating false negatives. From a cocktail perspective, the failure to select a phage that would have been active (false negative) is actually preferred to false positives, although a high rate of false negatives may lead to a paucity of phage predicted as active from which to select a cocktail. We note that with the better models presented in this study the frequency of false positives and false negatives falls within a range where if five phage are selected for a cocktail the majority will be correctly predicted. For example, for Chap1 the predictive model generated 7% false negative scores (22/314) and 10.8% were false positive scores (34/314).

Our analysis on a set of completely independent sequenced isolates did demonstrate that the predicted cocktails were more effective than a generic cocktail and in fact were comparable with the bespoke cocktails that were based on individual measured activity. This result on ‘unseen’ isolates provides confidence in the main concept of the study. To really move such predictive phage therapy into the clinic, there are areas the research has highlighted that require further development along with the safety and registration issues that are being tackled more globally by the phage community. A key improvement would be inclusion of more generalist phage in the interaction dataset. Based on the findings illustrated in Figure 3, the models trained on datasets with a class imbalance as high as 85:15 were able to produce results of practical value for phage selection, as defined by a predefined F1 cutoff of 0.6. For greater reliability, it would be preferable to use phage with a class imbalance less than this, and therefore phage that can target at least 30% of the population. More generalist phage will also provide more options for cocktail selection especially if they can be distributed across different activity groups. The work shown is a ‘proof of concept’ and valuable interaction datasets for prediction models could now be produced in a short period of time using high throughput platforms, including those that can measure 50 x 96 well plates with appropriate phage and strain collections in place.

Another major consideration is the actual activity of the selected phage in the infection environment, in this case urine which can vary considerably in terms of composition and constituent concentrations, especially pH and osmolarity, as well as factors associated with the host response to infection (36–38). We have tried to mitigate this by measuring phage activity in artificial urine but acknowledge this cannot capture the complexity of real urine as although it is a defined reproducible medium and a closer match to urine than many rich broths that are routinely used for phage activity studies that allow faster bacterial growth. The bespoke phage cocktails formulated in this study based on the predictive models showed good activity in canine urine, comparable with results in artificial urine (Fig. 4D), and this mirrored our preliminary work on the activity of a subset of individual phage in artificial urine vs pooled canine urine. (*SI Appendix*, Figure S1).

Predictive phage therapy would be used in conjunction with diagnostic sequencing. Direct sequencing of nucleic acids extracted from clinical sample has been successfully used for diagnosis for viruses, fungi and bacteria (39–41) including UTI (42, 43) and we have applied it to direct sequencing of *E. coli* in canine urine samples with respect to the selection of appropriate antimicrobials (28). There is the challenge of feature extraction for the phage models from long read data of more complex samples compared to assemblies of short reads from purified strains, as used in this study. If there are multiple strains associated with the infection then the models may still be useful to select phage, even without their separation at a genome level, although that remains to be tested. The testing of phage activity with sequenced clinical samples would be used to continually reinforce the learning dataset and improve predictions.

It should be emphasised that the primary aim of this study was to signpost what could be possible for phage therapy with machine learning or alternative predictive methods based on comprehensive interaction datasets. The study looked at both canine and human UTI *E. coli* isolates presented by patients at local hospitals and used a selection of 31 phage from an initial pool of just over 100. While these are only representative subpopulations, we consider they allowed the concept of predictive phage therapy to be tested with a promiscuous and important pathogen than can exhibit multi-drug resistance. We are optimistic that there will be a rapid expansion of advances in this space and the best methods could be combined in pipelines to transition the best approaches into clinical practice. However, the wider adoption of phage therapy in the veterinary or human clinic will require changes in licensing and management of expectations of both clinicians and the general public (44).

## Materials and Methods

### Strain Information

203 *E. coli* were isolated from UTIs in canine patients at the Royal (Dick) School of Veterinary Studies, Hospital for Small Animals between Jul. 2017 and Jun. 2019. In addition, 81 *E. coli* were isolated from urine and 30 *E. coli* from blood from patients with blood stream infections processed at The Royal Infirmary of Edinburgh (collected between Jan. 2019 and Nov. 2019). The canine, human urine and human blood isolates were designated “CANxx”, “HUxx” and “HBxx” respectively. Isolates were stored at -70 °C in 20% glycerol (G5516, Sigma, Germany) and streaked out to obtain pure, individual colonies on LB agar. A final set of 10 strains were collected from the same clinics towards the end of the study for validation of the prediction models. (canine, n=3, collected as part of a different study(28) and human, n=7, from The Royal Infirmary of Edinburgh collected Spring 2023).

### Phage Isolation and Propagation

Phage were propagated by growing the appropriate host strain to an OD_600_ 0.2 to 0.3 in LB or AU in a glass conical flask at 37ºC with shaking at 170 rpm before adding 500 μL of phage plaque filtrate in SM (50mM Tris-HCl pH7. 5, 100mM NaCl, 8mM MgSO4, 0.01% Gelatin (v/w)) buffer and continuing to incubate overnight. Infected cultures were centrifuged at 5,000 rpm for 10 minutes in a benchtop centrifuge to pellet the bacteria, and the phage lysate passed through a 0.22 μM syringe filter before long term storage at 4ºC. Phage were titred using double layer agar overlay plates and diluted to 1 x 10^6^ per mL working stock concentration using SM buffer.

### Growth media and Culture Conditions

1L volumes of artificial urine (AU) were prepared (according to the following protocol, https://dx.doi.org/10.17504/protocols.io.kxygx3exzg8j/v1), aliquoted into single use tubes, and frozen until required. Each aliquot was filter sterilised through a 0.22 μM syringe filter prior to use. Canine urine was collected from healthy dogs, filter sterilised using a 0.22 μM syringe filter, aliquoted and frozen until use.

### Interaction Assays including AUC analysis

Phage interactions assays were performed using 31 phage against 314 *E. coli* isolates using an MOI of 0.01 (∼1 phage per 100 bacteria). This MOI ratio was chosen to ensure that complete virus lifecycle infections were analyzed rather than abortive infections which can occur at higher MOIs. A single colony of an *E. coli* isolate was picked to inoculate 5 mL LB (LBL0102, Formedium Ltd, England) and incubated overnight at 37°C with shaking at 170 rpm. 50 μL of overnight culture was subcultured into 5 mL fresh LB and grown at 37ºC and 170 rpm shaking to an optical density (OD_600_) of 1.0 (equivalent to 10^8^ bacterial cells per mL). During this incubation, 10 μL of SM buffer (for no phage controls) or 10 μL phage at 1 x 10^6^ per mL (phage stocks were diluted in SM buffer) was added to a flat-bottomed 96-well microplate. 180 μL of AU was added to all wells. 10 μL of *E. coli* at OD_600_ of 1 was added to all wells resulting in a final reaction volume of 200 μL. A gas permeable, sterile, optically clear plate seal was applied to the plate (4ti-0516/96, Azenta Life Sciences) to control for evaporation and prevent phage contamination within the plate reader. Microplates were run on 1 of 3 identical Multiskan FC photometers (51119100, Thermo, China) with absorbance measurements at 620 nm taken every 20 minutes immediately after a 5 sec mix over a time course of 18 hours and constant incubation at 37ºC.

All interactions and controls were measured with at least three technical repeats and additional controls on each plate for growth and sterility, Media controls comprised 180 μL AU, 10 μL SM buffer, and 10 μL LB in place of bacterial culture. While the number of interactions involved were too large to carry out each interaction as triplicate biological repeats, biological repeats of a limited number of strains were carried out periodically to ensure repeatability of the assay over time and consistency of phage activity. Based on 31 phage measured on 314 bacterial strains with the technical repeats, a total of >27,000 growth curves were produced and analysed. The Multiskan FC photometers use SkanIt Software (Thermo) allowed raw data to be exported as a Microsoft Excel file. Phage interaction scores were calculated by comparing the average area under the curve (AUC) of the plus phage triplicate wells against the average AUC for the no phage control wells using PRISM software (GraphPad). (Score = (AUC plus phage/AUC no phage control) x 100). For the purposes of plotting data where the score generated was over 100 it has was recorded as a value of 100 representing no phage activity.

### Genome Sequencing

Genome sequencing of isolates was provided by MicrobesNG (http://www.microbesng.com) using paired Illumina reads. The reads were uploaded to Enterobase and assembled using the Enterobase Tool Kit (https://github.com/zheminzhou/EToKi). The assemblies were then annotated with prokka version 1.14.5 (45) Enterobase’s Tool Kit was also used to generate a maximum likelihood phylogenetic tree based on core SNP.

### Machine Learning and prediction pipelines

After filtering for fragmented assemblies (<600 contigs), 301 of the 314 isolates were used for model training and testing. The main features used for testing were predicted genes, defined as predicted protein variants (PVs) generated from a pangenome analysis of the 301 bacterial sequences using panaroo version 1.2.9 (46). PVs were encoded in a binary format (1/0) to indicate presence/absence in an isolate.

Random forest regression models were created for each of the 31 phage using the MUVR package (47) using 1 75-25 split of training/testing data and 10-fold cross validation. The predicted value was the interaction score calculated during the interaction assays analysis. (Score). Model performance was evaluated using the Root Mean Square Error (RMSE) statistic, which is calculated by taking the square root of the average of the squared differences between the predicted and actual values. RMSE provides a single numerical value that represents the overall accuracy of a prediction model, with lower values indicating better predictive performance. F1, recall, precision and related statistics were created by classifying each observed and predicted phage interaction based on the score (Score<60: infection; Score>60 No infection).

Feature reduction was performed recursively using the MUVR package. For each phage, an initial model using all PVs was created. Then the top-ranking features of a model were selected to create a new model with a reduced number of features and the RMSE values between the old and the new model were compared. If the models had a similar RMSE, the top-ranking features of the new model were used to create another new model, and the process repeated.

### Phage cocktail testing

Cocktail interaction assays were set up in a similar manner to the single phage assays described above. Phage were selected for the bespoke cocktails (both predicted and observed) using the same design rules; (i) to include the lowest scoring (highest activity) phage from each interaction grouping (Figure 2) as long as it scored under 80 (ii) if no active phage available from a group then additional phage(s) were added that were the second lowest scoring phage from activity groupings starting with the remaining phage with the lowest score until 5 phage in the cocktail (iii) if less than 5 phage available with scores less than 80 then less phage were used in the cocktail. Working stocks of phage cocktails were prepared by mixing equal volumes of each selected phage already prepared at 10^6^ per mL such that the 10 μL volume transferred to each triplicate well of a 96-well plate contained an equivalent total number of virus particles to the single phage assays, so that the overall MOI remained at 0.01. Additionally, the cocktail assays were allowed to proceed for 24 hr.

### Genomic Data

The genomic data generated for this work has been deposited in Enterobase (https://enterobase.warwick.ac.uk/species/ecoli/search_strains?query=workspace:97040).

## Acknowledgments

We are very grateful to Jennifer Harris in the veterinary microbiology laboratory at R(D)SVS for assistance in obtaining canine UPEC strains and Kate Templeton and Martin McHugh at Dept. of Medical Microbiology at the Royal Infirmary, Edinburgh for sorting approvals for human urine and blood *E. coli* isolates. The work was supported by a Canine Welfare Grant from The Dog’s Trust awarded to D.G. (*‘*Advanced phage therapy for multidrug resistant *E. coli* associated with canine urinary tract infections’), a PhD studentship from the National Council of Science and Technology of Mexico (CONACYT) alongside a joint studentship from the University of Edinburgh and the University of Glasgow awarded to A.P.T. and a Roslin Institute PhD studentship supported by the BBSRC awarded to A.C.

## References

1. I. U. Mysorekar, S. J. Hultgren, Mechanisms of uropathogenic Escherichia coli persistence and eradication from the urinary tract. Proc. Natl. Acad. Sci. 103, 14170–14175 (2006).

2. N. Nittayasut, et al., Multiple and High-Risk Clones of Extended-Spectrum CephalosporinResistant and blaNDM-5-Harbouring Uropathogenic Escherichia coli from Cats and Dogs in Thailand. Antibiotics 10, 1374 (2021).

3. A. L. Flores-Mireles, J. N. Walker, M. Caparon, S. J. Hultgren, Urinary tract infections: epidemiology, mechanisms of infection and treatment options. Nat. Rev. Microbiol. 13, 269–284 (2015).

4. R. Kaur, R. Kaur, Symptoms, risk factors, diagnosis and treatment of urinary tract infections. Postgrad. Med. J. 97, 803–812 (2021).

5. J. K. Byron, Urinary Tract Infection. Vet. Clin. North Am. Small Anim. Pract. 49, 211–221 (2019).

6. A. Sabih, S. W. Leslie, “Complicated Urinary Tract Infections” in StatPearls, (StatPearls Publishing, 2023) (July 15, 2023).

7. K. M. Hatfield, et al., Assessing Variability in Hospital-Level Mortality among U.S. Medicare Beneficiaries with Hospitalizations for Severe Sepsis and Septic Shock. Crit. Care Med. 46, 1753–1760 (2018).

8. F. Wagenlehner, et al., The Global Prevalence of Infections in Urology Study: A Long-Term, Worldwide Surveillance Study on Urological Infections. Pathogens 5, 10 (2016).

9. B. Kot, Antibiotic Resistance Among Uropathogenic Escherichia coli. Pol. J. Microbiol. 68, 403–415 (2019).

10. T. m. Sørensen, et al., Effects of Diagnostic Work-Up on Medical Decision-Making for Canine Urinary Tract Infection: An Observational Study in Danish Small Animal Practices. J. Vet. Intern. Med. 32, 743–751 (2018).

11. A. L. Zogg, K. Zurfluh, S. Schmitt, M. Nüesch-Inderbinen, R. Stephan, Antimicrobial resistance, multilocus sequence types and virulence profiles of ESBL producing and nonESBL producing uropathogenic Escherichia coli isolated from cats and dogs in Switzerland. Vet. Microbiol. 216, 79–84 (2018).

12. A. L. Zogg, Management of uncomplicated UTI in the post-antibiotic era (II): select non-antibiotic approaches - ScienceDirect (July 15, 2023).

13. A. L. Zogg, d’Herelle: Sur un microbe invisible antagoniste des… - Google Scholar (July 15, 2023).

14. F. d’Herelle, Bacteriophage as a Treatment in Acute Medical and Surgical Infections. Bull. N. Y. Acad. Med. 7, 329–348 (1931).

15. F. W. Twort, Further Investigations on the Nature of Ultra-Microscopic Viruses and their Cultivation. J. Hyg. (Lond.) 36, 204–235 (1936).

16. A. Bernheim, R. Sorek, The pan-immune system of bacteria: antiviral defence as a community resource. Nat. Rev. Microbiol. 18, 113–119 (2020).

17. W. P. J. Smith, B. R. Wucher, C. D. Nadell, K. R. Foster, Bacterial defences: mechanisms, evolution and antimicrobial resistance. Nat. Rev. Microbiol. 21, 519–534 (2023).

18. J. S. Athukoralage, M. F. White, Cyclic Nucleotide Signaling in Phage Defense and CounterDefense. Annu. Rev. Virol. 9, 451–468 (2022).

19. H. G. Hampton, B. N. J. Watson, P. C. Fineran, The arms race between bacteria and their phage foes. Nature 577, 327–336 (2020).

20. H. G. Hampton, Phage cocktails and the future of phage therapy | Future Microbiology (July 15, 2023).

21. F. L. Gordillo Altamirano, J. J. Barr, Unlocking the next generation of phage therapy: the key is in the receptors. Curr. Opin. Biotechnol. 68, 115–123 (2021).

22. C. Lood, P.-J. Haas, V. van Noort, R. Lavigne, Shopping for phages? Unpacking design rules for therapeutic phage cocktails. Curr. Opin. Virol. 52, 236–243 (2022).

23. M. Merabishvili, J.-P. Pirnay, D. De Vos, “Guidelines to Compose an Ideal Bacteriophage Cocktail” in Bacteriophage Therapy: From Lab to Clinical Practice, Methods in Molecular Biology., J. Azeredo, S. Sillankorva, Eds. (Springer, 2018), pp. 99–110.

24. Y. Nami, N. Imeni, B. Panahi, Application of machine learning in bacteriophage research. BMC Microbiol. 21, 193 (2021).

25. R. Aguas, N. M. Ferguson, Feature selection methods for identifying genetic determinants of host species in RNA viruses. PLoS Comput. Biol. 9, e1003254 (2013).

26. Q. Tang, et al., Inferring the hosts of coronavirus using dual statistical models based on nucleotide composition. Sci. Rep. 5, 17155 (2015).

27. M. Wardeh, M. S. C. Blagrove, K. J. Sharkey, M. Baylis, Divide-and-conquer: machinelearning integrates mammalian and viral traits with network features to predict virus-mammal associations. Nat. Commun. 12, 3954 (2021).

28. N. Ring, et al., Rapid metagenomic sequencing for diagnosis and antimicrobial sensitivity prediction of canine bacterial infections. Microb. Genomics 9 (2023).

29. N. Ring, The Clermont Escherichia coli phylo-typing method revisited: improvement of specificity and detection of new phylo-groups - PubMed (July 15, 2023).

30. J. R. Johnson, T. A. Russo, Extraintestinal pathogenic Escherichia coli: “the other bad E coli.” J. Lab. Clin. Med. 139, 155–162 (2002).

31. L. W. Riley, Pandemic lineages of extraintestinal pathogenic Escherichia coli. Clin. Microbiol. Infect. 20, 380–390 (2014).

32. A. M. Thanki, et al., A bacteriophage cocktail delivered in feed significantly reduced Salmonella colonization in challenged broiler chickens. Emerg. Microbes Infect. 12, 2217947.

33. W. Wang, et al., A network-based integrated framework for predicting virus-prokaryote interactions. NAR Genomics Bioinforma. 2, lqaa044 (2020).

34. C. Lood, et al., Digital phagograms: predicting phage infectivity through a multilayer machine learning approach. Curr. Opin. Virol. 52, 174–181 (2022).

35. N. Lupolova, S. J. Lycett, D. L. Gally, A guide to machine learning for bacterial host attribution using genome sequence data. Microb. Genomics 5, e000317 (2019).

36. C. Robinson-Cohen, et al., Estimation of 24-hour urine phosphate excretion from spot urine collection: development of a predictive equation. J. Ren. Nutr. Off. J. Counc. Ren. Nutr. Natl. Kidney Found. 24, 194–199 (2014).

37. D. B. Barr, et al., Urinary creatinine concentrations in the U.S. population: implications for urinary biologic monitoring measurements. Environ. Health Perspect. 113, 192–200 (2005).

38. E. N. Taylor, G. C. Curhan, Differences in 24-Hour Urine Composition between Black and White Women. J. Am. Soc. Nephrol. 18, 654 (2007).

39. K. Lewandowski, et al., Metagenomic Nanopore Sequencing of Influenza Virus Direct from Clinical Respiratory Samples. J. Clin. Microbiol. 58, e00963–19 (2019).

40. A. Ohta, K. Nishi, K. Hirota, Y. Matsuo, Using nanopore sequencing to identify fungi from clinical samples with high phylogenetic resolution. Sci. Rep. 13, 9785 (2023).

41. K. Nilgiriwala, et al., Genomic Sequencing from Sputum for Tuberculosis Disease Diagnosis, Lineage Determination, and Drug Susceptibility Prediction. J. Clin. Microbiol. 61, e0157822 (2023).

42. K. Schmidt, et al., Identification of bacterial pathogens and antimicrobial resistance directly from clinical urines by nanopore-based metagenomic sequencing. J. Antimicrob. Chemother. 72, 104–114 (2017).

43. L. Zhang, et al., Rapid Detection of Bacterial Pathogens and Antimicrobial Resistance Genes in Clinical Urine Samples With Urinary Tract Infection by Metagenomic Nanopore Sequencing. Front. Microbiol. 13, 858777 (2022).

44. J. D. Jones, H. J. Stacey, A. Brailey, M. Suleman, R. J. Langley, Managing Patient and Clinician Expectations of Phage Therapy in the United Kingdom. Antibiot. Basel Switz. 12, 502 (2023).

45. T. Seemann, Prokka: rapid prokaryotic genome annotation. Bioinforma. Oxf. Engl. 30, 2068–2069 (2014).

46. G. Tonkin-Hill, et al., Producing polished prokaryotic pangenomes with the Panaroo pipeline. Genome Biol. 21, 180 (2020).

47. L. Shi, J. A. Westerhuis, J. Rosén, R. Landberg, C. Brunius, Variable selection and validation in multivariate modelling. Bioinformatics 35, 972–980 (2019).

