## Supplementary Information Appendix for "Predictive phage therapy for *Escherichia coli* urinary tract infections: cocktail selection for therapy based on machine learning models"

This PDF includes:

Figures S1-S4
Tables S1-S7

Supplementary references

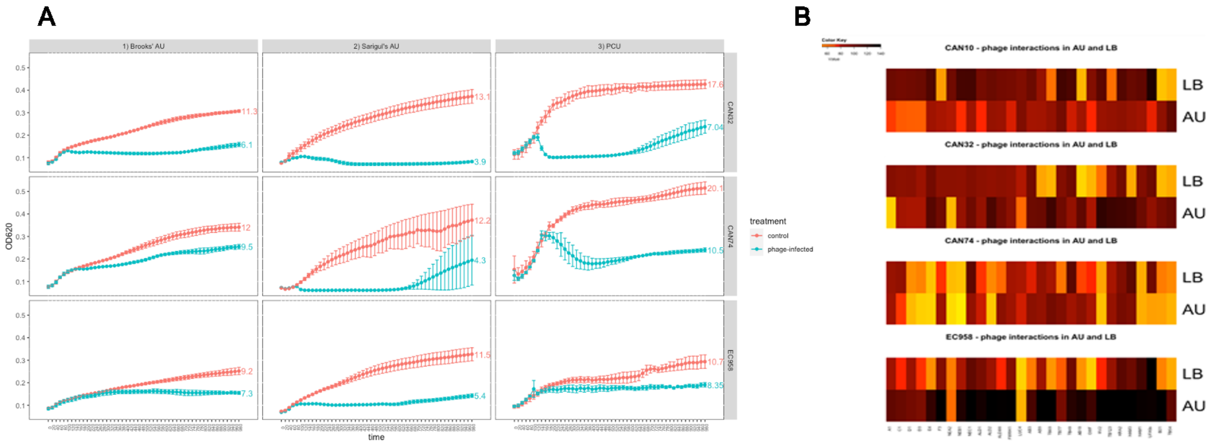

**Figure S1: (A)** To assess phage activity against UPEC the ideal growth media would be urine but for the number of interactions to be done in this study (n>10,000) the volume of pooled urine required was too great. To identify the optimum artificial urine (AU) for this work two different AU media recipes were first adapted from Brooks and Keevil(1) and Sarigul *et al.,(2)* then validated as representative media by comparing with phage activity against three clinical UPEC strains (two canine, one human) in pooled canine urine (PCU). A phage activity score was calculated using AUC (Area Under the Curve) of the non-phage infected and phage infected growth curves. Although growth was similar between the two AU medias the strains generally performed better in Sarigul's AU and the lysing effect of the phage was more evident in this media and compared well to the pooled canine urine. Interestingly the two canine isolated UPECs (CAN32 and CAN74) performed better in the pooled canine urine compared with the human isolated UPEC (EC958). All three strains grew well in adapted Sarigul's AU and phage activity reflected the PCU and therefore this was the media chosen for generation of the bacteria-phage interaction dataset. **(B)** Pairwise comparison of four UPEC interactions with 30 phage in LB and AU. The colour represents the score of the interaction: light yellow indicates a low score and therefore a strong inhibition of bacterial growth, whereas dark red represents high scores and no effect on bacterial growth. We then compared a subset of 30 phage against 4 bacterial strains in LB and our chosen AU media which confirmed the importance of using AU to generate our interaction data set as phage activity was often lost (and sometimes gained) in the AU media presumably indicative of these phage being less (or more) likely to be effective *in vivo*.

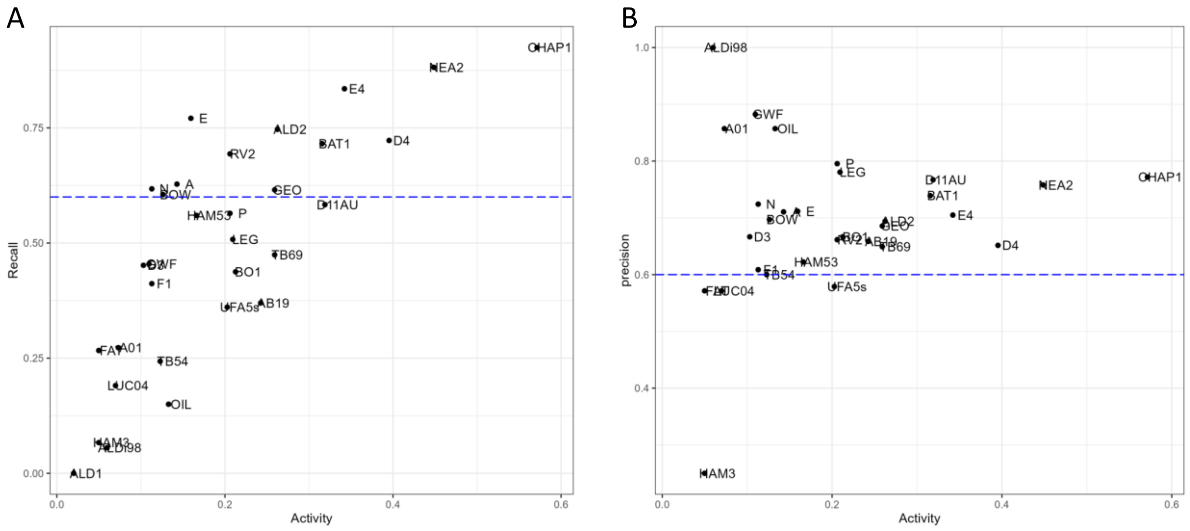

**Figure S2:** Recall (A) and precision (B) scores vs phage activity for the PV phage models. The recall scores show the proportion of actual positives that were identified correctly and the precision scores show the proportion of positive identifications that were actually correct. The F1 scores for the models (a combination of precision and recall) is shown in the main manuscript (Figure 3B).

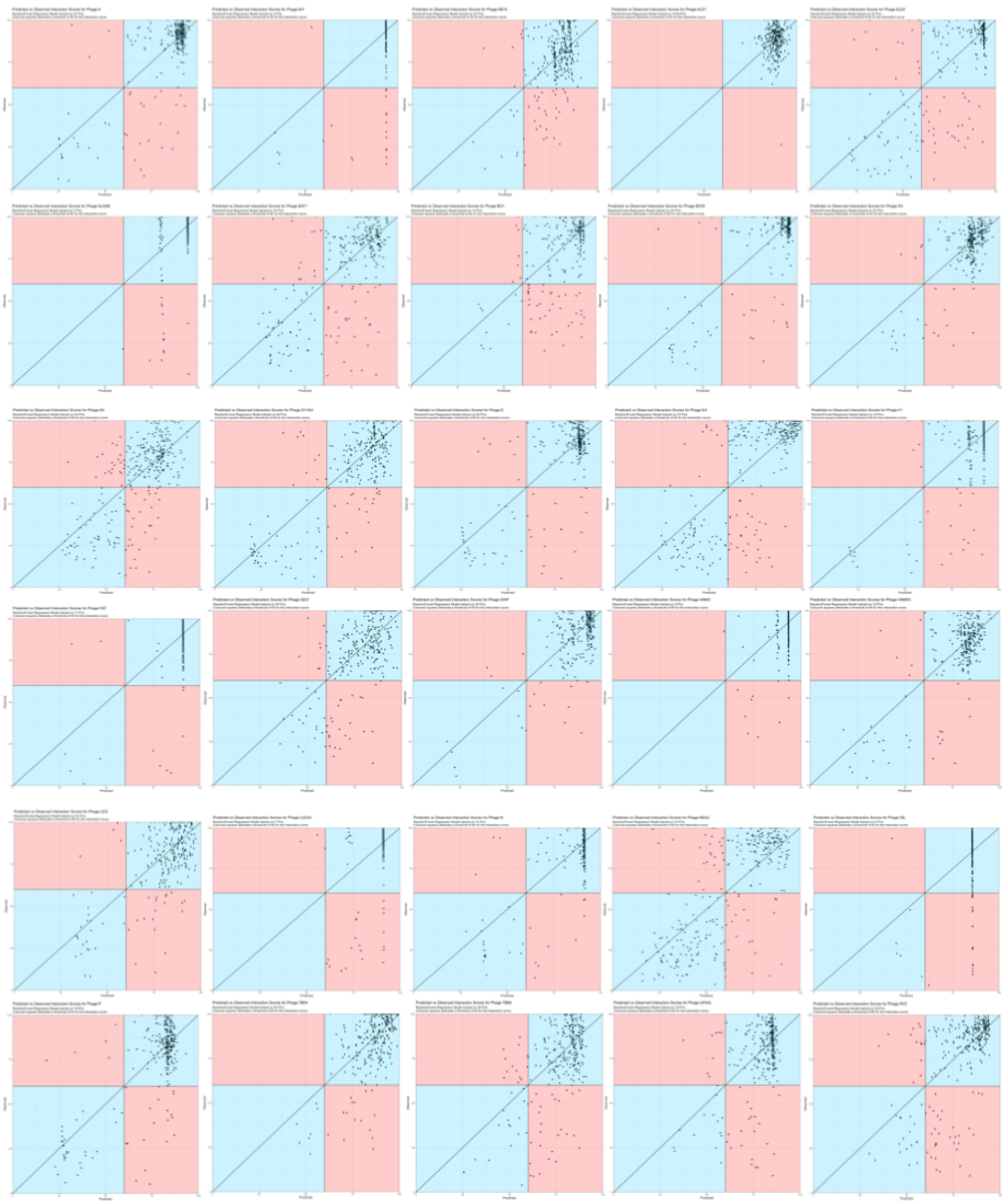

**Figure S3**: Scatter plots of the predicted vs observed interaction scores for the phage set (except CHAP1 which is shown in Figure 3C). The diagonal line represents a perfect prediction (observed=predicted). The chart is divided into 4 colour-coded quadrants to represent correct/incorrect predictions (True Negatives and True Positives as blue, False Negatives and False Positives as red). A score cut off 60 is used to define positive (less than 60) and negative (greater than 60).

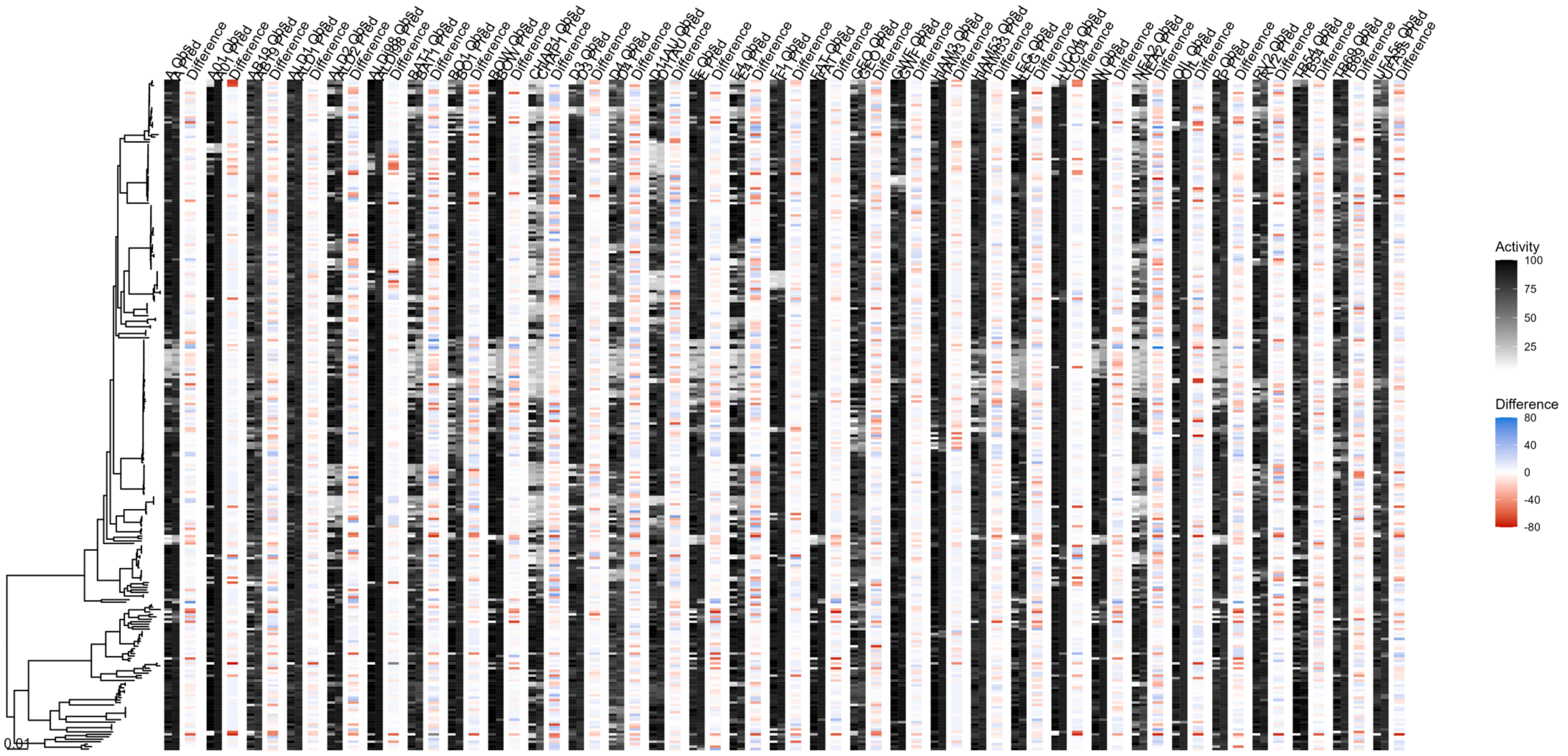

**Figure S4:** Phylogenetic SNP tree of the isolates used for model training. For each model, the observed and predicted interaction scores are shown in black and white, followed by the difference between the two scores as a blue/red gradient.

**Table S1:** The interaction scores for each strain-phage combination using the ratio of the area under the curve (AUC) for phage treatment over a no-phage control culture (Score = AUC plus phage/AUC no phage control) x 100).

| Isolate | CHAP1 | NEA2 | BAT1 | BO1 | N | Geo | ALD2 | OIL | D11 AU | A | ALD1 | D3 | E4 | F1 | HAM53 | LUC04 | RV2 | UFA5s | E | P | D4 | ALDi98 | GWF | AB19 | A01 | HAM3 | TB54 | TB69 | BOW | FAT | LEG |
|---|---|---|---|---|---|---|---|---|---|---|---|---|---|---|---|---|---|---|---|---|---|---|---|---|---|---|---|---|---|---|---|
| **CAN01** | 98 | 97 | 109 | 107 | 106 | 108 | 111 | 112 | 92 | 94 | 93 | 104 | 87 | 84 | 81 | 96 | 88 | 85 | 99 | 103 | 100 | 94 | 93 | 93 | 95 | 95 | 69 | 94 | 102 | 97 | 96 |
| **CAN02** | 76 | 93 | 105 | 95 | 75 | 109 | 89 | 105 | 111 | 86 | 82 | 79 | 39 | 88 | 86 | 90 | 65 | 83 | 95 | 96 | 89 | 92 | 92 | 95 | 101 | 93 | 94 | 84 | 90 | 94 | 94 |
| **CAN03** | 98 | 98 | 91 | 93 | 92 | 97 | 95 | 97 | 74 | 93 | 91 | 85 | 52 | 94 | 88 | 91 | 79 | 89 | 81 | 91 | 82 | 103 | 101 | 89 | 96 | 91 | 72 | 76 | 100 | 97 | 103 |
| **CAN04** | 63 | 103 | 88 | 48 | 92 | 101 | 114 | 59 | 93 | 85 | 83 | 61 | 80 | 80 | 28 | 95 | 16 | 25 | 81 | 70 | 77 | 91 | 94 | 91 | 107 | 98 | 97 | 63 | 91 | 90 | 40 |
| **CAN05** | 94 | 88 | 113 | 95 | 89 | 116 | 125 | 129 | 63 | 39 | 97 | 85 | 86 | 97 | 13 | 95 | 87 | 80 | 35 | 28 | 75 | 98 | 111 | 92 | 110 | 103 | 110 | 89 | 103 | 103 | 47 |
| **CAN06** | 99 | 79 | 96 | 96 | 93 | 107 | 92 | 94 | 92 | 91 | 89 | 96 | 106 | 105 | 71 | 91 | 95 | 76 | 96 | 95 | 92 | 98 | 94 | 93 | 100 | 97 | 95 | 93 | 102 | 100 | 99 |
| **CAN07** | 111 | 106 | 104 | 101 | 104 | 116 | 116 | 106 | 110 | 93 | 98 | 106 | 112 | 109 | 49 | 97 | 91 | 79 | 111 | 112 | 89 | 100 | 107 | 87 | 92 | 95 | 104 | 87 | 99 | 100 | 91 |
| **CAN08** | 91 | 112 | 84 | 71 | 85 | 67 | 116 | 81 | 99 | 57 | 81 | 90 | 103 | 98 | 22 | 83 | 23 | 41 | 49 | 41 | 85 | 100 | 100 | 98 | 106 | 100 | 103 | 93 | 92 | 100 | 89 |
| **CAN09** | 20 | 76 | 86 | 91 | 83 | 87 | 95 | 96 | 56 | 29 | 80 | 80 | 95 | 82 | 83 | 96 | 84 | 80 | 103 | 99 | 87 | 103 | 106 | 90 | 113 | 104 | 114 | 93 | 97 | 95 | 103 |
| **CAN10** | 108 | 80 | 101 | 90 | 100 | 99 | 87 | 100 | 103 | 92 | 90 | 94 | 91 | 94 | 86 | 94 | 92 | 92 | 25 | 29 | 82 | 104 | 79 | 99 | 109 | 103 | 105 | 97 | 99 | 101 | 87 |
| **CAN11** | 105 | 96 | 94 | 101 | 98 | 106 | 94 | 95 | 103 | 92 | 87 | 98 | 94 | 91 | 82 | 95 | 92 | 88 | 94 | 100 | 83 | 89 | 101 | 84 | 89 | 83 | 92 | 83 | 98 | 100 | 117 |
| **CAN12** | 91 | 87 | 116 | 114 | 130 | 133 | 124 | 128 | 85 | 96 | 91 | 89 | 97 | 91 | 80 | 99 | 93 | 87 | 66 | 81 | 72 | 96 | 107 | 82 | 111 | 100 | 104 | 83 | 90 | 105 | 80 |
| **CAN13** | 2 | 60 | 73 | 54 | 76 | 81 | 112 | 76 | 105 | 88 | 87 | 88 | 26 | 86 | 76 | 90 | 28 | 39 | 102 | 102 | 28 | 112 | 121 | 76 | 96 | 99 | 100 | 92 | 99 | 100 | 102 |
| **CAN14** | 126 | 123 | 118 | 105 | 116 | 112 | 130 | 40 | 97 | 99 | 87 | 84 | 89 | 98 | 71 | 100 | 91 | 78 | 80 | 82 | 76 | 107 | 108 | 92 | 118 | 108 | 108 | 93 | 98 | 95 | 81 |
| **CAN15** | 84 | 118 | 115 | 124 | 110 | 136 | 129 | 113 | 109 | 92 | 90 | 83 | 113 | 20 | 82 | 98 | 94 | 80 | 95 | 98 | 80 | 101 | 111 | 81 | 117 | 110 | 114 | 91 | 93 | 100 | 91 |
| **CAN16** | 23 | 28 | 42 | 81 | 130 | 128 | 56 | 87 | 100 | 86 | 71 | 76 | 37 | 94 | 92 | 83 | 39 | 43 | 86 | 85 | 38 | 90 | 86 | 83 | 99 | 87 | 94 | 82 | 92 | 95 | 79 |
| **CAN17** | 46 | 108 | 105 | 107 | 24 | 32 | 107 | 104 | 101 | 24 | 84 | 89 | 97 | 94 | 92 | 93 | 90 | 83 | 20 | 17 | 81 | 86 | 95 | 30 | 82 | 77 | 35 | 63 | 106 | 16 | 36 |
| **CAN18** | 59 | 115 | 111 | 102 | 83 | 46 | 79 | 117 | 96 | 56 | 94 | 94 | 32 | 97 | 28 | 89 | 31 | 90 | 39 | 37 | 102 | 94 | 91 | 61 | 95 | 66 | 87 | 87 | 52 | 92 | 65 |
| **CAN19** | 20 | 31 | 36 | 60 | 56 | 106 | 106 | 102 | 75 | 32 | 90 | 91 | 22 | 88 | 26 | 94 | 81 | 86 | 22 | 20 | 29 | 93 | 96 | 80 | 91 | 87 | 87 | 55 | 38 | 97 | 47 |
| **CAN20** | 115 | 111 | 98 | 54 | 106 | 30 | 111 | 57 | 74 | 83 | 84 | 89 | 71 | 91 | 14 | 91 | 79 | 76 | 88 | 94 | 92 | 95 | 89 | 46 | 98 | 92 | 88 | 66 | 92 | 94 | 97 |
| **CAN21** | 92 | 86 | 122 | 126 | 144 | 22 | 142 | 119 | 95 | 83 | 89 | 82 | 44 | 88 | 75 | 89 | 89 | 72 | 35 | 30 | 88 | 108 | 109 | 30 | 121 | 119 | 32 | 86 | 28 | 12 | 63 |
| **CAN23** | 78 | 78 | 83 | 65 | 88 | 49 | 92 | 96 | 24 | 98 | 99 | 90 | 93 | 97 | 89 | 99 | 99 | 87 | 84 | 90 | 90 | 97 | 94 | 100 | 98 | 76 | 94 | 100 | 99 | 98 | 96 |
| **CAN24** | 32 | 134 | 40 | 102 | 123 | 117 | 29 | 121 | 100 | 97 | 86 | 83 | 100 | 92 | 85 | 92 | 36 | 80 | 98 | 100 | 85 | 88 | 84 | 76 | 71 | 69 | 70 | 72 | 99 | 91 | 97 |
| **CAN25** | 32 | 30 | 29 | 124 | 124 | 125 | 32 | 125 | 89 | 68 | 90 | 98 | 26 | 129 | 136 | 103 | 97 | 115 | 84 | 91 | 75 | 98 | 97 | 92 | 101 | 97 | 97 | 101 | 95 | 97 | 98 |
| **CAN26** | 94 | 100 | 82 | 92 | 91 | 92 | 113 | 84 | 92 | 92 | 87 | 89 | 93 | 90 | 88 | 93 | 91 | 88 | 114 | 109 | 96 | 102 | 95 | 82 | 82 | 81 | 79 | 80 | 102 | 103 | 104 |
| **CAN27** | 7 | 9 | 6 | 81 | 86 | 85 | 7 | 91 | 24 | 91 | 89 | 94 | 8 | 37 | 81 | 94 | 93 | 83 | 102 | 95 | 8 | 99 | 88 | 66 | 94 | 96 | 90 | 17 | 100 | 101 | 96 |
| **CAN28** | 73 | 65 | 70 | 79 | 79 | 80 | 79 | 80 | 113 | 91 | 93 | 93 | 95 | 94 | 85 | 98 | 94 | 98 | 101 | 92 | 51 | 103 | 101 | 84 | 119 | 103 | 101 | 83 | 99 | 95 | 103 |
| **CAN29** | 22 | 22 | 22 | 99 | 44 | 111 | 116 | 125 | 92 | 45 | 84 | 71 | 31 | 90 | 71 | 92 | 86 | 72 | 36 | 30 | 32 | 94 | 84 | 80 | 95 | 92 | 91 | 80 | 34 | 96 | 36 |
| **CAN30** | 89 | 38 | 86 | 96 | 91 | 89 | 103 | 92 | 15 | 90 | 89 | 87 | 92 | 93 | 76 | 96 | 93 | 76 | 92 | 89 | 71 | 87 | 77 | 71 | 21 | 74 | 69 | 66 | 97 | 89 | 88 |
| **CAN31** | 74 | 78 | 71 | 90 | 82 | 67 | 84 | 89 | 64 | 86 | 84 | 77 | 98 | 102 | 80 | 91 | 86 | 77 | 100 | 92 | 88 | 99 | 93 | 85 | 93 | 98 | 97 | 86 | 106 | 106 | 90 |
| **CAN32** | 52 | 22 | 74 | 87 | 80 | 73 | 86 | 94 | 15 | 80 | 71 | 67 | 100 | 91 | 68 | 79 | 74 | 68 | 99 | 103 | 102 | 96 | 98 | 101 | 33 | 93 | 96 | 101 | 99 | 100 | 103 |
| **CAN33** | 9 | 17 | 15 | 73 | 21 | 64 | 32 | 82 | 57 | 30 | 80 | 88 | 86 | 84 | 88 | 36 | 95 | 95 | 21 | 23 | 28 | 98 | 89 | 73 | 100 | 95 | 90 | 74 | 14 | 88 | 24 |
| **CAN34** | 67 | 30 | 87 | 95 | 93 | 83 | 86 | 90 | 26 | 88 | 85 | 86 | 91 | 87 | 88 | 95 | 92 | 89 | 94 | 100 | 87 | 89 | 79 | 78 | 15 | 72 | 70 | 73 | 101 | 99 | 101 |
| **CAN35** | 72 | 107 | 95 | 32 | 93 | 90 | 91 | 64 | 81 | 64 | 100 | 92 | 84 | 92 | 24 | 97 | 36 | 51 | 14 | 10 | 83 | 93 | 59 | 89 | 100 | 95 | 60 | 86 | 37 | 59 | 86 |
| **CAN36** | 91 | 82 | 79 | 105 | 95 | 79 | 90 | 77 | 69 | 94 | 91 | 87 | 94 | 92 | 80 | 97 | 94 | 84 | 93 | 94 | 81 | 103 | 91 | 77 | 99 | 91 | 83 | 75 | 95 | 91 | 90 |
| **CAN37** | 111 | 113 | 124 | 92 | 121 | 121 | 119 | 124 | 102 | 96 | 90 | 88 | 88 | 94 | 92 | 93 | 89 | 88 | 102 | 97 | 98 | 101 | 91 | 95 | 93 | 79 | 74 | 75 | 98 | 112 | 42 |
| **CAN38** | 31 | 98 | 95 | 92 | 97 | 115 | 98 | 95 | 91 | 96 | 99 | 95 | 97 | 99 | 97 | 94 | 94 | 97 | 95 | 95 | 102 | 101 | 70 | 98 | 107 | 85 | 75 | 83 | 87 | 77 | 74 |
| **CAN39** | 23 | 84 | 88 | 98 | 107 | 118 | 86 | 79 | 73 | 98 | 103 | 96 | 100 | 101 | 84 | 98 | 99 | 85 | 97 | 95 | 95 | 101 | 81 | 99 | 101 | 101 | 102 | 98 | 107 | 105 | 96 |
| **CAN40** | 92 | 30 | 86 | 91 | 82 | 78 | 99 | 27 | 17 | 87 | 79 | 88 | 23 | 90 | 80 | 91 | 86 | 79 | 101 | 105 | 98 | 98 | 75 | 96 | 98 | 98 | 99 | 103 | 96 | 94 | 95 |
| **CAN41** | 81 | 81 | 106 | 61 | 8 | 22 | 101 | 72 | 101 | 17 | 85 | 93 | 70 | 112 | 13 | 82 | 55 | 54 | 13 | 13 | 82 | 107 | 108 | 101 | 81 | 106 | 107 | 103 | 14 | 21 | 15 |
| **CAN43** | 96 | 101 | 93 | 91 | 92 | 76 | 93 | 91 | 84 | 93 | 86 | 86 | 91 | 82 | 76 | 90 | 85 | 80 | 109 | 102 | 85 | 95 | 95 | 79 | 85 | 89 | 87 | 77 | 99 | 95 | 101 |
| **CAN44** | 29 | 79 | 77 | 93 | 90 | 78 | 71 | 87 | 19 | 90 | 92 | 86 | 88 | 92 | 79 | 90 | 90 | 80 | 86 | 70 | 67 | 62 | 90 | 75 | 109 | 99 | 88 | 80 | 93 | 86 | 74 |

| | | | | | | | | | | | | | | | | | | | | | | | | | | | | | | | |
|---|---|---|---|---|---|---|---|---|---|---|---|---|---|---|---|---|---|---|---|---|---|---|---|---|---|---|---|---|---|---|---|
| **CAN45** | 20 | 23 | 30 | 71 | 25 | 91 | 70 | 95 | 87 | 21 | 85 | 90 | 17 | 90 | 34 | 89 | 83 | 82 | 21 | 20 | 22 | 98 | 94 | 84 | 95 | 91 | 83 | 76 | 24 | 91 | 41 |
| **CAN46** | 22 | 86 | 91 | 91 | 95 | 106 | 90 | 91 | 23 | 102 | 82 | 76 | 92 | 12 | 66 | 101 | 87 | 69 | 91 | 91 | 85 | 65 | 107 | 99 | 107 | 101 | 98 | 98 | 99 | 98 | 91 |
| **CAN47** | 23 | 79 | 77 | 31 | 79 | 13 | 89 | 16 | 76 | 92 | 94 | 102 | 80 | 97 | 25 | 89 | 91 | 93 | 97 | 85 | 86 | 98 | 85 | 75 | 84 | 78 | 75 | 68 | 90 | 89 | 87 |
| **CAN48** | 15 | 15 | 45 | 77 | 90 | 67 | 17 | 87 | 69 | 100 | 94 | 92 | 13 | 100 | 73 | 105 | 43 | 69 | 89 | 96 | 102 | 109 | 104 | 99 | 112 | 109 | 109 | 96 | 99 | 100 | 85 |
| **CAN49** | 17 | 18 | 17 | 74 | 22 | 58 | 77 | 62 | 53 | 24 | 79 | 93 | 26 | 95 | 63 | 84 | 80 | 74 | 45 | 46 | 50 | 99 | 96 | 99 | 98 | 96 | 94 | 95 | 45 | 95 | 58 |
| **CAN50** | 91 | 78 | 96 | 101 | 73 | 97 | 101 | 97 | 104 | 85 | 84 | 90 | 87 | 110 | 109 | 88 | 81 | 94 | 83 | 82 | 40 | 105 | 105 | 87 | 110 | 103 | 104 | 102 | 100 | 102 | 98 |
| **CAN51** | 17 | 24 | 57 | 53 | 71 | 75 | 66 | 67 | 66 | 78 | 93 | 93 | 85 | 94 | 85 | 100 | 95 | 91 | 17 | 19 | 36 | 95 | 95 | 74 | 73 | 69 | 65 | 63 | 101 | 97 | 89 |
| **CAN52** | 98 | 100 | 101 | 104 | 101 | 97 | 100 | 60 | 65 | 95 | 93 | 95 | 98 | 99 | 97 | 98 | 97 | 98 | 101 | 101 | 104 | 95 | 67 | 97 | 96 | 96 | 92 | 95 | 102 | 97 | 97 |
| **CAN53** | 73 | 91 | 87 | 96 | 95 | 86 | 91 | 88 | 87 | 94 | 94 | 94 | 107 | 98 | 91 | 100 | 96 | 85 | 98 | 95 | 81 | 108 | 102 | 71 | 101 | 90 | 87 | 70 | 100 | 96 | 101 |
| **CAN54** | 42 | 94 | 103 | 94 | 101 | 119 | 97 | 100 | 93 | 95 | 95 | 91 | 100 | 93 | 88 | 99 | 97 | 89 | 82 | 73 | 69 | 99 | 71 | 77 | 106 | 97 | 92 | 74 | 94 | 92 | 80 |
| **CAN55** | 15 | 83 | 91 | 88 | 93 | 31 | 90 | 26 | 93 | 50 | 92 | 92 | 95 | 82 | 100 | 97 | 96 | 95 | 50 | 48 | 66 | 91 | 87 | 67 | 96 | 81 | 69 | 56 | 95 | 88 | 109 |
| **CAN56** | 98 | 95 | 88 | 95 | 93 | 82 | 98 | 94 | 87 | 101 | 102 | 100 | 98 | 101 | 95 | 101 | 100 | 102 | 84 | 78 | 76 | 107 | 102 | 93 | 116 | 106 | 98 | 99 | 95 | 92 | 83 |
| **CAN57** | 20 | 78 | 68 | 91 | 84 | 76 | 14 | 85 | 85 | 98 | 88 | 97 | 91 | 91 | 86 | 97 | 62 | 89 | 99 | 97 | 84 | 84 | 81 | 78 | 86 | 77 | 75 | 76 | 99 | 91 | 98 |
| **CAN58** | 38 | 85 | 54 | 119 | 124 | 142 | 46 | 121 | 33 | 72 | 62 | 57 | 32 | 89 | 69 | 93 | 89 | 62 | 86 | 77 | 46 | 107 | 102 | 82 | 111 | 105 | 99 | 88 | 96 | 91 | 75 |
| **CAN59** | 12 | 10 | 14 | 93 | 88 | 89 | 9 | 90 | 85 | 91 | 88 | 87 | 15 | 88 | 85 | 91 | 13 | 96 | 95 | 96 | 81 | 108 | 92 | 78 | 100 | 99 | 92 | 76 | 100 | 96 | 92 |
| **CAN60** | 21 | 19 | 37 | 85 | 29 | 113 | 97 | 102 | 92 | 25 | 117 | 111 | 18 | 96 | 31 | 90 | 90 | 98 | 24 | 23 | 24 | 91 | 81 | 77 | 101 | 93 | 84 | 70 | 22 | 87 | 29 |
| **CAN61** | 14 | 13 | 46 | 89 | 95 | 77 | 11 | 89 | 85 | 90 | 82 | 76 | 12 | 84 | 76 | 93 | 48 | 78 | 91 | 79 | 61 | 87 | 73 | 57 | 86 | 80 | 57 | 51 | 93 | 75 | 80 |
| **CAN62** | 19 | 56 | 34 | 83 | 63 | 31 | 83 | 51 | 29 | 87 | 87 | 85 | 97 | 90 | 84 | 87 | 92 | 74 | 90 | 78 | 70 | 102 | 95 | 79 | 107 | 103 | 96 | 10 | 100 | 94 | 80 |
| **CAN63** | 84 | 41 | 98 | 114 | 120 | 124 | 122 | 103 | 19 | 83 | 78 | 72 | 90 | 76 | 71 | 91 | 80 | 71 | 110 | 111 | 91 | 96 | 91 | 84 | 22 | 81 | 81 | 80 | 103 | 99 | 103 |
| **CAN64** | 20 | 34 | 27 | 54 | 92 | 94 | 29 | 99 | 112 | 94 | 94 | 30 | 33 | 96 | 88 | 94 | 43 | 42 | 89 | 81 | 79 | 104 | 96 | 90 | 107 | 102 | 97 | 97 | 96 | 93 | 81 |
| **CAN65** | 28 | 40 | 63 | 70 | 104 | 82 | 45 | 49 | 61 | 93 | 86 | 88 | 93 | 92 | 86 | 55 | 88 | 85 | 100 | 77 | 86 | 62 | 102 | 83 | 104 | 95 | 86 | 83 | 91 | 90 | 88 |
| **CAN66** | 28 | 74 | 93 | 95 | 100 | 98 | 110 | 111 | 21 | 91 | 87 | 77 | 92 | 19 | 76 | 95 | 90 | 79 | 82 | 71 | 71 | 48 | 84 | 81 | 110 | 105 | 92 | 84 | 101 | 88 | 78 |
| **CAN67** | 93 | 91 | 95 | 56 | 89 | 92 | 94 | 97 | 85 | 95 | 94 | 93 | 97 | 64 | 98 | 92 | 91 | 93 | 96 | 93 | 99 | 99 | 93 | 94 | 96 | 94 | 90 | 71 | 89 | 84 | 87 |
| **CAN68** | 47 | 99 | 97 | 90 | 94 | 94 | 96 | 26 | 97 | 94 | 94 | 89 | 102 | 66 | 92 | 98 | 97 | 91 | 89 | 79 | 78 | 92 | 85 | 81 | 97 | 92 | 83 | 81 | 93 | 87 | 82 |
| **CAN69** | 97 | 96 | 110 | 119 | 117 | 117 | 119 | 121 | 106 | 87 | 84 | 89 | 103 | 95 | 93 | 92 | 88 | 82 | 99 | 94 | 88 | 94 | 82 | 78 | 85 | 79 | 74 | 77 | 103 | 92 | 91 |
| **CAN70** | 68 | 79 | 84 | 97 | 98 | 100 | 104 | 104 | 33 | 88 | 88 | 90 | 116 | 108 | 110 | 91 | 87 | 95 | 88 | 67 | 64 | 95 | 25 | 69 | 102 | 95 | 81 | 71 | 95 | 85 | 70 |
| **CAN71** | 26 | 30 | 45 | 70 | 51 | 112 | 105 | 111 | 105 | 30 | 76 | 76 | 18 | 83 | 26 | 90 | 82 | 79 | 22 | 22 | 25 | 98 | 89 | 77 | 96 | 92 | 86 | 61 | 22 | 90 | 41 |
| **CAN72** | 26 | 85 | 109 | 126 | 125 | 134 | 149 | 105 | 84 | 95 | 101 | 91 | 90 | 88 | 81 | 93 | 91 | 90 | 74 | 60 | 58 | 96 | 63 | 73 | 106 | 93 | 78 | 69 | 94 | 84 | 68 |
| **CAN73** | 30 | 33 | 43 | 89 | 37 | 118 | 54 | 118 | 112 | 26 | 91 | 89 | 18 | 94 | 30 | 91 | 70 | 86 | 28 | 27 | 20 | 81 | 84 | 87 | 84 | 80 | 76 | 72 | 21 | 98 | 45 |
| **CAN74** | 24 | 25 | 34 | 48 | 110 | 78 | 30 | 121 | 118 | 94 | 88 | 29 | 36 | 71 | 77 | 95 | 42 | 35 | 95 | 92 | 45 | 101 | 99 | 89 | 98 | 99 | 99 | 96 | 102 | 104 | 83 |
| **CAN75** | 59 | 83 | 115 | 71 | 120 | 44 | 130 | 129 | 96 | 96 | 93 | 92 | 93 | 93 | 88 | 93 | 92 | 88 | 98 | 102 | 107 | 95 | 98 | 106 | 96 | 88 | 96 | 92 | 97 | 99 | 108 |
| **CAN76** | 29 | 16 | 31 | 81 | 108 | 109 | 89 | 130 | 117 | 98 | 100 | 98 | 98 | 104 | 85 | 99 | 96 | 89 | 86 | 67 | 61 | 93 | 5 | 71 | 106 | 93 | 76 | 71 | 94 | 78 | 64 |
| **CAN77** | 30 | 31 | 34 | 63 | 65 | 133 | 119 | 124 | 95 | 81 | 84 | 83 | 29 | 76 | 51 | 96 | 42 | 85 | 56 | 59 | 17 | 95 | 60 | 55 | 97 | 88 | 67 | 49 | 70 | 72 | 68 |
| **CAN78** | 98 | 97 | 121 | 122 | 128 | 108 | 130 | 134 | 106 | 85 | 71 | 73 | 104 | 90 | 87 | 91 | 78 | 75 | 91 | 88 | 88 | 103 | 101 | 94 | 110 | 81 | 99 | 97 | 99 | 98 | 90 |
| **CAN79** | 26 | 27 | 28 | 111 | 28 | 111 | 93 | 114 | 112 | 19 | 88 | 88 | 19 | 93 | 77 | 91 | 89 | 89 | 27 | 27 | 21 | 100 | 98 | 69 | 97 | 91 | 85 | 82 | 17 | 86 | 31 |
| **CAN80** | 22 | 39 | 112 | 68 | 116 | 112 | 111 | 122 | 113 | 98 | 97 | 97 | 88 | 93 | 90 | 97 | 89 | 82 | 86 | 73 | 29 | 101 | 91 | 86 | 107 | 100 | 91 | 35 | 96 | 89 | 62 |
| **CAN81** | 20 | 90 | 114 | 97 | 114 | 119 | 126 | 129 | 111 | 100 | 97 | 96 | 91 | 95 | 89 | 101 | 97 | 95 | 96 | 86 | 71 | 97 | 68 | 68 | 85 | 77 | 72 | 65 | 108 | 88 | 86 |
| **CAN82** | 88 | 86 | 102 | 79 | 115 | 55 | 127 | 133 | 136 | 93 | 85 | 72 | 89 | 85 | 69 | 94 | 85 | 71 | 88 | 85 | 81 | 103 | 102 | 81 | 112 | 51 | 102 | 102 | 96 | 94 | 90 |
| **CAN83** | 74 | 96 | 90 | 55 | 92 | 36 | 94 | 94 | 92 | 96 | 95 | 93 | 99 | 99 | 99 | 98 | 95 | 96 | 101 | 96 | 93 | 100 | 97 | 49 | 98 | 30 | 98 | 46 | 86 | 78 | 86 |
| **CAN84** | 96 | 95 | 98 | 104 | 105 | 104 | 109 | 107 | 102 | 94 | 89 | 84 | 98 | 99 | 91 | 93 | 91 | 86 | 95 | 95 | 93 | 100 | 99 | 95 | 100 | 97 | 92 | 92 | 93 | 93 | 90 |
| **CAN85** | 90 | 92 | 108 | 106 | 121 | 38 | 130 | 126 | 123 | 100 | 91 | 88 | 98 | 93 | 71 | 90 | 81 | 65 | 96 | 97 | 80 | 102 | 98 | 19 | 99 | 99 | 92 | 38 | 102 | 97 | 95 |
| **CAN86** | 7 | 10 | 75 | 88 | 90 | 88 | 88 | 89 | 85 | 97 | 94 | 92 | 100 | 98 | 95 | 96 | 95 | 93 | 100 | 98 | 88 | 96 | 84 | 96 | 96 | 96 | 95 | 93 | 104 | 102 | 98 |
| **CAN88** | 96 | 94 | 99 | 103 | 103 | 104 | 105 | 105 | 100 | 90 | 93 | 89 | 95 | 92 | 89 | 91 | 93 | 91 | 80 | 69 | 67 | 79 | 72 | 54 | 81 | 49 | 60 | 57 | 92 | 83 | 85 |
| **CAN89** | 25 | 87 | 108 | 111 | 122 | 122 | 124 | 115 | 102 | 97 | 96 | 98 | 94 | 90 | 83 | 97 | 94 | 90 | 86 | 84 | 85 | 102 | 86 | 96 | 111 | 98 | 98 | 99 | 96 | 92 | 87 |
| **CAN90** | 92 | 91 | 107 | 58 | 122 | 34 | 129 | 119 | 92 | 90 | 81 | 73 | 91 | 82 | 65 | 91 | 82 | 69 | 99 | 100 | 100 | 95 | 93 | 57 | 86 | 28 | 78 | 76 | 103 | 102 | 106 |
| **CAN91** | 15 | 15 | 71 | 20 | 92 | 21 | 96 | 103 | 113 | 95 | 93 | 99 | 15 | 91 | 86 | 91 | 28 | 17 | 92 | 88 | 12 | 97 | 85 | 59 | 104 | 95 | 86 | 89 | 95 | 96 | 88 |
| **CAN92** | 19 | 38 | 43 | 77 | 117 | 116 | 110 | 117 | 112 | 93 | 94 | 86 | 26 | 102 | 73 | 92 | 83 | 77 | 80 | 76 | 25 | 101 | 89 | 62 | 105 | 99 | 87 | 58 | 84 | 65 | 63 |

| | | | | | | | | | | | | | | | | | | | | | | | | | | | | | | | |
|---|---|---|---|---|---|---|---|---|---|---|---|---|---|---|---|---|---|---|---|---|---|---|---|---|---|---|---|---|---|---|---|
| **CAN93** | 95 | 90 | 116 | 115 | 115 | 117 | 130 | 118 | 102 | 86 | 81 | 84 | 103 | 97 | 84 | 86 | 83 | 77 | 87 | 102 | 81 | 93 | 46 | 75 | 96 | 80 | 79 | 70 | 92 | 103 | 93 |
| **CAN94** | 90 | 88 | 94 | 110 | 111 | 113 | 126 | 124 | 38 | 83 | 86 | 86 | 93 | 73 | 96 | 89 | 83 | 90 | 96 | 83 | 73 | 27 | 67 | 68 | 59 | 74 | 66 | 60 | 100 | 95 | 102 |
| **CAN95** | 16 | 57 | 129 | 132 | 188 | 208 | 155 | 92 | 83 | 96 | 101 | 64 | 77 | 94 | 70 | 98 | 98 | 61 | 65 | 70 | 45 | 116 | 112 | 103 | 117 | 114 | 117 | 67 | 97 | 101 | 57 |
| **CAN96** | 29 | 27 | 31 | 97 | 111 | 113 | 29 | 127 | 113 | 92 | 90 | 39 | 27 | 86 | 82 | 96 | 28 | 73 | 86 | 80 | 80 | 94 | 92 | 82 | 96 | 94 | 81 | 75 | 97 | 84 | 89 |
| **CAN97** | 26 | 27 | 59 | 99 | 102 | 100 | 29 | 114 | 80 | 93 | 94 | 92 | 25 | 95 | 88 | 93 | 90 | 83 | 77 | 70 | 71 | 78 | 62 | 77 | 84 | 80 | 74 | 73 | 91 | 89 | 90 |
| **CAN98** | 94 | 91 | 58 | 80 | 86 | 67 | 97 | 16 | 81 | 82 | 26 | 90 | 98 | 101 | 87 | 65 | 93 | 85 | 59 | 44 | 73 | 7 | 93 | 74 | 15 | 50 | 65 | 98 | 45 | 70 | 52 |
| **CAN99** | 98 | 94 | 107 | 119 | 121 | 121 | 116 | 115 | 101 | 97 | 98 | 82 | 99 | 99 | 85 | 96 | 96 | 81 | 80 | 73 | 78 | 95 | 100 | 95 | 116 | 102 | 98 | 94 | 94 | 91 | 84 |
| **CAN100** | 100 | 99 | 80 | 56 | 86 | 50 | 92 | 92 | 99 | 88 | 88 | 86 | 98 | 99 | 99 | 96 | 88 | 59 | 65 | 67 | 78 | 92 | 91 | 94 | 96 | 86 | 82 | 87 | 65 | 88 | 65 |
| **CAN101** | 91 | 78 | 82 | 87 | 18 | 86 | 90 | 79 | 107 | 6 | 88 | 85 | 104 | 98 | 80 | 91 | 84 | 82 | 16 | 11 | 68 | 101 | 77 | 82 | 108 | 96 | 90 | 80 | 91 | 19 | 84 |
| **CAN102** | 34 | 82 | 88 | 85 | 100 | 100 | 108 | 109 | 110 | 86 | 89 | 93 | 106 | 62 | 107 | 98 | 98 | 97 | 91 | 87 | 93 | 99 | 87 | 100 | 113 | 106 | 91 | 45 | 94 | 90 | 96 |
| **CAN103** | 34 | 31 | 32 | 87 | 91 | 89 | 34 | 97 | 96 | 85 | 72 | 28 | 32 | 71 | 64 | 92 | 33 | 65 | 87 | 86 | 87 | 106 | 103 | 99 | 121 | 112 | 106 | 103 | 90 | 87 | 83 |
| **CAN104** | 6 | 15 | 86 | 98 | 97 | 98 | 99 | 102 | 102 | 94 | 95 | 96 | 98 | 99 | 98 | 99 | 97 | 99 | 97 | 95 | 90 | 96 | 75 | 93 | 97 | 101 | 100 | 96 | 99 | 94 | 88 |
| **CAN105** | 11 | 22 | 18 | 51 | 87 | 76 | 88 | 90 | 79 | 89 | 89 | 87 | 11 | 88 | 26 | 94 | 17 | 85 | 43 | 45 | 26 | 93 | 80 | 67 | 101 | 98 | 87 | 64 | 66 | 85 | 64 |
| **CAN106** | 16 | 18 | 25 | 44 | 86 | 67 | 27 | 87 | 58 | 55 | 86 | 81 | 23 | 88 | 78 | 96 | 27 | 32 | 35 | 39 | 27 | 97 | 90 | 48 | 97 | 89 | 85 | 76 | 48 | 103 | 56 |
| **CAN107** | 38 | 95 | 90 | 89 | 89 | 78 | 98 | 92 | 79 | 88 | 93 | 92 | 93 | 87 | 76 | 97 | 93 | 78 | 85 | 85 | 81 | 109 | 106 | 101 | 118 | 117 | 109 | 100 | 90 | 89 | 80 |
| **CAN108** | 96 | 94 | 85 | 101 | 96 | 71 | 89 | 91 | 78 | 98 | 95 | 91 | 94 | 105 | 76 | 98 | 88 | 76 | 99 | 99 | 79 | 93 | 95 | 78 | 84 | 79 | 77 | 69 | 96 | 95 | 96 |
| **CAN109** | 38 | 84 | 83 | 102 | 98 | 72 | 83 | 72 | 24 | 100 | 93 | 87 | 107 | 90 | 81 | 88 | 85 | 71 | 96 | 101 | 100 | 98 | 101 | 102 | 96 | 98 | 105 | 105 | 106 | 112 | 104 |
| **CAN110** | 19 | 43 | 77 | 114 | 112 | 113 | 52 | 85 | 96 | 84 | 78 | 90 | 112 | 108 | 118 | 96 | 83 | 94 | 89 | 87 | 30 | 96 | 86 | 86 | 126 | 124 | 117 | 81 | 92 | 90 | 85 |
| **CAN111** | 23 | 43 | 66 | 87 | 89 | 87 | 34 | 98 | 93 | 92 | 87 | 87 | 28 | 83 | 80 | 97 | 88 | 86 | 100 | 106 | 78 | 98 | 101 | 104 | 96 | 95 | 98 | 98 | 99 | 100 | 96 |
| **CAN112** | 33 | 94 | 95 | 78 | 98 | 102 | 91 | 94 | 98 | 92 | 94 | 91 | 98 | 99 | 97 | 94 | 94 | 98 | 98 | 95 | 100 | 101 | 99 | 102 | 100 | 99 | 95 | 48 | 103 | 100 | 105 |
| **CAN113** | 11 | 19 | 18 | 95 | 21 | 85 | 76 | 91 | 87 | 19 | 84 | 75 | 23 | 88 | 63 | 91 | 85 | 75 | 33 | 35 | 36 | 113 | 127 | 125 | 116 | 111 | 127 | 125 | 19 | 111 | 32 |
| **CAN114** | 11 | 33 | 99 | 47 | 43 | 36 | 36 | 80 | 91 | 38 | 71 | 63 | 31 | 79 | 63 | 91 | 28 | 65 | 31 | 29 | 85 | 98 | 91 | 48 | 101 | 93 | 87 | 32 | 32 | 96 | 56 |
| **CAN115** | 87 | 91 | 92 | 95 | 96 | 90 | 95 | 50 | 83 | 91 | 91 | 90 | 97 | 92 | 86 | 91 | 92 | 89 | 85 | 79 | 54 | 105 | 45 | 91 | 110 | 109 | 103 | 93 | 97 | 94 | 83 |
| **CAN116** | 95 | 59 | 73 | 94 | 104 | 90 | 75 | 108 | 99 | 105 | 112 | 97 | 91 | 86 | 91 | 102 | 108 | 85 | 90 | 87 | 27 | 96 | 96 | 81 | 130 | 130 | 128 | 97 | 108 | 103 | 101 |
| **CAN117** | 16 | 9 | 6 | 29 | 56 | 31 | 42 | 14 | 105 | 24 | 73 | 68 | 18 | 9 | 75 | 17 | 10 | 9 | 54 | 41 | 12 | 37 | 70 | 30 | 19 | 93 | 24 | 77 | 71 | 30 | 80 |
| **CAN118** | 97 | 94 | 110 | 125 | 121 | 129 | 92 | 101 | 111 | 99 | 86 | 102 | 108 | 113 | 118 | 91 | 87 | 95 | 87 | 81 | 76 | 112 | 109 | 94 | 116 | 112 | 109 | 97 | 92 | 90 | 77 |
| **CAN119** | 31 | 27 | 43 | 43 | 81 | 80 | 71 | 96 | 21 | 89 | 69 | 57 | 39 | 45 | 61 | 26 | 33 | 27 | 75 | 53 | 24 | 81 | 57 | 56 | 38 | 84 | 29 | 51 | 81 | 64 | 53 |
| **CAN120** | 91 | 89 | 77 | 60 | 36 | 67 | 95 | 92 | 86 | 19 | 95 | 82 | 92 | 91 | 20 | 95 | 100 | 89 | 20 | 6 | 66 | 96 | 89 | 68 | 99 | 99 | 88 | 65 | 10 | 9 | 9 |
| **CAN121** | 90 | 82 | 88 | 36 | 83 | 27 | 92 | 82 | 67 | 94 | 88 | 93 | 92 | 90 | 89 | 95 | 91 | 94 | 84 | 80 | 76 | 95 | 92 | 75 | 108 | 36 | 91 | 51 | 92 | 85 | 83 |
| **CAN122** | 95 | 64 | 79 | 98 | 87 | 76 | 95 | 85 | 73 | 96 | 91 | 95 | 86 | 94 | 83 | 100 | 94 | 87 | 94 | 86 | 44 | 92 | 95 | 85 | 117 | 113 | 101 | 88 | 100 | 97 | 101 |
| **CAN123** | 92 | 83 | 80 | 92 | 82 | 13 | 89 | 84 | 71 | 89 | 90 | 90 | 89 | 91 | 80 | 96 | 92 | 82 | 81 | 69 | 61 | 96 | 84 | 65 | 102 | 95 | 84 | 68 | 93 | 84 | 72 |
| **CAN124** | 43 | 61 | 59 | 93 | 82 | 63 | 81 | 70 | 21 | 82 | 74 | 75 | 91 | 103 | 78 | 93 | 86 | 74 | 86 | 85 | 66 | 31 | 72 | 66 | 78 | 69 | 67 | 63 | 101 | 102 | 104 |
| **CAN125** | 90 | 91 | 84 | 99 | 90 | 95 | 87 | 85 | 85 | 66 | 79 | 77 | 98 | 65 | 91 | 89 | 79 | 80 | 104 | 66 | 84 | 96 | 92 | 87 | 101 | 84 | 100 | 89 | 96 | 93 | 88 |
| **CAN126** | 44 | 89 | 89 | 82 | 95 | 96 | 87 | 10 | 73 | 82 | 78 | 82 | 98 | 94 | 95 | 87 | 88 | 88 | 86 | 92 | 95 | 94 | 89 | 35 | 99 | 92 | 90 | 28 | 95 | 93 | 98 |
| **CAN127** | 21 | 30 | 87 | 41 | 82 | 12 | 38 | 80 | 89 | 49 | 85 | 84 | 37 | 16 | 7 | 94 | 50 | 74 | 41 | 40 | 56 | 89 | 86 | 40 | 92 | 84 | 57 | 64 | 42 | 91 | 43 |
| **CAN128** | 51 | 92 | 90 | 89 | 88 | 85 | 95 | 102 | 39 | 94 | 92 | 92 | 90 | 89 | 87 | 93 | 98 | 95 | 96 | 93 | 82 | 101 | 98 | 81 | 105 | 110 | 94 | 32 | 106 | 107 | 113 |
| **CAN129** | 98 | 94 | 83 | 91 | 79 | 37 | 101 | 95 | 88 | 49 | 90 | 87 | 85 | 71 | 82 | 96 | 90 | 91 | 35 | 43 | 59 | 111 | 81 | 52 | 118 | 97 | 94 | 73 | 44 | 57 | 42 |
| **CAN130** | 100 | 94 | 90 | 88 | 88 | 79 | 102 | 95 | 87 | 95 | 93 | 97 | 93 | 91 | 85 | 97 | 93 | 89 | 88 | 89 | 72 | 90 | 94 | 74 | 95 | 93 | 84 | 73 | 99 | 91 | 106 |
| **CAN131** | 92 | 91 | 85 | 98 | 39 | 95 | 94 | 92 | 94 | 18 | 90 | 90 | 100 | 98 | 98 | 94 | 95 | 95 | 14 | 14 | 89 | 90 | 101 | 97 | 94 | 102 | 80 | 78 | 75 | 74 | 75 |
| **CAN132** | 25 | 92 | 90 | 100 | 99 | 94 | 92 | 93 | 88 | 93 | 93 | 90 | 104 | 102 | 91 | 98 | 93 | 91 | 86 | 74 | 66 | 87 | 71 | 64 | 96 | 83 | 64 | 63 | 88 | 78 | 76 |
| **CAN133** | 89 | 86 | 36 | 111 | 109 | 89 | 69 | 85 | 91 | 84 | 76 | 68 | 103 | 100 | 72 | 83 | 69 | 70 | 75 | 91 | 80 | 89 | 108 | 91 | 106 | 89 | 106 | 85 | 83 | 100 | 79 |
| **CAN134** | 21 | 36 | 76 | 48 | 86 | 24 | 48 | 79 | 77 | 51 | 81 | 100 | 41 | 26 | 10 | 86 | 45 | 64 | 48 | 47 | 61 | 95 | 94 | 61 | 101 | 94 | 86 | 80 | 54 | 100 | 62 |
| **CAN135** | 92 | 93 | 101 | 94 | 91 | 88 | 97 | 93 | 52 | 82 | 77 | 73 | 92 | 88 | 83 | 91 | 92 | 82 | 85 | 79 | 72 | 90 | 19 | 84 | 107 | 88 | 94 | 88 | 90 | 85 | 77 |
| **CAN136** | 28 | 91 | 84 | 92 | 92 | 82 | 92 | 88 | 85 | 92 | 93 | 81 | 97 | 97 | 90 | 100 | 98 | 93 | 91 | 85 | 78 | 89 | 91 | 82 | 85 | 85 | 78 | 82 | 98 | 95 | 107 |
| **CAN137** | 39 | 51 | 88 | 98 | 94 | 85 | 72 | 91 | 89 | 90 | 89 | 88 | 94 | 103 | 102 | 102 | 66 | 97 | 71 | 56 | 53 | 96 | 85 | 72 | 113 | 97 | 82 | 78 | 90 | 74 | 57 |
| **CAN138** | 97 | 95 | 85 | 97 | 94 | 84 | 98 | 95 | 35 | 92 | 93 | 100 | 100 | 97 | 96 | 96 | 97 | 101 | 90 | 78 | 70 | 94 | 10 | 74 | 100 | 86 | 71 | 71 | 93 | 78 | 75 |
| **CAN139** | 8 | 33 | 29 | 64 | 81 | 62 | 49 | 82 | 41 | 94 | 88 | 31 | 23 | 96 | 100 | 97 | 20 | 21 | 80 | 57 | 33 | 84 | 61 | 44 | 69 | 92 | 74 | 9 | 82 | 74 | 55 |

| | | | | | | | | | | | | | | | | | | | | | | | | | | | | | | | |
|---|---|---|---|---|---|---|---|---|---|---|---|---|---|---|---|---|---|---|---|---|---|---|---|---|---|---|---|---|---|---|---|
| **CAN140** | 8 | 9 | 17 | 31 | 77 | 51 | 11 | 68 | 56 | 91 | 83 | 25 | 18 | 91 | 78 | 94 | 18 | 23 | 92 | 85 | 17 | 92 | 92 | 67 | 95 | 90 | 90 | 92 | 95 | 71 | 61 |
| **CAN141** | 90 | 85 | 87 | 105 | 95 | 91 | 96 | 90 | 85 | 92 | 93 | 103 | 97 | 96 | 95 | 97 | 93 | 102 | 100 | 76 | 60 | 84 | 59 | 67 | 95 | 86 | 76 | 68 | 92 | 81 | 72 |
| **CAN142** | 44 | 44 | 88 | 109 | 118 | 135 | 57 | 78 | 101 | 75 | 70 | 76 | 81 | 86 | 98 | 88 | 46 | 82 | 100 | 98 | 84 | 93 | 95 | 79 | 83 | 72 | 71 | 69 | 96 | 92 | 105 |
| **CAN143** | 87 | 92 | 91 | 93 | 89 | 86 | 94 | 96 | 86 | 80 | 82 | 81 | 93 | 89 | 90 | 92 | 85 | 89 | 87 | 85 | 26 | 108 | 102 | 104 | 111 | 102 | 105 | 101 | 91 | 94 | 91 |
| **CAN144** | 91 | 97 | 103 | 89 | 88 | 91 | 92 | 90 | 95 | 87 | 81 | 83 | 96 | 86 | 82 | 89 | 83 | 81 | 89 | 84 | 83 | 96 | 84 | 81 | 103 | 97 | 90 | 78 | 98 | 82 | 93 |
| **CAN145** | 48 | 86 | 74 | 92 | 78 | 68 | 95 | 82 | 69 | 91 | 87 | 84 | 92 | 87 | 86 | 90 | 86 | 85 | 81 | 79 | 84 | 96 | 56 | 88 | 103 | 99 | 111 | 87 | 97 | 95 | 83 |
| **CAN146** | 25 | 90 | 69 | 95 | 94 | 86 | 27 | 95 | 91 | 88 | 88 | 93 | 85 | 88 | 81 | 92 | 84 | 88 | 88 | 86 | 78 | 94 | 95 | 76 | 110 | 108 | 106 | 84 | 100 | 102 | 97 |
| **CAN147** | 92 | 63 | 89 | 46 | 99 | 95 | 96 | 40 | 90 | 89 | 88 | 99 | 97 | 97 | 97 | 96 | 93 | 42 | 92 | 82 | 69 | 94 | 83 | 73 | 100 | 94 | 84 | 72 | 93 | 87 | 78 |
| **CAN148** | 17 | 18 | 23 | 39 | 76 | 85 | 90 | 83 | 76 | 69 | 92 | 90 | 18 | 98 | 18 | 92 | 23 | 32 | 34 | 23 | 79 | 94 | 82 | 76 | 92 | 82 | 78 | 33 | 23 | 94 | 22 |
| **CAN149** | 20 | 32 | 24 | 50 | 46 | 87 | 89 | 91 | 83 | 25 | 98 | 111 | 20 | 106 | 25 | 98 | 25 | 60 | 19 | 21 | 82 | 103 | 102 | 95 | 110 | 101 | 101 | 65 | 26 | 89 | 24 |
| **CAN150** | 86 | 87 | 87 | 78 | 98 | 97 | 93 | 91 | 87 | 85 | 86 | 92 | 91 | 87 | 92 | 96 | 92 | 97 | 94 | 94 | 100 | 99 | 94 | 103 | 102 | 98 | 96 | 34 | 95 | 94 | 99 |
| **CAN151** | 87 | 90 | 86 | 92 | 88 | 88 | 94 | 85 | 84 | 87 | 78 | 69 | 90 | 76 | 69 | 88 | 77 | 71 | 85 | 80 | 77 | 97 | 91 | 89 | 109 | 103 | 93 | 90 | 93 | 89 | 81 |
| **CAN152** | 37 | 43 | 88 | 75 | 94 | 82 | 49 | 91 | 49 | 79 | 79 | 81 | 69 | 101 | 103 | 81 | 81 | 88 | 89 | 87 | 30 | 100 | 67 | 80 | 110 | 103 | 101 | 33 | 113 | 105 | 103 |
| **CAN153** | 96 | 94 | 88 | 84 | 94 | 51 | 99 | 97 | 63 | 97 | 94 | 93 | 96 | 90 | 88 | 89 | 88 | 86 | 81 | 74 | 72 | 100 | 102 | 37 | 109 | 98 | 45 | 88 | 92 | 85 | 80 |
| **CAN154** | 96 | 96 | 97 | 89 | 91 | 89 | 94 | 93 | 85 | 97 | 97 | 97 | 89 | 87 | 87 | 93 | 92 | 90 | 97 | 93 | 90 | 96 | 101 | 101 | 99 | 96 | 91 | 93 | 104 | 103 | 116 |
| **CAN155** | 20 | 26 | 97 | 97 | 131 | 83 | 44 | 90 | 65 | 97 | 95 | 86 | 70 | 93 | 84 | 95 | 92 | 91 | 85 | 77 | 40 | 100 | 91 | 84 | 110 | 94 | 103 | 64 | 94 | 88 | 79 |
| **CAN156** | 74 | 85 | 138 | 83 | 15 | 88 | 96 | 82 | 93 | 8 | 79 | 72 | 77 | 87 | 80 | 86 | 77 | 74 | 5 | 5 | 83 | 88 | 82 | 27 | 87 | 80 | 41 | 71 | 28 | 8 | 22 |
| **CAN157** | 34 | 38 | 64 | 96 | 95 | 71 | 51 | 92 | 59 | 90 | 89 | 79 | 56 | 80 | 69 | 89 | 81 | 71 | 79 | 61 | 29 | 83 | 70 | 65 | 100 | 87 | 78 | 42 | 85 | 78 | 64 |
| **CAN158** | 16 | 62 | 59 | 85 | 67 | 62 | 86 | 64 | 59 | 93 | 91 | 85 | 103 | 101 | 92 | 95 | 93 | 88 | 92 | 91 | 93 | 95 | 98 | 91 | 101 | 96 | 97 | 94 | 94 | 95 | 97 |
| **CAN159** | 90 | 88 | 115 | 91 | 86 | 86 | 106 | 96 | 95 | 91 | 76 | 74 | 106 | 83 | 69 | 88 | 79 | 64 | 78 | 69 | 64 | 97 | 88 | 81 | 106 | 102 | 90 | 81 | 94 | 86 | 74 |
| **CAN160** | 22 | 67 | 73 | 84 | 93 | 83 | 38 | 89 | 30 | 88 | 85 | 96 | 69 | 65 | 103 | 48 | 91 | 84 | 81 | 83 | 80 | 88 | 90 | 78 | 79 | 82 | 66 | 73 | 101 | 98 | 105 |
| **CAN161** | 90 | 91 | 90 | 92 | 93 | 90 | 91 | 90 | 85 | 92 | 91 | 91 | 99 | 98 | 95 | 94 | 94 | 87 | 93 | 84 | 77 | 85 | 83 | 80 | 93 | 85 | 83 | 81 | 91 | 86 | 79 |
| **CAN162** | 23 | 42 | 93 | 55 | 101 | 102 | 28 | 96 | 91 | 94 | 91 | 92 | 98 | 56 | 96 | 94 | 46 | 93 | 90 | 86 | 83 | 93 | 90 | 58 | 106 | 95 | 53 | 38 | 37 | 89 | 64 |
| **CAN163** | 18 | 91 | 92 | 97 | 95 | 91 | 92 | 91 | 86 | 91 | 91 | 87 | 95 | 94 | 92 | 94 | 92 | 91 | 91 | 85 | 83 | 102 | 101 | 88 | 102 | 100 | 98 | 94 | 99 | 95 | 88 |
| **CAN164** | 16 | 35 | 63 | 64 | 30 | 85 | 81 | 87 | 81 | 30 | 89 | 93 | 58 | 93 | 78 | 91 | 92 | 90 | 33 | 32 | 75 | 101 | 91 | 79 | 91 | 88 | 87 | 76 | 93 | 98 | 49 |
| **CAN165** | 29 | 29 | 31 | 94 | 67 | 67 | 72 | 68 | 50 | 78 | 89 | 99 | 77 | 97 | 87 | 92 | 66 | 94 | 93 | 90 | 99 | 96 | 89 | 99 | 95 | 96 | 97 | 104 | 97 | 100 | 103 |
| **CAN166** | 17 | 75 | 83 | 107 | 103 | 101 | 79 | 77 | 85 | 95 | 99 | 98 | 93 | 99 | 93 | 99 | 102 | 90 | 85 | 81 | 84 | 94 | 84 | 86 | 97 | 90 | 87 | 80 | 102 | 91 | 110 |
| **CAN167** | 14 | 11 | 9 | 35 | 10 | 77 | 22 | 12 | 86 | 5 | 79 | 71 | 12 | 90 | 15 | 91 | 15 | 14 | 13 | 15 | 7 | 99 | 88 | 69 | 111 | 108 | 79 | 83 | 13 | 1 | 8 |
| **CAN168** | 91 | 91 | 99 | 87 | 93 | 90 | 94 | 94 | 93 | 93 | 82 | 99 | 136 | 127 | 129 | 85 | 82 | 93 | 89 | 94 | 99 | 102 | 88 | 89 | 100 | 105 | 101 | 18 | 97 | 95 | 106 |
| **CAN169** | 81 | 87 | 94 | 91 | 89 | 90 | 88 | 87 | 72 | 83 | 79 | 78 | 96 | 93 | 90 | 91 | 90 | 84 | 88 | 75 | 66 | 88 | 45 | 68 | 101 | 89 | 74 | 68 | 90 | 74 | 64 |
| **CAN170** | 32 | 32 | 13 | 93 | 93 | 91 | 12 | 93 | 19 | 92 | 93 | 95 | 39 | 92 | 93 | 90 | 91 | 101 | 91 | 86 | 31 | 97 | 99 | 83 | 110 | 104 | 90 | 85 | 103 | 91 | 93 |
| **CAN171** | 96 | 182 | 87 | 169 | 182 | 89 | 174 | 177 | 89 | 94 | 93 | 98 | 96 | 94 | 85 | 95 | 95 | 94 | 106 | 97 | 81 | 90 | 84 | 83 | 93 | 89 | 86 | 82 | 99 | 95 | 85 |
| **CAN172** | 16 | 15 | 18 | 65 | 27 | 78 | 32 | 80 | 56 | 27 | 82 | 80 | 27 | 85 | 24 | 94 | 82 | 70 | 18 | 20 | 17 | 96 | 75 | 64 | 92 | 82 | 73 | 52 | 20 | 89 | 25 |
| **CAN173** | 100 | 98 | 91 | 106 | 94 | 92 | 100 | 98 | 85 | 39 | 82 | 87 | 86 | 81 | 73 | 91 | 86 | 75 | 90 | 29 | 86 | 102 | 95 | 96 | 106 | 102 | 100 | 98 | 30 | 87 | 20 |
| **CAN174** | 20 | 28 | 23 | 69 | 23 | 102 | 77 | 76 | 81 | 27 | 85 | 83 | 25 | 103 | 71 | 93 | 92 | 82 | 20 | 23 | 24 | 95 | 86 | 83 | 91 | 87 | 77 | 74 | 24 | 88 | 38 |
| **CAN175** | 34 | 47 | 13 | 94 | 88 | 86 | 9 | 93 | 32 | 93 | 87 | 75 | 35 | 104 | 71 | 93 | 63 | 76 | 88 | 80 | 29 | 99 | 81 | 66 | 113 | 98 | 84 | 71 | 91 | 82 | 64 |
| **CAN176** | 34 | 41 | 37 | 94 | 84 | 111 | 69 | 99 | 76 | 60 | 76 | 88 | 100 | 103 | 105 | 87 | 67 | 97 | 86 | 74 | 73 | 92 | 75 | 84 | 126 | 124 | 117 | 96 | 92 | 91 | 78 |
| **CAN177** | 5 | 95 | 93 | 89 | 96 | 96 | 97 | 96 | 45 | 86 | 86 | 88 | 93 | 85 | 81 | 103 | 93 | 87 | 87 | 85 | 81 | 99 | 95 | 100 | 108 | 103 | 98 | 99 | 94 | 92 | 89 |
| **CAN178** | 24 | 92 | 93 | 93 | 92 | 91 | 94 | 94 | 90 | 93 | 92 | 92 | 90 | 85 | 84 | 94 | 99 | 94 | 82 | 77 | 77 | 95 | 83 | 77 | 90 | 84 | 76 | 71 | 102 | 91 | 100 |
| **CAN179** | 17 | 95 | 97 | 93 | 90 | 84 | 90 | 92 | 87 | 95 | 91 | 80 | 83 | 88 | 78 | 98 | 84 | 84 | 84 | 76 | 72 | 105 | 97 | 84 | 116 | 110 | 101 | 100 | 95 | 89 | 83 |
| **CAN180** | 3 | 10 | 17 | 22 | 96 | 88 | 8 | 50 | 71 | 91 | 92 | 32 | 21 | 91 | 77 | 99 | 22 | 25 | 87 | 86 | 24 | 100 | 96 | 97 | 96 | 95 | 92 | 92 | 101 | 88 | 98 |
| **CAN181** | 106 | 91 | 88 | 112 | 91 | 80 | 97 | 87 | 86 | 91 | 96 | 95 | 91 | 90 | 79 | 101 | 97 | 85 | 85 | 78 | 73 | 103 | 102 | 82 | 120 | 106 | 102 | 85 | 100 | 94 | 81 |
| **CAN182** | 45 | 80 | 90 | 95 | 96 | 59 | 80 | 79 | 89 | 94 | 91 | 96 | 97 | 91 | 87 | 92 | 86 | 93 | 89 | 78 | 72 | 86 | 80 | 43 | 101 | 39 | 96 | 37 | 93 | 77 | 82 |
| **CAN183** | 33 | 85 | 84 | 59 | 94 | 40 | 101 | 102 | 96 | 113 | 116 | 114 | 103 | 106 | 111 | 106 | 111 | 115 | 94 | 109 | 92 | 86 | 92 | 47 | 97 | 55 | 91 | 40 | 98 | 90 | 84 |
| **CAN184** | 95 | 94 | 97 | 87 | 89 | 85 | 97 | 95 | 92 | 93 | 96 | 92 | 100 | 99 | 82 | 93 | 97 | 93 | 90 | 105 | 90 | 116 | 95 | 79 | 98 | 103 | 103 | 18 | 117 | 111 | 132 |
| **CAN185** | 94 | 43 | 89 | 94 | 91 | 84 | 96 | 59 | 82 | 83 | 78 | 77 | 52 | 76 | 73 | 101 | 92 | 82 | 80 | 75 | 70 | 107 | 87 | 87 | 112 | 106 | 95 | 101 | 99 | 89 | 86 |
| **CAN186** | 100 | 100 | 103 | 106 | 107 | 98 | 103 | 103 | 88 | 96 | 97 | 90 | 96 | 96 | 87 | 92 | 90 | 99 | 94 | 94 | 91 | 99 | 92 | 85 | 86 | 85 | 80 | 73 | 100 | 94 | 104 |

| | | | | | | | | | | | | | | | | | | | | | | | | | | | | | | |
|---|---|---|---|---|---|---|---|---|---|---|---|---|---|---|---|---|---|---|---|---|---|---|---|---|---|---|---|---|---|---|
| **CAN187** | 10 | 22 | 37 | 24 | 100 | 36 | 83 | 16 | 11 | 86 | 95 | 97 | 26 | 98 | 4 | 95 | 43 | 43 | 86 | 80 | 46 | 97 | 33 | 42 | 99 | 94 | 50 | 11 | 34 | 94 | 27 |
| **CAN188** | 9 | 18 | 38 | 22 | 100 | 39 | 92 | 16 | 12 | 85 | 84 | 80 | 28 | 88 | 32 | 92 | 49 | 40 | 74 | 73 | 45 | 100 | 40 | 52 | 102 | 99 | 75 | 11 | 44 | 99 | 32 |
| **CAN189** | 90 | 84 | 86 | 55 | 89 | 65 | 85 | 80 | 78 | 90 | 88 | 81 | 87 | 84 | 71 | 94 | 88 | 69 | 100 | 89 | 82 | 92 | 89 | 96 | 103 | 93 | 94 | 93 | 91 | 91 | 87 |
| **CAN190** | 19 | 20 | 28 | 71 | 22 | 100 | 69 | 72 | 82 | 33 | 83 | 82 | 22 | 103 | 31 | 96 | 70 | 84 | 21 | 22 | 73 | 97 | 99 | 80 | 94 | 89 | 77 | 69 | 47 | 87 | 40 |
| **CAN191** | 92 | 90 | 89 | 100 | 99 | 94 | 94 | 63 | 90 | 90 | 85 | 85 | 102 | 94 | 87 | 91 | 87 | 86 | 91 | 88 | 85 | 86 | 99 | 81 | 96 | 102 | 86 | 97 | 85 | 83 | 76 |
| **CAN192** | 12 | 57 | 93 | 48 | 81 | 22 | 87 | 73 | 87 | 90 | 85 | 85 | 102 | 94 | 87 | 91 | 87 | 86 | 30 | 28 | 106 | 100 | 98 | 90 | 97 | 73 | 89 | 106 | 52 | 96 | 95 |
| **CAN193** | 94 | 95 | 102 | 91 | 92 | 88 | 92 | 93 | 90 | 86 | 80 | 80 | 93 | 84 | 82 | 97 | 79 | 80 | 90 | 87 | 83 | 100 | 103 | 98 | 114 | 105 | 101 | 101 | 93 | 89 | 85 |
| **CAN194** | 50 | 62 | 94 | 40 | 103 | 68 | 97 | 100 | 92 | 92 | 89 | 84 | 86 | 75 | 88 | 28 | 41 | 32 | 101 | 100 | 37 | 101 | 96 | 82 | 31 | 60 | 44 | 94 | 104 | 105 | 99 |
| **CAN195** | 24 | 43 | 87 | 88 | 92 | 78 | 35 | 92 | 79 | 93 | 92 | 97 | 37 | 93 | 75 | 93 | 31 | 82 | 84 | 74 | 66 | 96 | 72 | 75 | 106 | 99 | 74 | 78 | 92 | 80 | 66 |
| **CAN196** | 90 | 90 | 87 | 96 | 99 | 73 | 94 | 87 | 79 | 96 | 88 | 82 | 88 | 85 | 73 | 87 | 84 | 73 | 91 | 86 | 74 | 84 | 83 | 73 | 90 | 85 | 84 | 75 | 116 | 104 | 120 |
| **CAN197** | 22 | 48 | 81 | 120 | 109 | 79 | 54 | 92 | 78 | 103 | 99 | 83 | 89 | 29 | 78 | 44 | 96 | 76 | 89 | 88 | 81 | 102 | 103 | 82 | 58 | 103 | 80 | 26 | 93 | 80 | 98 |
| **CAN198** | 86 | 82 | 89 | 116 | 112 | 112 | 88 | 83 | 97 | 99 | 107 | 103 | 100 | 105 | 89 | 95 | 94 | 78 | 90 | 88 | 90 | 101 | 101 | 99 | 111 | 107 | 102 | 103 | 89 | 92 | 85 |
| **CAN199** | 99 | 96 | 94 | 110 | 106 | 103 | 104 | 99 | 64 | 83 | 68 | 68 | 99 | 77 | 66 | 87 | 72 | 63 | 100 | 104 | 95 | 92 | 90 | 86 | 86 | 83 | 84 | 85 | 99 | 107 | 96 |
| **CAN200** | 29 | 88 | 82 | 96 | 100 | 38 | 93 | 105 | 98 | 92 | 90 | 96 | 103 | 61 | 106 | 89 | 88 | 94 | 86 | 76 | 69 | 98 | 90 | 77 | 102 | 100 | 91 | 84 | 94 | 85 | 72 |
| **CAN201** | 92 | 88 | 86 | 91 | 84 | 80 | 100 | 98 | 101 | 88 | 83 | 86 | 97 | 94 | 89 | 103 | 101 | 95 | 90 | 95 | 92 | 100 | 99 | 89 | 96 | 94 | 87 | 87 | 102 | 101 | 112 |
| **CAN202** | 32 | 94 | 100 | 97 | 103 | 108 | 98 | 98 | 100 | 94 | 95 | 88 | 100 | 99 | 96 | 98 | 98 | 97 | 99 | 99 | 93 | 92 | 82 | 101 | 95 | 91 | 93 | 55 | 96 | 93 | 104 |
| **CAN203** | 92 | 93 | 92 | 102 | 102 | 95 | 97 | 96 | 98 | 87 | 87 | 87 | 93 | 92 | 85 | 88 | 87 | 83 | 91 | 84 | 80 | 88 | 79 | 77 | 95 | 85 | 78 | 77 | 146 | 140 | 139 |
| **CAN204** | 95 | 97 | 88 | 83 | 101 | 73 | 92 | 96 | 101 | 84 | 76 | 69 | 109 | 94 | 71 | 89 | 76 | 70 | 88 | 85 | 79 | 103 | 98 | 75 | 113 | 107 | 43 | 93 | 94 | 89 | 81 |
| **CAN205** | 90 | 80 | 86 | 110 | 107 | 83 | 92 | 92 | 101 | 95 | 90 | 83 | 88 | 87 | 74 | 91 | 94 | 84 | 91 | 83 | 71 | 96 | 94 | 74 | 116 | 104 | 95 | 72 | 118 | 101 | 92 |
| **HU01** | 41 | 59 | 63 | 83 | 77 | 73 | 65 | 107 | 112 | 90 | 85 | 36 | 94 | 92 | 87 | 100 | 92 | 90 | 88 | 76 | 69 | 103 | 105 | 86 | 120 | 109 | 100 | 89 | 94 | 84 | 77 |
| **HU02** | 93 | 93 | 88 | 86 | 83 | 75 | 103 | 97 | 104 | 94 | 96 | 93 | 90 | 93 | 87 | 97 | 93 | 95 | 92 | 82 | 70 | 88 | 90 | 65 | 89 | 87 | 73 | 62 | 103 | 96 | 98 |
| **HU03** | 90 | 93 | 101 | 44 | 88 | 28 | 106 | 105 | 18 | 90 | 91 | 95 | 97 | 20 | 39 | 25 | 93 | 98 | 84 | 70 | 72 | 101 | 36 | 29 | 29 | 71 | 30 | 81 | 90 | 83 | 79 |
| **HU05** | 27 | 23 | 86 | 42 | 94 | 85 | 37 | 98 | 31 | 97 | 96 | 86 | 17 | 73 | 86 | 91 | 97 | 34 | 77 | 73 | 30 | 79 | 95 | 71 | 91 | 96 | 82 | 68 | 88 | 74 | 77 |
| **HU06** | 96 | 32 | 85 | 85 | 86 | 76 | 91 | 87 | 93 | 90 | 88 | 82 | 43 | 86 | 76 | 97 | 90 | 81 | 85 | 88 | 70 | 96 | 79 | 73 | 110 | 96 | 102 | 88 | 90 | 95 | 83 |
| **HU07** | 96 | 98 | 84 | 65 | 95 | 29 | 88 | 82 | 50 | 94 | 93 | 86 | 96 | 95 | 87 | 93 | 94 | 89 | 86 | 85 | 75 | 100 | 90 | 22 | 115 | 110 | 101 | 16 | 100 | 99 | 89 |
| **HU09** | 89 | 78 | 84 | 114 | 106 | 81 | 85 | 82 | 84 | 97 | 106 | 95 | 95 | 62 | 91 | 96 | 105 | 93 | 87 | 85 | 72 | 102 | 100 | 89 | 105 | 99 | 96 | 95 | 99 | 94 | 87 |
| **HU11** | 28 | 14 | 35 | 90 | 75 | 71 | 32 | 76 | 25 | 87 | 86 | 82 | 30 | 92 | 90 | 94 | 89 | 86 | 86 | 77 | 26 | 80 | 73 | 59 | 89 | 76 | 62 | 55 | 92 | 81 | 78 |
| **HU12** | 88 | 90 | 92 | 93 | 93 | 91 | 94 | 95 | 26 | 98 | 91 | 79 | 30 | 98 | 81 | 100 | 99 | 85 | 82 | 79 | 74 | 109 | 114 | 96 | 125 | 113 | 107 | 101 | 96 | 90 | 83 |
| **HU13** | 4 | 19 | 94 | 24 | 92 | 78 | 93 | 92 | 92 | 87 | 90 | 89 | 13 | 91 | 88 | 96 | 12 | 25 | 95 | 88 | 23 | 15 | 93 | 87 | 92 | 72 | 77 | 25 | 97 | 87 | 88 |
| **HU14** | 18 | 19 | 12 | 95 | 91 | 82 | 26 | 94 | 15 | 90 | 92 | 85 | 25 | 49 | 90 | 19 | 94 | 93 | 76 | 60 | 27 | 100 | 73 | 60 | 40 | 91 | 75 | 61 | 87 | 69 | 52 |
| **HU17** | 16 | 9 | 69 | 95 | 90 | 82 | 46 | 89 | 82 | 88 | 84 | 89 | 18 | 92 | 87 | 92 | 90 | 86 | 89 | 91 | 82 | 99 | 101 | 83 | 118 | 122 | 115 | 97 | 96 | 92 | 97 |
| **HU18** | 5 | 22 | 56 | 96 | 92 | 49 | 34 | 93 | 98 | 95 | 108 | 29 | 12 | 101 | 95 | 92 | 57 | 93 | 75 | 63 | 52 | 84 | 75 | 22 | 96 | 89 | 27 | 62 | 85 | 76 | 62 |
| **HU19** | 97 | 98 | 91 | 87 | 100 | 70 | 91 | 91 | 68 | 88 | 78 | 78 | 116 | 98 | 83 | 92 | 94 | 77 | 86 | 83 | 70 | 92 | 84 | 71 | 97 | 90 | 88 | 73 | 103 | 94 | 104 |
| **HU21** | 15 | 77 | 89 | 105 | 93 | 76 | 95 | 93 | 23 | 81 | 76 | 86 | 67 | 21 | 123 | 84 | 80 | 98 | 80 | 72 | 67 | 107 | 92 | 84 | 116 | 111 | 98 | 89 | 96 | 87 | 78 |
| **HU22** | 70 | 77 | 84 | 90 | 91 | 94 | 80 | 76 | 13 | 87 | 89 | 86 | 94 | 93 | 92 | 94 | 90 | 88 | 87 | 85 | 75 | 52 | 68 | 74 | 81 | 84 | 76 | 67 | 98 | 88 | 96 |
| **HU23** | 6 | 73 | 78 | 94 | 90 | 83 | 96 | 86 | 23 | 98 | 97 | 79 | 90 | 24 | 80 | 92 | 93 | 80 | 90 | 73 | 37 | 15 | 78 | 67 | 123 | 114 | 92 | 67 | 92 | 77 | 55 |
| **HU25** | 91 | 92 | 81 | 97 | 91 | 76 | 101 | 98 | 33 | 93 | 77 | 68 | 34 | 97 | 68 | 90 | 69 | 65 | 91 | 86 | 69 | 107 | 110 | 79 | 113 | 112 | 94 | 75 | 108 | 93 | 105 |
| **HU26** | 22 | 44 | 88 | 95 | 86 | 80 | 45 | 88 | 67 | 96 | 93 | 90 | 42 | 95 | 90 | 92 | 27 | 89 | 92 | 81 | 74 | 97 | 71 | 85 | 101 | 98 | 92 | 83 | 96 | 91 | 36 |
| **HU27** | 24 | 32 | 58 | 92 | 94 | 84 | 31 | 87 | 82 | 89 | 80 | 31 | 41 | 88 | 73 | 91 | 81 | 54 | 84 | 72 | 56 | 92 | 69 | 54 | 90 | 86 | 66 | 57 | 96 | 86 | 80 |
| **HU28** | 86 | 76 | 83 | 86 | 77 | 79 | 97 | 82 | 84 | 91 | 94 | 103 | 93 | 89 | 90 | 89 | 87 | 92 | 87 | 73 | 62 | 85 | 70 | 70 | 101 | 88 | 71 | 72 | 88 | 71 | 70 |
| **HU30** | 38 | 96 | 76 | 92 | 96 | 70 | 94 | 97 | 81 | 98 | 90 | 75 | 99 | 102 | 75 | 96 | 93 | 75 | 87 | 85 | 77 | 102 | 85 | 78 | 93 | 99 | 94 | 85 | 100 | 82 | 94 |
| **HU31** | 21 | 41 | 78 | 109 | 103 | 75 | 86 | 79 | 82 | 87 | 81 | 82 | 85 | 89 | 86 | 93 | 90 | 86 | 88 | 75 | 71 | 96 | 82 | 73 | 110 | 97 | 86 | 77 | 88 | 80 | 66 |
| **HU32** | 70 | 57 | 63 | 116 | 87 | 91 | 68 | 61 | 8 | 102 | 67 | 74 | 90 | 16 | 74 | 89 | 97 | 72 | 87 | 83 | 70 | 93 | 96 | 73 | 121 | 119 | 111 | 77 | 109 | 106 | 102 |
| **HU33** | 10 | 10 | 8 | 34 | 77 | 34 | 29 | 89 | 77 | 84 | 77 | 90 | 13 | 114 | 120 | 92 | 14 | 19 | 84 | 77 | 18 | 98 | 103 | 88 | 111 | 104 | 99 | 91 | 97 | 91 | 81 |
| **HU34** | 15 | 15 | 25 | 93 | 89 | 76 | 28 | 91 | 20 | 83 | 84 | 82 | 30 | 85 | 82 | 96 | 95 | 86 | 89 | 82 | 16 | 86 | 80 | 65 | 88 | 77 | 75 | 62 | 99 | 91 | 82 |
| **HU35** | 26 | 50 | 84 | 68 | 89 | 64 | 95 | 91 | 89 | 97 | 95 | 86 | 17 | 86 | 80 | 27 | 33 | 47 | 85 | 67 | 30 | 104 | 97 | 46 | 44 | 113 | 32 | 97 | 98 | 92 | 84 |
| **HU36** | 19 | 50 | 57 | 88 | 70 | 53 | 87 | 77 | 53 | 94 | 93 | 87 | 95 | 92 | 80 | 95 | 94 | 86 | 66 | 82 | 65 | 83 | 113 | 95 | 123 | 108 | 109 | 104 | 92 | 93 | 84 |

| | | | | | | | | | | | | | | | | | | | | | | | | | | | | | | | |
|---|---|---|---|---|---|---|---|---|---|---|---|---|---|---|---|---|---|---|---|---|---|---|---|---|---|---|---|---|---|---|---|
| **HU37** | 97 | 89 | 82 | 88 | 85 | 74 | 89 | 86 | 72 | 92 | 91 | 88 | 28 | 92 | 80 | 98 | 32 | 90 | 88 | 85 | 82 | 101 | 97 | 89 | 112 | 101 | 103 | 90 | 92 | 95 | 79 |
| **HU38** | 27 | 21 | 45 | 95 | 92 | 76 | 23 | 92 | 67 | 84 | 91 | 94 | 42 | 92 | 80 | 92 | 25 | 33 | 90 | 86 | 81 | 96 | 92 | 84 | 120 | 117 | 112 | 89 | 105 | 109 | 101 |
| **HU39** | 7 | 15 | 21 | 32 | 94 | 34 | 23 | 83 | 62 | 58 | 89 | 82 | 27 | 90 | 83 | 90 | 19 | 26 | 45 | 46 | 85 | 95 | 96 | 69 | 96 | 95 | 76 | 90 | 35 | 94 | 11 |
| **HU40** | 79 | 76 | 80 | 119 | 114 | 17 | 86 | 77 | 85 | 99 | 106 | 105 | 96 | 97 | 93 | 44 | 101 | 107 | 88 | 88 | 81 | 84 | 95 | 28 | 90 | 101 | 21 | 91 | 93 | 99 | 108 |
| **HU41** | 96 | 99 | 92 | 94 | 88 | 77 | 95 | 88 | 85 | 82 | 72 | 65 | 109 | 86 | 65 | 75 | 67 | 69 | 82 | 84 | 89 | 111 | 110 | 99 | 114 | 113 | 110 | 109 | 89 | 89 | 82 |
| **HU42** | 29 | 25 | 39 | 30 | 89 | 78 | 25 | 92 | 82 | 101 | 102 | 91 | 31 | 87 | 84 | 96 | 92 | 91 | 22 | 92 | 25 | 103 | 98 | 46 | 101 | 114 | 105 | 99 | 102 | 94 | 108 |
| **HU43** | 36 | 36 | 65 | 84 | 90 | 79 | 46 | 95 | 86 | 90 | 88 | 93 | 63 | 86 | 87 | 94 | 70 | 74 | 92 | 87 | 50 | 103 | 97 | 94 | 116 | 103 | 106 | 93 | 92 | 97 | 82 |
| **HU44** | 77 | 22 | 87 | 97 | 97 | 84 | 93 | 93 | 16 | 97 | 98 | 87 | 95 | 92 | 90 | 93 | 92 | 87 | 86 | 85 | 74 | 42 | 87 | 73 | 95 | 87 | 83 | 71 | 101 | 99 | 101 |
| **HU45** | 97 | 96 | 80 | 95 | 99 | 75 | 97 | 85 | 70 | 82 | 79 | 73 | 82 | 85 | 81 | 90 | 85 | 81 | 85 | 80 | 70 | 102 | 106 | 85 | 115 | 104 | 99 | 86 | 93 | 88 | 74 |
| **HU46** | 50 | 84 | 88 | 117 | 113 | 82 | 95 | 94 | 77 | 102 | 96 | 90 | 88 | 91 | 78 | 93 | 94 | 79 | 93 | 82 | 61 | 91 | 88 | 70 | 93 | 94 | 77 | 69 | 104 | 91 | 94 |
| **HU47** | 18 | 27 | 40 | 36 | 118 | 90 | 28 | 84 | 43 | 94 | 94 | 90 | 9 | 34 | 78 | 51 | 91 | 18 | 81 | 76 | 12 | 47 | 97 | 81 | 55 | 106 | 40 | 94 | 90 | 86 | 79 |
| **HU48** | 93 | 86 | 87 | 95 | 92 | 90 | 102 | 95 | 91 | 90 | 94 | 99 | 93 | 93 | 86 | 93 | 92 | 87 | 77 | 78 | 94 | 95 | 97 | 92 | 98 | 93 | 86 | 91 | 100 | 100 | 104 |
| **HU49** | 95 | 92 | 93 | 93 | 91 | 80 | 96 | 94 | 94 | 103 | 98 | 91 | 92 | 90 | 82 | 94 | 92 | 80 | 82 | 79 | 73 | 101 | 99 | 81 | 111 | 107 | 102 | 86 | 92 | 88 | 72 |
| **HU50** | 92 | 55 | 88 | 97 | 95 | 80 | 94 | 22 | 85 | 110 | 110 | 94 | 64 | 98 | 78 | 102 | 107 | 83 | 90 | 92 | 67 | 99 | 63 | 76 | 118 | 125 | 125 | 93 | 110 | 101 | 112 |
| **HU51** | 36 | 85 | 97 | 92 | 89 | 82 | 62 | 95 | 89 | 94 | 97 | 30 | 43 | 89 | 91 | 91 | 93 | 96 | 88 | 82 | 83 | 99 | 95 | 104 | 112 | 100 | 105 | 97 | 92 | 95 | 86 |
| **HU52** | 57 | 92 | 89 | 102 | 97 | 99 | 96 | 96 | 77 | 55 | 90 | 92 | 99 | 99 | 58 | 95 | 91 | 94 | 78 | 80 | 96 | 98 | 94 | 95 | 89 | 88 | 86 | 89 | 99 | 97 | 101 |
| **HU53** | 12 | 10 | 30 | 96 | 100 | 81 | 17 | 98 | 16 | 98 | 94 | 91 | 36 | 90 | 81 | 96 | 56 | 86 | 88 | 81 | 37 | 98 | 101 | 98 | 113 | 111 | 101 | 103 | 90 | 85 | 81 |
| **HU54** | 18 | 42 | 55 | 110 | 102 | 71 | 50 | 76 | 67 | 93 | 87 | 83 | 86 | 82 | 69 | 48 | 87 | 73 | 88 | 90 | 77 | 102 | 95 | 81 | 83 | 104 | 88 | 86 | 98 | 90 | 100 |
| **HU55** | 64 | 58 | 86 | 107 | 96 | 94 | 77 | 74 | 66 | 86 | 86 | 71 | 55 | 83 | 71 | 92 | 92 | 74 | 88 | 83 | 9 | 102 | 95 | 84 | 116 | 101 | 103 | 79 | 90 | 90 | 74 |
| **HU56** | 87 | 87 | 86 | 94 | 95 | 92 | 92 | 91 | 89 | 80 | 71 | 75 | 107 | 96 | 74 | 88 | 83 | 69 | 90 | 86 | 75 | 95 | 101 | 79 | 115 | 118 | 111 | 85 | 106 | 101 | 100 |
| **HU57** | 96 | 102 | 92 | 92 | 91 | 86 | 93 | 90 | 94 | 89 | 86 | 91 | 114 | 114 | 118 | 90 | 87 | 96 | 83 | 78 | 53 | 103 | 99 | 85 | 114 | 108 | 96 | 87 | 93 | 89 | 79 |
| **HU58** | 30 | 92 | 88 | 94 | 91 | 83 | 90 | 93 | 25 | 99 | 100 | 96 | 95 | 14 | 87 | 95 | 94 | 91 | 86 | 84 | 69 | 22 | 81 | 70 | 87 | 74 | 77 | 38 | 109 | 98 | 105 |
| **HU59** | 21 | 19 | 51 | 93 | 89 | 78 | 20 | 92 | 21 | 102 | 101 | 91 | 49 | 93 | 84 | 101 | 95 | 89 | 80 | 76 | 43 | 105 | 105 | 87 | 121 | 107 | 96 | 92 | 92 | 82 | 72 |
| **HU60** | 12 | 93 | 73 | 92 | 90 | 74 | 92 | 90 | 71 | 92 | 90 | 86 | 95 | 93 | 90 | 19 | 91 | 89 | 93 | 81 | 59 | 89 | 78 | 62 | 85 | 79 | 61 | 60 | 91 | 77 | 72 |
| **HU61** | 17 | 22 | 21 | 44 | 90 | 27 | 24 | 88 | 77 | 96 | 101 | 45 | 17 | 96 | 77 | 93 | 84 | 33 | 94 | 84 | 68 | 97 | 91 | 80 | 105 | 98 | 94 | 83 | 91 | 87 | 73 |
| **HU62** | 97 | 91 | 91 | 106 | 105 | 73 | 94 | 90 | 74 | 97 | 93 | 84 | 94 | 90 | 81 | 99 | 92 | 81 | 87 | 89 | 76 | 98 | 105 | 71 | 112 | 117 | 113 | 89 | 100 | 98 | 103 |
| **HU63** | 29 | 36 | 29 | 40 | 103 | 77 | 40 | 72 | 83 | 102 | 91 | 86 | 91 | 94 | 77 | 98 | 95 | 74 | 95 | 91 | 82 | 103 | 101 | 89 | 96 | 91 | 72 | 76 | 101 | 99 | 104 |
| **HU64** | 92 | 95 | 87 | 92 | 86 | 82 | 94 | 87 | 78 | 98 | 100 | 105 | 97 | 96 | 92 | 94 | 95 | 96 | 84 | 96 | 91 | 88 | 96 | 97 | 97 | 93 | 100 | 109 | 103 | 105 | 105 |
| **HU65** | 20 | 16 | 64 | 31 | 31 | 65 | 90 | 3 | 36 | 33 | 93 | 88 | 24 | 91 | 84 | 91 | 24 | 28 | 33 | 33 | 25 | 101 | 54 | 64 | 66 | 104 | 84 | 65 | 36 | 72 | 54 |
| **HU66** | 93 | 93 | 88 | 86 | 83 | 75 | 103 | 97 | 104 | 92 | 93 | 87 | 89 | 91 | 77 | 92 | 94 | 84 | 82 | 76 | 41 | 100 | 87 | 69 | 97 | 98 | 86 | 73 | 95 | 81 | 80 |
| **HU67** | 25 | 24 | 37 | 87 | 85 | 65 | 30 | 87 | 69 | 92 | 92 | 90 | 101 | 91 | 98 | 95 | 93 | 94 | 93 | 104 | 101 | 98 | 96 | 102 | 99 | 97 | 104 | 108 | 98 | 99 | 100 |
| **HU68** | 90 | 94 | 84 | 92 | 91 | 83 | 95 | 91 | 67 | 86 | 83 | 24 | 39 | 105 | 88 | 97 | 37 | 42 | 87 | 82 | 73 | 94 | 100 | 75 | 127 | 117 | 116 | 81 | 120 | 105 | 105 |
| **HU69** | 28 | 22 | 29 | 37 | 91 | 68 | 30 | 77 | 63 | 89 | 90 | 86 | 31 | 87 | 86 | 93 | 31 | 34 | 90 | 73 | 42 | 93 | 72 | 77 | 105 | 92 | 90 | 74 | 90 | 86 | 73 |
| **HU70** | 12 | 8 | 39 | 110 | 108 | 75 | 21 | 89 | 17 | 91 | 96 | 84 | 33 | 87 | 72 | 95 | 77 | 75 | 90 | 85 | 34 | 90 | 90 | 66 | 99 | 97 | 93 | 74 | 102 | 95 | 114 |
| **HU71** | 85 | 80 | 84 | 120 | 100 | 100 | 92 | 80 | 83 | 93 | 90 | 84 | 99 | 105 | 80 | 93 | 89 | 82 | 86 | 76 | 73 | 94 | 85 | 85 | 104 | 95 | 84 | 84 | 89 | 80 | 80 |
| **HU72** | 79 | 61 | 55 | 73 | 59 | 53 | 72 | 59 | 53 | 93 | 93 | 95 | 98 | 99 | 92 | 95 | 93 | 89 | 94 | 83 | 78 | 98 | 80 | 74 | 91 | 89 | 80 | 75 | 98 | 91 | 84 |
| **HU73** | 24 | 20 | 60 | 86 | 74 | 60 | 91 | 78 | 25 | 93 | 89 | 73 | 19 | 39 | 69 | 87 | 87 | 72 | 88 | 80 | 60 | 69 | 92 | 77 | 114 | 102 | 97 | 78 | 91 | 87 | 68 |
| **HU74** | 95 | 88 | 88 | 91 | 86 | 76 | 89 | 92 | 82 | 92 | 87 | 87 | 104 | 99 | 78 | 90 | 88 | 75 | 86 | 78 | 41 | 97 | 89 | 65 | 117 | 111 | 95 | 70 | 100 | 87 | 81 |
| **HU75** | 28 | 23 | 30 | 67 | 86 | 79 | 31 | 88 | 80 | 91 | 90 | 35 | 31 | 63 | 69 | 87 | 27 | 14 | 86 | 82 | 48 | 101 | 100 | 99 | 106 | 104 | 101 | 33 | 96 | 93 | 84 |
| **HU76** | 85 | 85 | 75 | 82 | 83 | 66 | 87 | 83 | 66 | 103 | 90 | 76 | 110 | 91 | 77 | 91 | 84 | 69 | 87 | 88 | 73 | 93 | 92 | 70 | 96 | 101 | 90 | 78 | 105 | 101 | 117 |
| **HU77** | 20 | 6 | 73 | 93 | 93 | 63 | 55 | 84 | 64 | 83 | 78 | 87 | 24 | 108 | 113 | 87 | 79 | 96 | 82 | 77 | 76 | 106 | 101 | 95 | 121 | 113 | 103 | 105 | 99 | 94 | 84 |
| **HU78** | 5 | 10 | 19 | 116 | 117 | 83 | 19 | 88 | 16 | 75 | 75 | 86 | 18 | 72 | 104 | 78 | 73 | 84 | 83 | 80 | 30 | 102 | 91 | 81 | 98 | 100 | 95 | 74 | 96 | 83 | 94 |
| **HU79** | 70 | 65 | 84 | 115 | 115 | 27 | 72 | 66 | 85 | 83 | 76 | 74 | 96 | 92 | 86 | 12 | 97 | 83 | 88 | 79 | 79 | 101 | 100 | 14 | 17 | 83 | 101 | 4 | 96 | 90 | 78 |
| **HU80** | 31 | 47 | 81 | 91 | 83 | 82 | 94 | 88 | 22 | 94 | 96 | 79 | 93 | 65 | 81 | 82 | 88 | 82 | 93 | 89 | 56 | 27 | 101 | 80 | 92 | 95 | 81 | 71 | 103 | 100 | 100 |
| **HU81** | 98 | 102 | 84 | 90 | 88 | 19 | 98 | 96 | 83 | 89 | 92 | 87 | 94 | 90 | 82 | 96 | 94 | 85 | 87 | 83 | 78 | 103 | 98 | 20 | 110 | 105 | 91 | 86 | 92 | 87 | 74 |
| **HU82** | 97 | 39 | 83 | 89 | 80 | 77 | 90 | 9 | 73 | 90 | 88 | 94 | 111 | 103 | 108 | 87 | 86 | 90 | 87 | 87 | 75 | 90 | 75 | 72 | 89 | 88 | 79 | 71 | 100 | 100 | 107 |
| **HU83** | 94 | 94 | 82 | 89 | 17 | 73 | 92 | 92 | 80 | 12 | 91 | 79 | 89 | 91 | 79 | 95 | 96 | 76 | 8 | 7 | 78 | 100 | 104 | 90 | 115 | 105 | 101 | 93 | 91 | 9 | 77 |

| | | | | | | | | | | | | | | | | | | | | | | | | | | | | | | | |
|---|---|---|---|---|---|---|---|---|---|---|---|---|---|---|---|---|---|---|---|---|---|---|---|---|---|---|---|---|---|---|---|
| **HU84** | 5 | 6 | 37 | 84 | 84 | 74 | 18 | 87 | 19 | 92 | 99 | 84 | 32 | 90 | 76 | 93 | 87 | 73 | 95 | 85 | 22 | 97 | 106 | 69 | 100 | 108 | 99 | 69 | 113 | 112 | 121 |
| **HU85** | 14 | 18 | 59 | 61 | 93 | 30 | 56 | 95 | 93 | 94 | 92 | 22 | 24 | 59 | 82 | 15 | 74 | 67 | 91 | 89 | 70 | 107 | 110 | 81 | 24 | 60 | 50 | 90 | 108 | 98 | 110 |
| **HU93** | 89 | 86 | 74 | 88 | 80 | 68 | 94 | 84 | 75 | 96 | 96 | 96 | 95 | 95 | 86 | 98 | 96 | 93 | 101 | 98 | 93 | 97 | 101 | 89 | 104 | 99 | 97 | 93 | 89 | 94 | 69 |
| **HU94** | 71 | 33 | 57 | 86 | 82 | 70 | 33 | 86 | 76 | 93 | 98 | 95 | 36 | 87 | 78 | 95 | 93 | 78 | 93 | 90 | 84 | 98 | 109 | 93 | 115 | 123 | 117 | 104 | 103 | 105 | 114 |
| **HU95** | 20 | 89 | 79 | 98 | 96 | 71 | 87 | 79 | 71 | 97 | 97 | 82 | 99 | 104 | 70 | 95 | 86 | 69 | 84 | 81 | 61 | 103 | 100 | 87 | 112 | 109 | 104 | 88 | 99 | 96 | 77 |
| **HU96** | 22 | 47 | 83 | 110 | 106 | 84 | 89 | 92 | 78 | 94 | 80 | 88 | 103 | 108 | 87 | 97 | 90 | 77 | 87 | 85 | 70 | 98 | 95 | 74 | 100 | 99 | 99 | 83 | 109 | 98 | 119 |
| **HU97** | 93 | 77 | 80 | 104 | 97 | 104 | 97 | 78 | 83 | 92 | 85 | 93 | 125 | 114 | 121 | 90 | 84 | 101 | 86 | 88 | 76 | 93 | 90 | 71 | 93 | 87 | 78 | 74 | 99 | 98 | 105 |
| **HB01** | 78 | 52 | 85 | 115 | 102 | 117 | 80 | 29 | 87 | 107 | 124 | 114 | 38 | 101 | 91 | 98 | 110 | 98 | 86 | 74 | 56 | 100 | 31 | 82 | 113 | 100 | 90 | 81 | 97 | 85 | 84 |
| **HB02** | 30 | 90 | 69 | 93 | 86 | 68 | 94 | 87 | 63 | 86 | 65 | 57 | 113 | 97 | 65 | 92 | 73 | 59 | 102 | 96 | 78 | 99 | 98 | 78 | 92 | 96 | 82 | 72 | 93 | 88 | 89 |
| **HB03** | 5 | 4 | 9 | 89 | 83 | 67 | 6 | 85 | 71 | 102 | 93 | 80 | 5 | 102 | 72 | 88 | 83 | 61 | 85 | 82 | 80 | 107 | 103 | 89 | 115 | 112 | 106 | 92 | 93 | 93 | 81 |
| **HB05** | 20 | 91 | 91 | 101 | 96 | 72 | 88 | 77 | 5 | 90 | 88 | 87 | 92 | 27 | 86 | 96 | 89 | 89 | 83 | 80 | 69 | 99 | 87 | 47 | 112 | 98 | 88 | 19 | 109 | 100 | 102 |
| **HB06** | 17 | 81 | 89 | 101 | 84 | 81 | 94 | 88 | 80 | 94 | 94 | 103 | 92 | 98 | 95 | 93 | 94 | 102 | 107 | 97 | 79 | 96 | 87 | 76 | 93 | 92 | 87 | 77 | 96 | 91 | 58 |
| **HB07** | 95 | 92 | 101 | 136 | 134 | 86 | 94 | 91 | 101 | 98 | 95 | 93 | 94 | 90 | 86 | 93 | 96 | 86 | 83 | 81 | 79 | 106 | 101 | 69 | 116 | 90 | 103 | 98 | 97 | 94 | 87 |
| **HB08** | 90 | 59 | 73 | 84 | 76 | 70 | 93 | 23 | 75 | 97 | 89 | 82 | 61 | 76 | 86 | 82 | 87 | 78 | 87 | 85 | 65 | 112 | 30 | 92 | 100 | 115 | 107 | 90 | 103 | 93 | 107 |
| **HB09** | 26 | 100 | 98 | 88 | 82 | 80 | 94 | 83 | 91 | 97 | 94 | 94 | 92 | 87 | 78 | 89 | 86 | 77 | 85 | 81 | 75 | 98 | 91 | 81 | 108 | 102 | 94 | 82 | 91 | 88 | 73 |
| **HB10** | 6 | 7 | 23 | 80 | 78 | 67 | 23 | 86 | 73 | 97 | 83 | 75 | 89 | 20 | 77 | 90 | 82 | 67 | 80 | 82 | 46 | 99 | 98 | 70 | 106 | 109 | 106 | 72 | 93 | 101 | 103 |
| **HB11** | 20 | 34 | 40 | 53 | 93 | 40 | 40 | 94 | 6 | 88 | 85 | 91 | 29 | 106 | 100 | 99 | 27 | 88 | 102 | 85 | 17 | 95 | 93 | 35 | 99 | 63 | 85 | 26 | 99 | 91 | 82 |
| **HB12** | 7 | 17 | 69 | 89 | 91 | 76 | 31 | 97 | 79 | 105 | 98 | 124 | 150 | 167 | 180 | 107 | 104 | 125 | 95 | 93 | 23 | 97 | 102 | 74 | 97 | 101 | 84 | 83 | 105 | 88 | 138 |
| **HB13** | 26 | 22 | 84 | 100 | 102 | 72 | 96 | 89 | 19 | 90 | 87 | 86 | 58 | 22 | 93 | 95 | 95 | 91 | 84 | 86 | 81 | 95 | 109 | 86 | 114 | 107 | 108 | 92 | 88 | 90 | 74 |
| **HB14** | 32 | 53 | 83 | 112 | 105 | 86 | 86 | 82 | 15 | 98 | 97 | 82 | 43 | 19 | 83 | 97 | 97 | 93 | 81 | 79 | 78 | 20 | 106 | 93 | 112 | 110 | 105 | 90 | 90 | 87 | 73 |
| **HB15** | 87 | 86 | 88 | 86 | 79 | 77 | 95 | 86 | 82 | 95 | 95 | 96 | 61 | 96 | 88 | 101 | 96 | 93 | 89 | 92 | 84 | 95 | 99 | 89 | 121 | 126 | 133 | 124 | 100 | 99 | 107 |
| **HB16** | 87 | 91 | 71 | 88 | 81 | 66 | 94 | 85 | 73 | 94 | 88 | 84 | 91 | 89 | 87 | 97 | 66 | 84 | 90 | 95 | 18 | 103 | 99 | 81 | 99 | 106 | 96 | 89 | 96 | 92 | 102 |
| **HB17** | 74 | 94 | 76 | 90 | 87 | 70 | 98 | 89 | 77 | 96 | 99 | 88 | 85 | 88 | 76 | 86 | 29 | 78 | 89 | 82 | 25 | 100 | 96 | 84 | 117 | 104 | 104 | 87 | 89 | 90 | 72 |
| **HB18** | 88 | 84 | 74 | 93 | 82 | 73 | 94 | 87 | 75 | 93 | 95 | 87 | 93 | 96 | 83 | 95 | 95 | 86 | 80 | 84 | 77 | 97 | 101 | 80 | 111 | 115 | 111 | 86 | 109 | 106 | 107 |
| **HB19** | 92 | 92 | 81 | 101 | 101 | 32 | 98 | 93 | 79 | 96 | 91 | 86 | 100 | 96 | 91 | 32 | 95 | 89 | 89 | 88 | 85 | 20 | 89 | 12 | 34 | 96 | 30 | 84 | 95 | 88 | 81 |
| **HB20** | 25 | 35 | 42 | 45 | 99 | 48 | 36 | 94 | 76 | 90 | 94 | 51 | 40 | 133 | 141 | 98 | 86 | 53 | 88 | 83 | 80 | 99 | 96 | 88 | 101 | 101 | 98 | 88 | 98 | 95 | 85 |
| **HB21** | 85 | 81 | 82 | 105 | 92 | 69 | 85 | 85 | 67 | 82 | 83 | 84 | 94 | 92 | 91 | 100 | 88 | 89 | 80 | 82 | 82 | 100 | 98 | 96 | 108 | 103 | 99 | 94 | 100 | 96 | 90 |
| **HB22** | 25 | 29 | 33 | 67 | 86 | 81 | 31 | 64 | 65 | 94 | 92 | 31 | 26 | 87 | 77 | 94 | 34 | 71 | 89 | 87 | 67 | 98 | 106 | 75 | 110 | 110 | 103 | 23 | 102 | 95 | 95 |
| **HB23** | 20 | 14 | 19 | 66 | 68 | 61 | 41 | 67 | 21 | 88 | 80 | 78 | 17 | 23 | 67 | 14 | 82 | 69 | 87 | 84 | 27 | 103 | 96 | 83 | 19 | 104 | 38 | 90 | 78 | 92 | 76 |
| **HB24** | 80 | 22 | 70 | 86 | 67 | 64 | 88 | 67 | 65 | 98 | 96 | 102 | 42 | 96 | 97 | 97 | 97 | 100 | 99 | 95 | 100 | 99 | 52 | 101 | 109 | 108 | 106 | 100 | 96 | 85 | 90 |
| **HB25** | 92 | 92 | 93 | 101 | 99 | 100 | 97 | 96 | 93 | 93 | 91 | 94 | 103 | 100 | 100 | 95 | 95 | 95 | 98 | 99 | 95 | 98 | 96 | 96 | 98 | 94 | 93 | 94 | 98 | 95 | 98 |
| **HB26** | 82 | 95 | 82 | 106 | 97 | 82 | 100 | 94 | 22 | 105 | 96 | 80 | 92 | 93 | 71 | 97 | 97 | 69 | 82 | 80 | 77 | 88 | 116 | 97 | 121 | 111 | 110 | 104 | 99 | 95 | 86 |
| **HB27** | 86 | 82 | 76 | 91 | 81 | 67 | 88 | 84 | 78 | 88 | 81 | 73 | 112 | 100 | 71 | 95 | 83 | 71 | 99 | 90 | 79 | 97 | 91 | 84 | 91 | 94 | 83 | 9 | 103 | 98 | 95 |
| **HB28** | 67 | 96 | 83 | 92 | 98 | 76 | 91 | 88 | 67 | 84 | 73 | 88 | 100 | 99 | 108 | 75 | 73 | 89 | 87 | 74 | 71 | 102 | 95 | 83 | 111 | 108 | 99 | 85 | 97 | 95 | 76 |
| **HB29** | 61 | 58 | 50 | 106 | 66 | 50 | 88 | 58 | 47 | 89 | 86 | 86 | 91 | 89 | 89 | 95 | 92 | 90 | 90 | 88 | 75 | 96 | 109 | 84 | 131 | 122 | 112 | 78 | 120 | 106 | 107 |
| **HB30** | 17 | 17 | 8 | 46 | 90 | 80 | 11 | 77 | 75 | 95 | 93 | 44 | 11 | 93 | 83 | 25 | 81 | 16 | 83 | 77 | 68 | 98 | 91 | 81 | 45 | 97 | 50 | 79 | 87 | 84 | 70 |
| **HB35** | 14 | 17 | 50 | 54 | 76 | 22 | 58 | 78 | 70 | 97 | 87 | 21 | 32 | 57 | 81 | 17 | 56 | 65 | 89 | 86 | 66 | 103 | 106 | 66 | 23 | 94 | 47 | 81 | 100 | 90 | 99 |

**Table S2: Prediction scores generated for each strain from the training strain set and the difference in score from the observed data (Table S1). Pred. – prediction score, Diff. – the score difference (Prediction – Observed).**

| | A | | A01 | | AB19 | | ALD1 | | ALD2 | | ALDi98 | | BAT1 | | BO1 | | BOW | | CHAP1 | | D3 | | D4 | | D11AU | | E | | E4 | |
|---|---|---|---|---|---|---|---|---|---|---|---|---|---|---|---|---|---|---|---|---|---|---|---|---|---|---|---|---|---|---|
| Isolate | Pred. | Diff. | Pred. | Diff. | Pred. | Diff | Pred. | Diff. | Pred. | Diff. | Pred. | Diff. | Pred. | Diff. | Pred. | Diff. | Pred. | Diff | Pred. | Diff. | Pred. | Diff. | Pred. | Diff. | Pred. | Diff. | Pred. | Diff. | Pred. | Diff |
| CAN1 | 91 | -3 | 94 | -2 | 81 | -12 | 89 | -5 | 93 | -7 | 95 | 1 | 85 | -15 | 91 | -9 | 95 | -5 | 92 | -6 | 90 | -10 | 82 | -18 | 92 | 0 | 85 | -14 | 90 | 3 |
| CAN10 | 91 | -2 | 93 | -7 | 84 | -15 | 90 | 0 | 94 | 7 | 95 | -5 | 82 | -18 | 93 | 4 | 96 | -3 | 73 | -28 | 84 | -10 | 77 | -5 | 86 | -14 | 70 | 45 | 96 | 5 |
| CAN100 | 81 | -7 | 93 | -2 | 81 | -14 | 85 | -2 | 74 | -18 | 95 | 3 | 91 | 10 | 71 | 15 | 92 | 27 | 66 | -34 | 82 | -4 | 87 | 8 | 85 | -14 | 86 | 21 | 86 | -12 |
| CAN101 | 25 | 19 | 94 | -7 | 84 | 3 | 88 | 1 | 94 | 4 | 95 | -5 | 91 | 9 | 91 | 5 | 68 | -23 | 78 | -13 | 89 | 4 | 66 | -3 | 93 | -7 | 31 | 15 | 89 | -11 |
| CAN102 | 92 | 6 | 93 | -7 | 84 | -16 | 94 | 5 | 93 | -7 | 95 | -4 | 85 | -3 | 92 | 6 | 95 | 1 | 43 | 9 | 93 | 0 | 93 | 0 | 82 | -18 | 88 | -3 | 97 | -3 |
| CAN103 | 92 | 7 | 94 | -6 | 85 | -14 | 90 | 18 | 38 | 5 | 95 | -5 | 42 | 10 | 80 | -7 | 96 | 6 | 35 | 1 | 60 | 32 | 77 | -10 | 92 | -4 | 88 | 1 | 39 | 8 |
| CAN104 | 92 | -2 | 94 | -4 | 84 | -9 | 92 | -2 | 91 | -7 | 95 | -2 | 70 | -16 | 91 | -6 | 95 | -4 | 21 | 15 | 91 | -5 | 80 | -10 | 82 | -18 | 88 | -9 | 95 | -3 |
| CAN105 | 88 | 0 | 94 | -6 | 61 | -6 | 88 | -1 | 94 | 6 | 95 | 1 | 38 | 20 | 64 | 13 | 86 | 20 | 30 | 20 | 87 | 0 | 33 | 7 | 87 | 9 | 60 | 18 | 33 | 22 |
| CAN106 | 77 | 23 | 94 | -4 | 68 | 20 | 87 | 1 | 67 | 40 | 95 | -2 | 67 | 42 | 38 | -6 | 92 | 44 | 36 | 20 | 78 | -3 | 52 | 24 | 88 | 30 | 78 | 44 | 52 | 29 |
| CAN107 | 91 | 3 | 93 | -7 | 87 | -13 | 91 | -2 | 61 | -37 | 95 | -5 | 63 | -27 | 92 | 2 | 96 | 6 | 78 | 40 | 85 | -8 | 70 | -11 | 83 | 5 | 88 | 3 | 71 | -22 |
| CAN108 | 90 | -7 | 94 | 10 | 71 | -7 | 89 | -6 | 92 | 3 | 95 | 2 | 77 | -8 | 91 | -9 | 96 | -1 | 88 | -8 | 86 | -6 | 78 | 0 | 91 | 13 | 85 | -14 | 95 | 2 |
| CAN109 | 91 | -9 | 93 | -3 | 81 | -19 | 88 | -5 | 36 | -47 | 80 | -18 | 72 | -11 | 88 | -12 | 95 | -5 | 45 | 7 | 91 | 5 | 72 | -28 | 38 | 14 | 85 | -11 | 77 | -23 |
| CAN11 | 91 | -1 | 94 | 4 | 80 | -4 | 92 | 5 | 94 | -1 | 95 | 6 | 86 | -8 | 93 | -7 | 95 | -2 | 96 | -4 | 90 | -7 | 81 | -2 | 88 | -12 | 79 | -14 | 92 | -2 |
| CAN110 | 93 | 9 | 94 | -6 | 89 | 2 | 94 | 16 | 86 | 34 | 95 | -1 | 88 | 11 | 93 | -7 | 96 | 4 | 85 | 66 | 85 | -5 | 46 | 16 | 95 | -1 | 85 | -3 | 92 | -8 |
| CAN111 | 93 | 0 | 94 | -3 | 79 | -21 | 88 | 1 | 64 | 30 | 80 | -18 | 74 | 8 | 89 | 3 | 95 | -3 | 19 | -3 | 91 | 4 | 71 | -7 | 37 | -56 | 83 | -17 | 36 | 9 |
| CAN112 | 91 | -1 | 94 | -6 | 81 | -19 | 91 | -3 | 94 | 3 | 95 | -5 | 87 | -8 | 90 | 13 | 95 | -5 | 56 | 23 | 94 | 3 | 85 | -15 | 82 | -17 | 89 | -9 | 98 | 0 |
| CAN113 | 27 | 8 | 94 | -6 | 76 | -24 | 86 | 2 | 58 | -17 | 95 | -5 | 31 | 13 | 72 | -22 | 34 | 15 | 19 | 8 | 86 | 11 | 36 | 0 | 81 | -5 | 26 | -7 | 33 | 10 |
| CAN114 | 52 | 14 | 93 | -7 | 78 | 30 | 86 | 16 | 67 | 31 | 95 | -3 | 33 | -66 | 68 | 21 | 53 | 21 | 31 | 20 | 86 | 22 | 55 | -30 | 85 | -6 | 26 | -5 | 42 | 11 |
| CAN115 | 68 | -23 | 93 | -7 | 79 | -13 | 89 | -1 | 80 | -15 | 95 | -5 | 59 | -33 | 57 | -38 | 93 | -4 | 61 | -26 | 86 | -4 | 57 | 4 | 73 | -10 | 69 | -15 | 49 | -48 |
| CAN116 | 91 | -9 | 93 | -7 | 84 | 3 | 84 | -16 | 77 | 3 | 95 | -1 | 89 | 16 | 93 | -1 | 96 | -4 | 60 | -34 | 83 | -14 | 54 | 27 | 95 | -4 | 85 | -5 | 92 | 1 |
| CAN117 | 86 | 63 | 94 | 75 | 76 | 46 | 90 | 18 | 86 | 44 | 95 | 58 | 89 | 82 | 92 | 64 | 97 | 26 | 69 | 53 | 90 | 22 | 43 | 31 | 89 | -11 | 89 | 35 | 82 | 64 |
| CAN118 | 92 | -7 | 93 | -7 | 77 | -17 | 71 | -16 | 93 | 1 | 95 | -5 | 84 | -16 | 91 | -9 | 92 | 0 | 87 | -10 | 92 | -8 | 81 | 5 | 79 | -21 | 88 | 1 | 85 | -15 |
| CAN119 | 74 | -15 | 64 | 27 | 77 | 21 | 86 | 16 | 69 | -1 | 79 | -2 | 67 | 23 | 89 | 47 | 88 | 6 | 58 | 28 | 86 | 29 | 54 | 30 | 21 | 0 | 84 | 10 | 59 | 20 |
| CAN12 | 90 | -6 | 94 | -6 | 79 | -3 | 91 | -1 | 94 | -6 | 95 | -1 | 94 | -6 | 89 | -11 | 96 | 5 | 93 | 2 | 86 | -3 | 79 | 7 | 91 | 6 | 91 | 25 | 92 | -5 |
| CAN120 | 27 | 8 | 94 | -5 | 68 | 0 | 86 | -9 | 93 | -2 | 95 | -2 | 70 | -7 | 71 | 12 | 46 | 37 | 57 | -34 | 85 | 3 | 44 | -22 | 78 | -8 | 37 | 17 | 66 | -26 |
| CAN121 | 88 | -6 | 94 | -6 | 71 | -4 | 89 | 1 | 61 | -31 | 95 | 0 | 66 | -22 | 53 | 16 | 87 | -6 | 78 | -12 | 85 | -8 | 59 | -17 | 87 | 20 | 65 | -20 | 83 | -9 |
| CAN122 | 92 | -3 | 94 | -6 | 85 | 0 | 89 | -2 | 92 | -4 | 95 | 3 | 86 | 7 | 93 | -5 | 95 | -5 | 84 | -11 | 89 | -6 | 49 | 6 | 90 | 18 | 90 | -5 | 91 | 5 |
| CAN123 | 91 | 1 | 93 | -7 | 86 | 21 | 91 | 1 | 94 | 5 | 95 | -1 | 87 | 7 | 91 | -1 | 96 | 3 | 94 | 2 | 91 | 1 | 88 | 27 | 72 | 1 | 88 | 8 | 95 | 6 |
| CAN124 | 90 | 7 | 94 | 15 | 77 | 11 | 87 | 13 | 66 | -15 | 82 | 51 | 76 | 17 | 91 | -2 | 92 | -8 | 41 | -2 | 78 | 3 | 77 | 11 | 23 | 3 | 91 | 5 | 69 | -22 |
| CAN125 | 78 | 12 | 94 | -6 | 73 | -14 | 86 | 8 | 87 | 0 | 95 | -1 | 76 | -8 | 80 | -19 | 92 | -4 | 57 | -33 | 83 | 6 | 74 | -10 | 84 | -1 | 88 | -12 | 71 | -27 |
| CAN126 | 82 | 0 | 93 | -5 | 67 | 32 | 90 | 12 | 94 | 7 | 95 | 1 | 90 | 1 | 74 | -8 | 83 | -12 | 68 | 24 | 91 | 8 | 90 | -6 | 86 | 14 | 76 | -10 | 91 | -7 |
| CAN127 | 71 | 22 | 94 | 2 | 66 | 26 | 83 | -2 | 57 | 18 | 95 | 6 | 81 | -5 | 72 | 31 | 72 | 30 | 32 | 11 | 91 | 7 | 66 | 9 | 86 | -4 | 53 | 12 | 46 | 9 |
| CAN128 | 93 | -1 | 94 | -6 | 85 | 4 | 88 | -4 | 94 | -2 | 95 | -5 | 91 | 1 | 85 | -3 | 96 | -4 | 34 | -17 | 87 | -5 | 79 | -3 | 82 | 43 | 87 | -9 | 94 | 4 |
| CAN129 | 93 | 44 | 94 | -6 | 84 | 31 | 79 | -11 | 93 | -7 | 95 | -5 | 81 | -3 | 93 | 2 | 78 | 33 | 93 | -5 | 91 | 3 | 77 | 18 | 88 | -1 | 69 | 35 | 67 | -18 |
| CAN13 | 92 | 4 | 94 | -3 | 79 | 3 | 89 | 1 | 93 | -7 | 95 | -5 | 70 | -3 | 83 | 30 | 95 | -4 | 43 | 41 | 91 | 3 | 51 | 22 | 87 | -13 | 87 | -13 | 54 | 27 |
| CAN131 | 46 | 28 | 94 | 0 | 82 | -15 | 87 | -3 | 83 | -12 | 95 | 4 | 86 | 1 | 91 | -6 | 94 | 19 | 96 | 4 | 92 | 2 | 83 | -6 | 85 | -9 | 47 | 33 | 78 | -21 |
| CAN132 | 92 | -1 | 93 | -3 | 87 | 22 | 90 | -4 | 94 | 2 | 95 | 8 | 85 | -5 | 91 | -9 | 95 | 7 | 19 | -6 | 85 | -4 | 86 | 20 | 76 | -12 | 88 | 2 | 97 | -3 |
| CAN133 | 71 | -12 | 94 | -6 | 83 | -7 | 88 | 12 | 78 | 9 | 95 | 6 | 85 | 49 | 72 | -28 | 73 | -9 | 74 | -15 | 85 | 18 | 85 | 5 | 85 | -5 | 79 | 4 | 79 | -21 |
| CAN134 | 70 | 19 | 93 | -7 | 55 | -6 | 86 | 5 | 53 | 4 | 95 | 0 | 86 | 11 | 70 | 21 | 66 | 12 | 33 | 13 | 90 | -10 | 63 | 2 | 86 | 9 | 49 | 0 | 43 | 2 |
| CAN135 | 92 | 10 | 93 | -7 | 86 | 2 | 93 | 16 | 94 | -3 | 95 | 4 | 79 | -21 | 91 | -3 | 95 | 5 | 76 | -16 | 86 | 13 | 77 | 4 | 49 | -3 | 89 | 4 | 97 | 5 |
| CAN136 | 91 | -1 | 93 | 9 | 85 | 3 | 90 | -3 | 90 | -2 | 95 | 6 | 93 | 9 | 92 | 0 | 96 | -2 | 47 | 20 | 86 | 5 | 80 | 2 | 72 | -13 | 87 | -4 | 95 | -3 |
| CAN137 | 89 | -1 | 93 | -7 | 77 | 4 | 77 | -12 | 79 | 7 | 95 | -1 | 90 | 2 | 82 | -16 | 91 | 1 | 40 | 2 | 72 | -15 | 78 | 26 | 90 | 1 | 91 | 20 | 84 | -10 |
| CAN138 | 91 | 0 | 93 | -7 | 86 | 12 | 90 | -3 | 94 | -4 | 95 | 1 | 82 | -3 | 91 | -6 | 95 | 3 | 74 | -23 | 85 | -14 | 77 | 7 | 55 | 20 | 88 | -2 | 96 | -4 |
| CAN139 | 89 | -5 | 94 | 24 | 72 | 28 | 87 | -2 | 91 | 42 | 80 | -4 | 68 | 39 | 90 | 26 | 93 | 11 | 56 | 48 | 86 | 55 | 71 | 38 | 55 | 14 | 91 | 11 | 61 | 38 |

| | | | | | | | | | | | | | | | | | | | | | | | | | | | | | | |
|---|---|---|---|---|---|---|---|---|---|---|---|---|---|---|---|---|---|---|---|---|---|---|---|---|---|---|---|---|---|---|
| **CAN14** | 94 | -5 | 93 | -7 | 89 | -3 | 89 | 3 | 91 | -9 | 95 | -5 | 84 | -16 | 90 | -10 | 95 | -3 | 67 | -33 | 89 | 5 | 78 | 2 | 86 | -11 | 90 | 10 | 80 | -9 |
| **CAN140** | 86 | -4 | 93 | -2 | 81 | 13 | 89 | 6 | 31 | 20 | 95 | 2 | 31 | 14 | 41 | 9 | 94 | 0 | 18 | 10 | 45 | 20 | 34 | 17 | 73 | 18 | 83 | -8 | 36 | 18 |
| **CAN141** | 90 | -2 | 94 | -2 | 73 | 5 | 88 | -5 | 93 | -3 | 95 | 11 | 70 | -18 | 71 | -29 | 65 | -27 | 79 | -10 | 90 | -10 | 47 | -12 | 70 | -15 | 88 | -11 | 93 | -4 |
| **CAN142** | 91 | 16 | 93 | 11 | 77 | -2 | 89 | 18 | 82 | 26 | 95 | 1 | 90 | 2 | 83 | -17 | 92 | -4 | 38 | -6 | 80 | 4 | 63 | -22 | 88 | -12 | 87 | -14 | 91 | 11 |
| **CAN143** | 94 | 14 | 93 | -7 | 88 | -12 | 87 | 5 | 93 | -1 | 95 | -5 | 61 | -30 | 91 | -2 | 95 | 4 | 71 | -15 | 80 | -1 | 52 | 26 | 90 | 4 | 88 | 1 | 78 | -15 |
| **CAN144** | 92 | 5 | 94 | -6 | 77 | -4 | 92 | 11 | 94 | 1 | 95 | -1 | 80 | -20 | 91 | 2 | 95 | -3 | 43 | -48 | 86 | 2 | 63 | -21 | 85 | -10 | 89 | 0 | 94 | -2 |
| **CAN145** | 91 | 1 | 94 | -6 | 85 | -3 | 84 | -3 | 89 | -6 | 95 | -2 | 77 | 3 | 91 | -1 | 95 | -2 | 68 | 20 | 88 | 4 | 75 | -9 | 49 | -20 | 89 | 8 | 97 | 5 |
| **CAN146** | 91 | 3 | 94 | -6 | 89 | 13 | 88 | 0 | 42 | 14 | 95 | 1 | 47 | -23 | 79 | -16 | 96 | -4 | 34 | 9 | 68 | -25 | 79 | 1 | 85 | -6 | 88 | 0 | 61 | -24 |
| **CAN147** | 90 | 1 | 94 | -6 | 70 | -2 | 89 | 0 | 93 | -4 | 95 | 1 | 88 | -1 | 38 | -8 | 95 | 2 | 60 | -32 | 89 | -10 | 58 | -11 | 84 | -7 | 81 | -11 | 60 | -37 |
| **CAN148** | 80 | 12 | 94 | 2 | 80 | 4 | 90 | -2 | 93 | 3 | 95 | 1 | 41 | 17 | 65 | 26 | 46 | 23 | 25 | 8 | 87 | -3 | 68 | -11 | 85 | 8 | 47 | 13 | 30 | 12 |
| **CAN149** | 81 | 56 | 94 | -6 | 85 | -10 | 89 | -8 | 94 | 4 | 95 | -5 | 84 | 60 | 60 | 10 | 46 | 21 | 26 | 6 | 90 | -10 | 79 | -3 | 80 | -3 | 49 | 30 | 45 | 25 |
| **CAN15** | 92 | 0 | 93 | -7 | 84 | 4 | 88 | -1 | 73 | -27 | 95 | -5 | 82 | -18 | 93 | -7 | 96 | 3 | 59 | -26 | 85 | 1 | 48 | -32 | 96 | -4 | 84 | -10 | 89 | -11 |
| **CAN150** | 92 | 7 | 94 | -6 | 84 | -16 | 90 | 4 | 94 | 0 | 95 | -4 | 77 | -10 | 91 | 13 | 95 | 0 | 50 | -36 | 93 | 1 | 86 | -14 | 86 | -1 | 88 | -6 | 97 | 6 |
| **CAN151** | 89 | 2 | 93 | -7 | 77 | -11 | 84 | 7 | 86 | -8 | 95 | -2 | 93 | 7 | 86 | -6 | 96 | 3 | 71 | -15 | 77 | 8 | 78 | 1 | 91 | 7 | 88 | 3 | 76 | -14 |
| **CAN152** | 92 | 13 | 94 | -6 | 76 | -4 | 91 | 12 | 59 | 10 | 95 | -5 | 83 | -6 | 93 | 18 | 93 | -7 | 37 | -1 | 86 | 5 | 44 | 14 | 64 | 15 | 85 | -3 | 66 | -2 |
| **CAN153** | 32 | -65 | 93 | -7 | 69 | 32 | 87 | -7 | 64 | -35 | 95 | -5 | 90 | 2 | 93 | 9 | 81 | -11 | 73 | -23 | 84 | -9 | 77 | 5 | 85 | 22 | 34 | -47 | 69 | -26 |
| **CAN154** | 91 | -6 | 93 | -5 | 84 | -16 | 89 | -8 | 92 | -1 | 95 | -1 | 94 | -2 | 93 | 4 | 95 | -5 | 88 | -8 | 89 | -8 | 57 | -33 | 85 | 0 | 90 | -7 | 92 | 2 |
| **CAN155** | 92 | -5 | 93 | -7 | 75 | -9 | 85 | -10 | 61 | 17 | 95 | -5 | 82 | -15 | 93 | -4 | 94 | 1 | 46 | 26 | 86 | 0 | 41 | 1 | 59 | -6 | 86 | 1 | 67 | -4 |
| **CAN156** | 71 | 63 | 94 | 7 | 79 | 53 | 90 | 11 | 92 | -3 | 95 | 7 | 83 | -17 | 89 | 6 | 77 | 49 | 84 | 10 | 91 | 19 | 80 | -3 | 74 | -19 | 79 | 73 | 64 | -13 |
| **CAN157** | 92 | 2 | 94 | -6 | 77 | 12 | 87 | -2 | 60 | 9 | 95 | 12 | 89 | 25 | 93 | -4 | 95 | 11 | 39 | 5 | 86 | 7 | 45 | 16 | 61 | 2 | 87 | 8 | 73 | 17 |
| **CAN158** | 92 | -1 | 93 | -7 | 84 | -6 | 91 | 0 | 94 | 8 | 95 | -1 | 91 | 31 | 92 | 7 | 95 | 1 | 22 | 6 | 86 | 0 | 76 | -17 | 87 | 27 | 88 | -4 | 97 | -3 |
| **CAN159** | 85 | -7 | 94 | -6 | 80 | -2 | 84 | 7 | 89 | -11 | 95 | -2 | 85 | -15 | 90 | -1 | 92 | -2 | 57 | -33 | 79 | 6 | 62 | -2 | 79 | -16 | 89 | 11 | 85 | -15 |
| **CAN16** | 88 | 2 | 93 | -6 | 81 | -3 | 88 | 17 | 32 | -24 | 95 | 5 | 53 | 11 | 83 | 2 | 93 | 0 | 28 | 5 | 56 | -20 | 51 | 13 | 90 | -10 | 89 | 3 | 40 | 3 |
| **CAN160** | 89 | 1 | 94 | 15 | 76 | -2 | 85 | 0 | 82 | 44 | 81 | -7 | 71 | -2 | 91 | 7 | 92 | -8 | 49 | 27 | 86 | -10 | 68 | -12 | 20 | -11 | 91 | 10 | 85 | 16 |
| **CAN161** | 92 | 0 | 93 | 0 | 87 | 7 | 89 | -1 | 94 | 3 | 95 | 9 | 85 | -5 | 91 | 0 | 95 | 4 | 58 | -32 | 93 | 3 | 92 | 15 | 62 | -23 | 88 | -5 | 97 | -2 |
| **CAN162** | 89 | -5 | 93 | -7 | 76 | 18 | 83 | -8 | 61 | 33 | 80 | -13 | 83 | -10 | 91 | 36 | 94 | 57 | 36 | 13 | 86 | -7 | 69 | -14 | 47 | -44 | 91 | 0 | 71 | -27 |
| **CAN163** | 92 | 1 | 94 | -6 | 85 | -4 | 90 | 0 | 94 | 1 | 95 | -5 | 85 | -7 | 91 | -6 | 95 | -4 | 22 | 3 | 86 | -1 | 79 | -4 | 77 | -9 | 88 | -3 | 99 | 4 |
| **CAN164** | 25 | -4 | 94 | 2 | 77 | -2 | 83 | -6 | 69 | -12 | 95 | -5 | 28 | -36 | 59 | -5 | 27 | -66 | 23 | 7 | 84 | -9 | 29 | -46 | 83 | 2 | 26 | -7 | 26 | -31 |
| **CAN165** | 86 | 8 | 94 | -1 | 85 | -14 | 81 | -8 | 59 | -13 | 95 | -1 | 38 | 7 | 92 | -2 | 95 | -2 | 37 | 9 | 90 | -8 | 76 | -22 | 71 | 21 | 93 | 0 | 89 | 12 |
| **CAN166** | 90 | -5 | 94 | -3 | 86 | -1 | 89 | -10 | 95 | 16 | 95 | 0 | 89 | 6 | 87 | -13 | 91 | -9 | 21 | 4 | 95 | -3 | 79 | -5 | 78 | -7 | 88 | 3 | 95 | 1 |
| **CAN167** | 30 | 25 | 94 | -6 | 72 | 2 | 90 | 12 | 67 | 46 | 95 | -4 | 81 | 72 | 85 | 50 | 42 | 29 | 78 | 64 | 88 | 16 | 60 | 53 | 78 | -8 | 30 | 17 | 84 | 71 |
| **CAN168** | 91 | -2 | 94 | -6 | 83 | -6 | 94 | 12 | 94 | -1 | 95 | -5 | 88 | -11 | 73 | -13 | 96 | -1 | 85 | -6 | 89 | -9 | 89 | -10 | 89 | -4 | 88 | -1 | 95 | -5 |
| **CAN169** | 92 | 9 | 94 | -6 | 86 | 18 | 91 | 11 | 94 | 6 | 95 | 7 | 73 | -20 | 91 | 1 | 95 | 6 | 47 | -35 | 78 | 0 | 89 | 23 | 91 | 19 | 88 | 0 | 95 | -1 |
| **CAN17** | 60 | 36 | 94 | 12 | 60 | 30 | 90 | 6 | 93 | -7 | 95 | 9 | 87 | -13 | 89 | -11 | 77 | -23 | 67 | 21 | 91 | 2 | 78 | -3 | 91 | -9 | 43 | 23 | 97 | 0 |
| **CAN170** | 88 | -3 | 94 | -6 | 80 | -2 | 87 | -6 | 35 | 23 | 95 | -2 | 32 | 18 | 90 | -3 | 96 | -4 | 44 | 12 | 86 | -9 | 46 | 15 | 36 | 17 | 88 | -3 | 40 | 2 |
| **CAN171** | 92 | -2 | 93 | 0 | 80 | -3 | 88 | -5 | 89 | -11 | 95 | 5 | 61 | -26 | 91 | -9 | 95 | -4 | 49 | -48 | 92 | -6 | 80 | 0 | 82 | -7 | 92 | -8 | 64 | -31 |
| **CAN172** | 69 | 42 | 94 | 2 | 77 | 13 | 84 | 3 | 58 | 26 | 95 | -1 | 42 | 24 | 75 | 10 | 40 | 20 | 21 | 5 | 84 | 4 | 44 | 27 | 74 | 18 | 32 | 15 | 50 | 23 |
| **CAN173** | 50 | 11 | 93 | -7 | 81 | -14 | 89 | 7 | 59 | -41 | 95 | -5 | 55 | -36 | 91 | -9 | 54 | 25 | 85 | -15 | 86 | -1 | 70 | -16 | 90 | 4 | 55 | -35 | 82 | -4 |
| **CAN174** | 35 | 8 | 94 | 3 | 76 | -7 | 86 | 1 | 90 | 13 | 95 | -1 | 40 | 17 | 74 | 6 | 33 | 9 | 21 | 1 | 85 | 2 | 44 | 19 | 83 | 2 | 30 | 9 | 32 | 7 |
| **CAN175** | 88 | -5 | 94 | -6 | 84 | 18 | 92 | 5 | 37 | 27 | 95 | -4 | 31 | 18 | 90 | -5 | 96 | 5 | 47 | 12 | 87 | 12 | 46 | 17 | 29 | -3 | 89 | 1 | 42 | 7 |
| **CAN176** | 90 | 29 | 93 | -7 | 88 | 4 | 89 | 13 | 59 | -10 | 95 | 3 | 36 | -1 | 92 | -3 | 95 | 3 | 36 | 2 | 89 | 1 | 82 | 9 | 54 | -23 | 93 | 7 | 77 | -22 |
| **CAN177** | 90 | 4 | 93 | -7 | 84 | -16 | 95 | 9 | 92 | -4 | 95 | -4 | 87 | -6 | 88 | -1 | 92 | -2 | 19 | 14 | 96 | 9 | 81 | 0 | 81 | 36 | 88 | 1 | 94 | 1 |
| **CAN178** | 91 | -2 | 93 | 4 | 86 | 9 | 91 | -1 | 93 | -1 | 95 | 0 | 91 | -2 | 79 | -15 | 94 | -6 | 51 | 26 | 89 | -2 | 79 | 2 | 90 | 0 | 89 | 7 | 92 | 2 |
| **CAN179** | 83 | -12 | 94 | -6 | 54 | -30 | 86 | -5 | 94 | 4 | 95 | -5 | 89 | -7 | 92 | -2 | 93 | -2 | 61 | 44 | 85 | 5 | 62 | -11 | 92 | 5 | 89 | 6 | 75 | -8 |
| **CAN18** | 64 | 7 | 94 | -1 | 89 | 28 | 88 | -5 | 94 | 15 | 95 | 1 | 84 | -16 | 58 | -42 | 41 | -12 | 54 | -5 | 90 | -4 | 78 | -22 | 85 | -12 | 30 | -8 | 61 | 29 |
| **CAN180** | 86 | -5 | 94 | -2 | 80 | -16 | 85 | -7 | 33 | 25 | 95 | -5 | 31 | 14 | 43 | 21 | 93 | -7 | 19 | 16 | 44 | 12 | 29 | 5 | 66 | -5 | 84 | -3 | 38 | 17 |
| **CAN181** | 90 | -1 | 93 | -7 | 86 | 4 | 90 | -6 | 94 | -4 | 95 | -5 | 90 | 2 | 91 | -9 | 95 | -5 | 92 | -8 | 91 | -5 | 66 | -7 | 88 | 3 | 81 | -4 | 97 | 6 |
| **CAN182** | 96 | 1 | 93 | -7 | 55 | 11 | 96 | 5 | 90 | 10 | 95 | 9 | 89 | -1 | 68 | -27 | 96 | 3 | 51 | 6 | 94 | -2 | 91 | 19 | 86 | -4 | 93 | 4 | 96 | -1 |
| **CAN183** | 94 | -6 | 93 | -3 | 53 | 6 | 90 | -10 | 88 | -12 | 95 | 9 | 90 | 6 | 64 | 4 | 93 | -5 | 57 | 24 | 93 | -7 | 84 | -8 | 85 | -11 | 90 | -4 | 95 | -5 |
| **CAN184** | 91 | -1 | 94 | -5 | 85 | 6 | 85 | -11 | 94 | -3 | 95 | -5 | 90 | -7 | 72 | -15 | 96 | -4 | 83 | -12 | 91 | -1 | 93 | 3 | 89 | -2 | 88 | -2 | 95 | -5 |
| **CAN185** | 89 | 6 | 93 | -7 | 80 | -7 | 87 | 9 | 94 | -2 | 95 | -5 | 84 | -5 | 83 | -11 | 92 | -6 | 74 | -20 | 85 | 8 | 68 | -2 | 73 | -9 | 93 | 12 | 66 | 15 |
| **CAN186** | 91 | -5 | 94 | 8 | 85 | 0 | 88 | -9 | 53 | -47 | 95 | -5 | 84 | -16 | 91 | -9 | 95 | -5 | 55 | -46 | 79 | -10 | 76 | -15 | 78 | -9 | 89 | -5 | 49 | -47 |
| **CAN187** | 81 | -6 | 93 | -6 | 78 | 35 | 85 | -10 | 94 | 10 | 95 | -2 | 36 | -2 | 38 | 14 | 57 | 23 | 13 | 4 | 91 | -7 | 55 | 9 | 41 | 29 | 68 | -18 | 27 | 2 |

| | | | | | | | | | | | | | | | | | | | | | | | | | | | | | | |
|---|---|---|---|---|---|---|---|---|---|---|---|---|---|---|---|---|---|---|---|---|---|---|---|---|---|---|---|---|---|---|
| **CAN188** | 80 | -5 | 93 | -7 | 67 | 15 | 91 | 7 | 91 | -1 | 95 | -5 | 36 | -3 | 39 | 17 | 43 | -1 | 16 | 7 | 88 | 8 | 53 | 8 | 38 | 26 | 73 | -1 | 26 | -1 |
| **CAN189** | 88 | -2 | 94 | -6 | 85 | -11 | 89 | 1 | 94 | 9 | 95 | 3 | 84 | -2 | 76 | 21 | 84 | -7 | 85 | -5 | 84 | 3 | 90 | 8 | 86 | 8 | 82 | -18 | 91 | 4 |
| **CAN19** | 62 | 30 | 94 | 2 | 80 | -1 | 92 | 2 | 94 | -6 | 95 | 2 | 85 | 49 | 63 | 3 | 88 | 50 | 67 | 46 | 87 | -4 | 76 | 47 | 86 | 11 | 57 | 35 | 68 | 46 |
| **CAN190** | 61 | 28 | 94 | 0 | 76 | -4 | 83 | -1 | 71 | 2 | 95 | -2 | 29 | 1 | 58 | -14 | 49 | 2 | 27 | 8 | 85 | 3 | 54 | -19 | 82 | 0 | 27 | 6 | 33 | 11 |
| **CAN191** | 93 | 3 | 94 | -2 | 82 | 1 | 89 | 4 | 93 | -1 | 81 | -5 | 83 | -5 | 93 | -7 | 95 | 10 | 68 | -24 | 83 | -2 | 68 | -17 | 44 | -46 | 88 | -3 | 66 | -34 |
| **CAN192** | 62 | -27 | 93 | -3 | 75 | -15 | 91 | 6 | 94 | 6 | 95 | -5 | 79 | -15 | 76 | 28 | 88 | 36 | 73 | 61 | 85 | 0 | 82 | -18 | 78 | -8 | 63 | 32 | 87 | -13 |
| **CAN193** | 89 | 3 | 93 | -7 | 74 | -24 | 82 | 2 | 86 | -6 | 95 | -5 | 86 | -14 | 87 | -5 | 96 | 3 | 73 | -21 | 72 | -8 | 73 | -10 | 90 | 0 | 88 | -1 | 85 | -8 |
| **CAN194** | 93 | 1 | 94 | 63 | 75 | -7 | 89 | 0 | 93 | -4 | 95 | -5 | 83 | -11 | 70 | 30 | 95 | -5 | 18 | -33 | 83 | -1 | 73 | 36 | 84 | -8 | 88 | -12 | 88 | 2 |
| **CAN195** | 90 | -3 | 93 | -7 | 83 | 8 | 90 | -2 | 88 | 52 | 95 | -1 | 83 | -4 | 91 | 3 | 92 | -1 | 55 | 30 | 86 | -11 | 70 | 4 | 81 | 2 | 86 | 2 | 91 | 54 |
| **CAN196** | 84 | -12 | 93 | 4 | 79 | 7 | 81 | -7 | 85 | -9 | 95 | 11 | 81 | -5 | 93 | -3 | 91 | -9 | 74 | -16 | 83 | 2 | 75 | 1 | 73 | -6 | 90 | -1 | 76 | -12 |
| **CAN197** | 88 | -12 | 94 | 35 | 82 | 0 | 89 | -10 | 83 | 29 | 95 | -5 | 86 | 4 | 91 | -9 | 96 | 4 | 70 | 48 | 91 | 8 | 79 | -1 | 89 | 11 | 89 | 0 | 95 | 6 |
| **CAN199** | 89 | 6 | 94 | 7 | 82 | -3 | 90 | 22 | 92 | -8 | 95 | 3 | 86 | -8 | 86 | -14 | 96 | -3 | 50 | -49 | 89 | 22 | 82 | -13 | 71 | 6 | 88 | -12 | 89 | -9 |
| **CAN2** | 92 | 6 | 93 | -7 | 84 | -11 | 90 | 8 | 24 | -65 | 95 | 3 | 56 | -44 | 89 | -6 | 95 | 5 | 22 | -54 | 80 | 1 | 70 | -19 | 79 | -21 | 85 | -10 | 27 | -12 |
| **CAN20** | 72 | -11 | 94 | -4 | 71 | 25 | 82 | -2 | 94 | -6 | 95 | 0 | 89 | -10 | 63 | 9 | 76 | -16 | 83 | -17 | 82 | -7 | 82 | -10 | 89 | 15 | 77 | -11 | 91 | 20 |
| **CAN200** | 90 | -2 | 93 | -7 | 72 | -5 | 87 | -3 | 81 | -12 | 95 | -3 | 70 | -12 | 92 | -5 | 91 | -2 | 43 | 14 | 91 | -5 | 69 | 0 | 85 | -13 | 88 | 2 | 80 | -20 |
| **CAN201** | 64 | -24 | 94 | -2 | 59 | -30 | 90 | 6 | 94 | -6 | 95 | -5 | 81 | -6 | 63 | -29 | 72 | -28 | 74 | -17 | 94 | 8 | 75 | -17 | 87 | -13 | 64 | -27 | 67 | -30 |
| **CAN202** | 93 | -2 | 93 | -1 | 84 | -16 | 89 | -6 | 94 | -4 | 95 | 3 | 84 | -16 | 91 | -5 | 90 | -7 | 53 | 21 | 86 | -2 | 67 | -26 | 82 | -18 | 84 | -15 | 94 | -6 |
| **CAN203** | 92 | 5 | 94 | -1 | 82 | 5 | 87 | 0 | 74 | -23 | 95 | 7 | 89 | -3 | 91 | -9 | 95 | -5 | 95 | 3 | 91 | 4 | 80 | 0 | 89 | -9 | 89 | -2 | 96 | 3 |
| **CAN204** | 89 | 5 | 94 | -6 | 80 | 5 | 90 | 14 | 94 | 2 | 95 | -5 | 79 | -9 | 74 | -9 | 87 | -7 | 59 | -36 | 86 | 17 | 83 | 4 | 79 | -21 | 88 | 0 | 77 | -23 |
| **CAN205** | 92 | -3 | 94 | -6 | 84 | 10 | 92 | 2 | 93 | 1 | 95 | -1 | 93 | 8 | 93 | -7 | 95 | -5 | 86 | -4 | 89 | 6 | 59 | -12 | 95 | -5 | 90 | 0 | 95 | 8 |
| **CAN21** | 82 | -1 | 94 | -6 | 72 | 42 | 85 | -4 | 93 | -7 | 95 | -5 | 82 | -18 | 93 | -7 | 75 | 47 | 93 | 1 | 86 | 4 | 80 | -8 | 88 | -7 | 79 | 44 | 71 | 27 |
| **CAN23** | 88 | -10 | 93 | -5 | 87 | -13 | 87 | -11 | 93 | 1 | 95 | -2 | 90 | 7 | 75 | 10 | 84 | -15 | 76 | -3 | 89 | -1 | 63 | -26 | 84 | 60 | 82 | -2 | 91 | -2 |
| **CAN24** | 92 | -5 | 94 | 22 | 86 | 10 | 92 | 7 | 80 | 51 | 95 | 7 | 74 | 34 | 91 | -9 | 95 | -4 | 35 | 3 | 85 | 2 | 81 | -4 | 87 | -13 | 88 | -10 | 96 | -4 |
| **CAN25** | 93 | 24 | 94 | -6 | 59 | -33 | 90 | 0 | 77 | 45 | 95 | -3 | 64 | 35 | 91 | -9 | 95 | 0 | 74 | 42 | 95 | -3 | 73 | -1 | 84 | -5 | 89 | 4 | 71 | 45 |
| **CAN26** | 89 | -2 | 93 | 11 | 86 | 4 | 89 | 3 | 93 | -7 | 95 | -5 | 86 | 4 | 93 | 1 | 61 | -39 | 82 | -13 | 85 | -4 | 77 | -19 | 86 | -6 | 89 | -11 | 88 | -6 |
| **CAN27** | 93 | 2 | 93 | -1 | 91 | 25 | 86 | -4 | 54 | 47 | 79 | -20 | 62 | 56 | 90 | 9 | 96 | -4 | 26 | 19 | 90 | -4 | 57 | 49 | 55 | 30 | 85 | -15 | 49 | 41 |
| **CAN28** | 93 | 2 | 94 | -6 | 83 | -1 | 88 | -5 | 95 | 16 | 95 | -5 | 88 | 18 | 93 | 15 | 95 | -4 | 92 | 19 | 89 | -5 | 48 | -2 | 94 | -6 | 89 | -11 | 87 | -9 |
| **CAN29** | 30 | -14 | 93 | -2 | 77 | -3 | 87 | 3 | 88 | -12 | 95 | 1 | 46 | 24 | 84 | -16 | 31 | -3 | 22 | 0 | 85 | 13 | 29 | -2 | 86 | -7 | 25 | -10 | 25 | -6 |
| **CAN3** | 88 | -4 | 94 | -3 | 86 | -3 | 87 | -4 | 94 | -1 | 95 | -5 | 89 | -2 | 61 | -32 | 83 | -17 | 69 | -29 | 84 | -1 | 75 | -7 | 88 | 14 | 80 | -1 | 67 | 14 |
| **CAN30** | 89 | -1 | 36 | 16 | 80 | 9 | 79 | -10 | 86 | -14 | 80 | -7 | 85 | -1 | 88 | -7 | 93 | -4 | 68 | -22 | 86 | 0 | 84 | 13 | 23 | 8 | 91 | -1 | 94 | 1 |
| **CAN31** | 92 | 7 | 93 | 0 | 86 | 1 | 87 | 3 | 94 | 10 | 95 | -4 | 89 | 18 | 91 | 1 | 95 | -5 | 77 | 2 | 85 | 8 | 82 | -6 | 86 | 22 | 90 | -10 | 86 | -13 |
| **CAN32** | 90 | 10 | 34 | 0 | 78 | -22 | 85 | 13 | 89 | 3 | 80 | -16 | 87 | 13 | 92 | 5 | 92 | -7 | 75 | 24 | 86 | 19 | 78 | -22 | 23 | 9 | 90 | -9 | 92 | -8 |
| **CAN33** | 25 | -5 | 94 | -6 | 77 | 5 | 85 | 5 | 67 | 34 | 95 | -4 | 32 | 17 | 74 | 1 | 34 | 20 | 20 | 11 | 86 | -2 | 28 | 0 | 87 | 30 | 42 | 22 | 37 | -49 |
| **CAN34** | 89 | 1 | 36 | 21 | 76 | -2 | 81 | -4 | 89 | 3 | 80 | -9 | 86 | -1 | 91 | -4 | 92 | -8 | 69 | 2 | 85 | -1 | 81 | -7 | 21 | -5 | 91 | -3 | 89 | -2 |
| **CAN35** | 63 | -1 | 94 | -6 | 65 | -24 | 90 | -10 | 94 | 3 | 95 | 2 | 78 | -16 | 65 | 33 | 45 | 8 | 61 | -11 | 91 | 0 | 84 | 1 | 92 | 10 | 38 | 24 | 86 | 3 |
| **CAN36** | 92 | -2 | 93 | -6 | 83 | 7 | 92 | 0 | 94 | 3 | 95 | -5 | 75 | -4 | 91 | -9 | 95 | 0 | 34 | -57 | 85 | -3 | 83 | 2 | 91 | 22 | 90 | -3 | 98 | 4 |
| **CAN37** | 90 | -7 | 94 | 0 | 62 | -33 | 89 | -2 | 93 | -7 | 95 | -5 | 90 | -10 | 91 | 0 | 96 | -2 | 92 | -8 | 86 | -2 | 74 | -24 | 87 | -13 | 88 | -12 | 90 | 2 |
| **CAN38** | 92 | -4 | 94 | -6 | 84 | -14 | 90 | -8 | 94 | -5 | 95 | -5 | 87 | -8 | 91 | 0 | 95 | 8 | 48 | 17 | 93 | -2 | 90 | -10 | 90 | 0 | 88 | -7 | 99 | 2 |
| **CAN39** | 92 | -6 | 93 | -7 | 85 | -14 | 94 | -6 | 94 | 8 | 95 | -5 | 84 | -5 | 91 | -6 | 95 | -5 | 45 | 22 | 92 | -4 | 77 | -18 | 90 | 17 | 88 | -9 | 98 | -1 |
| **CAN4** | 84 | -1 | 93 | -7 | 63 | -28 | 89 | 6 | 93 | -7 | 95 | 4 | 69 | -19 | 64 | 16 | 69 | -22 | 50 | -13 | 86 | 25 | 63 | -14 | 85 | -7 | 68 | -14 | 62 | -18 |
| **CAN40** | 89 | 2 | 94 | -4 | 85 | -11 | 87 | 8 | 75 | -24 | 95 | -4 | 79 | -7 | 91 | 0 | 96 | 0 | 77 | -15 | 86 | -2 | 78 | -20 | 54 | 36 | 89 | -11 | 76 | 53 |
| **CAN41** | 66 | 49 | 94 | 12 | 88 | -12 | 90 | 5 | 93 | -7 | 95 | -5 | 88 | -12 | 83 | 21 | 82 | 68 | 91 | 10 | 89 | -4 | 80 | -3 | 90 | -10 | 69 | 56 | 89 | 18 |
| **CAN43** | 74 | -19 | 94 | 9 | 76 | -3 | 83 | -4 | 69 | -23 | 95 | -1 | 85 | -8 | 92 | 1 | 92 | -7 | 79 | -17 | 72 | -14 | 68 | -17 | 90 | 6 | 92 | -8 | 54 | -37 |
| **CAN44** | 89 | -1 | 94 | -6 | 73 | -2 | 84 | -8 | 90 | 18 | 81 | 19 | 87 | 10 | 91 | -2 | 92 | -1 | 62 | 32 | 84 | -2 | 78 | 11 | 25 | 6 | 91 | 5 | 75 | -13 |
| **CAN45** | 27 | 6 | 94 | -2 | 76 | -8 | 89 | 3 | 70 | 0 | 95 | -3 | 31 | 1 | 66 | -5 | 35 | 11 | 22 | 2 | 85 | -5 | 27 | 4 | 91 | 4 | 27 | 6 | 24 | 7 |
| **CAN46** | 92 | -8 | 93 | -7 | 82 | -17 | 87 | 5 | 94 | 4 | 81 | 15 | 85 | -6 | 93 | 2 | 95 | -4 | 24 | 2 | 83 | 7 | 67 | -18 | 20 | -2 | 88 | -2 | 81 | -11 |
| **CAN47** | 89 | -3 | 94 | 9 | 87 | 12 | 87 | -6 | 94 | 5 | 95 | -4 | 90 | 14 | 65 | 33 | 84 | -7 | 77 | 55 | 83 | -17 | 85 | -1 | 88 | 11 | 78 | -20 | 92 | 11 |
| **CAN48** | 92 | -8 | 93 | -7 | 84 | -16 | 85 | -10 | 43 | 26 | 95 | -5 | 63 | 17 | 92 | 15 | 95 | -4 | 26 | 10 | 79 | -13 | 85 | -15 | 91 | 22 | 88 | -1 | 35 | 22 |
| **CAN49** | 41 | 17 | 94 | -5 | 79 | -20 | 89 | 9 | 69 | -8 | 95 | -4 | 41 | 24 | 79 | 4 | 32 | -14 | 22 | 5 | 84 | -9 | 32 | -19 | 85 | 32 | 52 | 7 | 42 | 16 |
| **CAN5** | 80 | 41 | 93 | -7 | 85 | -6 | 89 | -8 | 93 | -7 | 95 | -3 | 88 | -12 | 61 | -34 | 84 | -16 | 55 | -38 | 86 | 1 | 86 | 11 | 86 | 23 | 61 | 27 | 93 | 7 |
| **CAN50** | 93 | 8 | 94 | -6 | 83 | -4 | 91 | 7 | 91 | -9 | 95 | -5 | 80 | -17 | 93 | -7 | 95 | -5 | 84 | -7 | 89 | -1 | 53 | 12 | 97 | -3 | 91 | 8 | 90 | 3 |
| **CAN51** | 42 | -37 | 94 | 20 | 77 | 3 | 90 | -3 | 70 | 4 | 95 | 0 | 34 | -23 | 80 | 27 | 55 | -45 | 27 | 10 | 84 | -9 | 64 | 28 | 86 | 20 | 28 | 10 | 57 | -28 |
| **CAN52** | 91 | -4 | 94 | -3 | 85 | -13 | 89 | -4 | 92 | -8 | 95 | -1 | 80 | -20 | 91 | -9 | 95 | -5 | 79 | -20 | 84 | -11 | 71 | -29 | 55 | -10 | 87 | -13 | 96 | -3 |

| | | | | | | | | | | | | | | | | | | | | | | | | | | | | | | |
|---|---|---|---|---|---|---|---|---|---|---|---|---|---|---|---|---|---|---|---|---|---|---|---|---|---|---|---|---|---|---|
| CAN53 | 91 | -3 | 93 | -7 | 68 | -4 | 90 | -4 | 94 | 3 | 95 | -5 | 78 | -9 | 91 | -5 | 93 | -7 | 65 | -8 | 92 | -2 | 84 | 3 | 87 | 0 | 86 | -13 | 87 | -13 |
| CAN54 | 92 | -3 | 94 | -6 | 86 | 9 | 97 | 2 | 94 | -3 | 95 | -4 | 78 | -22 | 91 | -2 | 95 | 1 | 36 | -6 | 93 | 2 | 88 | 19 | 83 | -10 | 88 | 6 | 98 | -1 |
| CAN55 | 89 | 39 | 94 | -3 | 78 | 11 | 89 | -3 | 72 | -18 | 80 | -12 | 89 | -2 | 93 | 5 | 87 | -7 | 49 | 34 | 85 | -7 | 73 | 6 | 46 | -47 | 75 | 25 | 82 | -13 |
| CAN56 | 89 | -11 | 93 | -7 | 85 | -8 | 89 | -11 | 93 | -4 | 95 | -5 | 89 | 1 | 90 | -5 | 96 | 0 | 90 | -9 | 90 | -10 | 80 | 4 | 83 | -4 | 89 | 6 | 82 | -16 |
| CAN57 | 92 | -6 | 94 | 8 | 80 | 2 | 89 | 1 | 52 | 39 | 95 | 10 | 55 | -13 | 89 | -2 | 95 | -4 | 25 | 5 | 89 | -8 | 68 | -15 | 81 | -4 | 88 | -11 | 64 | -27 |
| CAN58 | 89 | 17 | 93 | -7 | 81 | -1 | 87 | 25 | 43 | -3 | 80 | -20 | 51 | -3 | 91 | -9 | 92 | -4 | 31 | -7 | 66 | 9 | 62 | 16 | 49 | 16 | 90 | 4 | 74 | 43 |
| CAN59 | 62 | -29 | 93 | -6 | 77 | -1 | 86 | -2 | 40 | 31 | 95 | -5 | 71 | 57 | 91 | -2 | 76 | -24 | 43 | 31 | 85 | -2 | 61 | -21 | 87 | 1 | 66 | -28 | 52 | 37 |
| CAN6 | 90 | -1 | 93 | -6 | 85 | -8 | 91 | 2 | 94 | 2 | 95 | -4 | 94 | -2 | 91 | -4 | 95 | -5 | 92 | -7 | 85 | -11 | 73 | -20 | 91 | 0 | 87 | -9 | 94 | -6 |
| CAN60 | 36 | 11 | 94 | -6 | 80 | 3 | 87 | -13 | 88 | -9 | 95 | 4 | 47 | 9 | 76 | -9 | 33 | 10 | 22 | 1 | 84 | -16 | 33 | 9 | 85 | -7 | 29 | 5 | 33 | 15 |
| CAN61 | 92 | 3 | 94 | 7 | 70 | 13 | 88 | 6 | 43 | 31 | 95 | 8 | 63 | 16 | 91 | 2 | 95 | 2 | 46 | 32 | 84 | 9 | 73 | 12 | 81 | -3 | 88 | -3 | 37 | 24 |
| CAN62 | 90 | 4 | 94 | -6 | 85 | 7 | 93 | 6 | 94 | 10 | 95 | -5 | 68 | 34 | 89 | 6 | 91 | -9 | 22 | 4 | 93 | 8 | 78 | 8 | 65 | 36 | 88 | -2 | 92 | -5 |
| CAN63 | 89 | 6 | 35 | 13 | 76 | -8 | 82 | 4 | 87 | -13 | 80 | -16 | 85 | -14 | 91 | -9 | 92 | -8 | 68 | -16 | 85 | 13 | 80 | -12 | 23 | 4 | 90 | -10 | 94 | 4 |
| CAN64 | 89 | -6 | 94 | -6 | 82 | -8 | 88 | -6 | 60 | 31 | 95 | -5 | 52 | 25 | 75 | 21 | 96 | 0 | 40 | 20 | 50 | 20 | 57 | -22 | 87 | -13 | 88 | 0 | 43 | 9 |
| CAN65 | 86 | -7 | 94 | -6 | 76 | -7 | 78 | -8 | 42 | -3 | 81 | 19 | 47 | -16 | 91 | 21 | 94 | 3 | 45 | 17 | 74 | -14 | 53 | -33 | 42 | -19 | 92 | -8 | 47 | -47 |
| CAN66 | 92 | 1 | 94 | -6 | 83 | 2 | 84 | -3 | 93 | -7 | 80 | 32 | 85 | -8 | 93 | -2 | 95 | -5 | 22 | -7 | 83 | 5 | 73 | 3 | 21 | 0 | 88 | 6 | 81 | -11 |
| CAN67 | 73 | -22 | 93 | -3 | 59 | -35 | 87 | -7 | 94 | 0 | 95 | -4 | 91 | -4 | 63 | 7 | 81 | -8 | 80 | -13 | 94 | 1 | 87 | -12 | 85 | 1 | 74 | -22 | 96 | -1 |
| CAN68 | 52 | -41 | 93 | -4 | 79 | -2 | 90 | -4 | 89 | -7 | 95 | 3 | 43 | -55 | 64 | -26 | 57 | -36 | 19 | -27 | 85 | -5 | 45 | -33 | 85 | -11 | 37 | -52 | 67 | -33 |
| CAN69 | 92 | 5 | 94 | 9 | 82 | 4 | 91 | 6 | 93 | -7 | 95 | 1 | 94 | -6 | 91 | -9 | 95 | -5 | 95 | -2 | 86 | -3 | 81 | -8 | 95 | -5 | 89 | -10 | 95 | -5 |
| CAN7 | 89 | -4 | 94 | 2 | 82 | -5 | 88 | -11 | 79 | -21 | 95 | -5 | 86 | -14 | 93 | -7 | 95 | -3 | 92 | -8 | 90 | -10 | 82 | -7 | 90 | -10 | 88 | -12 | 97 | -3 |
| CAN70 | 91 | 4 | 94 | -6 | 86 | 17 | 95 | 7 | 94 | -6 | 95 | 0 | 81 | -3 | 91 | -6 | 95 | 0 | 61 | -7 | 86 | -4 | 78 | 14 | 71 | 38 | 88 | 0 | 95 | -5 |
| CAN71 | 33 | 3 | 93 | -2 | 81 | 4 | 85 | 9 | 86 | -14 | 95 | -3 | 45 | 0 | 87 | 18 | 32 | 11 | 15 | -11 | 86 | 10 | 31 | 7 | 85 | -15 | 29 | 7 | 31 | 13 |
| CAN72 | 68 | -27 | 94 | -7 | 83 | 10 | 83 | -17 | 94 | -6 | 95 | -1 | 85 | -15 | 88 | -12 | 95 | 1 | 24 | -2 | 84 | -7 | 73 | 15 | 75 | -9 | 88 | 13 | 92 | 3 |
| CAN73 | 26 | 0 | 94 | 10 | 77 | -10 | 85 | -7 | 72 | 18 | 95 | 14 | 27 | -15 | 74 | -14 | 34 | 13 | 21 | -9 | 84 | -5 | 28 | 9 | 87 | -13 | 26 | -2 | 23 | 5 |
| CAN74 | 84 | -10 | 93 | -5 | 82 | -7 | 86 | -2 | 30 | -1 | 95 | -5 | 32 | -1 | 36 | -12 | 92 | -8 | 24 | 0 | 43 | 14 | 43 | -1 | 82 | -18 | 93 | -3 | 43 | 7 |
| CAN75 | 95 | -1 | 94 | -2 | 87 | -13 | 91 | -3 | 94 | -6 | 95 | 0 | 88 | -12 | 63 | -8 | 92 | -5 | 62 | 3 | 91 | -1 | 86 | -14 | 85 | -10 | 81 | -18 | 91 | -1 |
| CAN76 | 91 | -7 | 93 | -7 | 85 | 15 | 88 | -12 | 95 | 6 | 95 | 2 | 90 | 58 | 92 | 11 | 95 | 2 | 71 | 43 | 85 | -13 | 79 | 18 | 48 | -52 | 89 | 3 | 96 | -3 |
| CAN77 | 89 | 8 | 93 | -4 | 66 | 11 | 91 | 7 | 94 | -6 | 95 | -1 | 33 | -1 | 62 | -1 | 86 | 16 | 23 | -7 | 88 | 4 | 38 | 21 | 82 | -12 | 64 | 8 | 31 | 2 |
| CAN78 | 89 | 4 | 93 | -7 | 76 | -18 | 89 | 18 | 94 | -6 | 95 | -5 | 88 | -12 | 72 | -28 | 89 | -10 | 86 | -12 | 85 | 12 | 84 | -4 | 81 | -19 | 84 | -7 | 83 | -17 |
| CAN79 | 27 | 8 | 93 | -3 | 77 | 8 | 85 | -3 | 67 | -27 | 95 | -5 | 29 | 0 | 83 | -17 | 34 | 17 | 21 | -4 | 84 | -4 | 30 | 10 | 88 | -12 | 26 | -1 | 34 | 15 |
| CAN8 | 64 | 6 | 93 | -7 | 85 | -13 | 91 | 10 | 94 | -6 | 95 | -5 | 89 | 6 | 74 | 3 | 66 | -26 | 72 | -20 | 85 | -5 | 77 | -7 | 90 | -8 | 45 | -4 | 73 | -27 |
| CAN80 | 88 | -9 | 93 | -7 | 85 | -1 | 89 | -8 | 32 | -68 | 95 | -5 | 43 | -57 | 83 | 15 | 96 | 0 | 26 | 4 | 68 | -28 | 62 | 34 | 87 | -13 | 88 | 3 | 62 | -26 |
| CAN81 | 84 | -16 | 94 | 9 | 86 | 18 | 93 | -4 | 92 | -8 | 95 | -2 | 83 | -17 | 87 | -10 | 88 | -12 | 22 | 2 | 95 | -1 | 81 | 10 | 86 | -14 | 79 | -18 | 91 | 0 |
| CAN82 | 88 | -5 | 93 | -7 | 83 | 2 | 88 | 2 | 93 | -7 | 95 | -5 | 77 | -23 | 62 | -16 | 83 | -13 | 83 | -6 | 87 | 16 | 90 | 9 | 86 | -14 | 51 | -36 | 88 | 0 |
| CAN83 | 95 | -1 | 93 | -4 | 63 | 14 | 92 | -3 | 89 | -5 | 95 | -5 | 89 | -1 | 63 | 8 | 98 | 12 | 48 | -27 | 87 | -6 | 83 | -11 | 86 | -6 | 88 | -12 | 94 | -5 |
| CAN84 | 91 | -3 | 93 | -6 | 82 | -13 | 88 | 0 | 93 | -7 | 95 | -5 | 94 | -3 | 91 | -9 | 95 | 2 | 95 | -1 | 86 | 2 | 79 | -15 | 95 | -5 | 90 | -5 | 96 | -3 |
| CAN85 | 89 | -10 | 94 | -5 | 48 | 29 | 90 | -1 | 93 | -7 | 95 | -5 | 89 | -11 | 91 | -9 | 95 | -5 | 95 | 5 | 89 | 1 | 75 | -6 | 76 | -24 | 88 | -8 | 96 | -2 |
| CAN86 | 92 | -5 | 94 | -3 | 84 | -12 | 93 | -2 | 93 | 5 | 95 | -2 | 74 | -2 | 91 | 3 | 95 | -5 | 20 | 13 | 91 | -2 | 81 | -7 | 89 | 4 | 88 | -12 | 94 | -6 |
| CAN88 | 91 | 2 | 94 | 13 | 84 | 30 | 87 | -6 | 93 | -7 | 95 | 15 | 94 | -5 | 91 | -9 | 95 | 3 | 96 | 0 | 85 | -3 | 89 | 21 | 95 | -5 | 92 | 11 | 97 | 2 |
| CAN89 | 86 | -11 | 94 | -6 | 84 | -12 | 95 | -1 | 92 | -9 | 95 | -5 | 83 | -17 | 87 | -13 | 90 | -6 | 19 | -5 | 95 | -3 | 74 | -11 | 87 | -13 | 82 | -4 | 90 | -4 |
| CAN9 | 74 | 45 | 93 | -7 | 70 | -20 | 89 | 9 | 95 | 0 | 95 | -5 | 91 | 5 | 90 | 0 | 92 | -5 | 26 | 6 | 85 | 5 | 72 | -15 | 88 | 32 | 92 | -8 | 85 | -10 |
| CAN90 | 88 | -2 | 94 | 7 | 82 | 25 | 90 | 9 | 92 | -8 | 95 | 0 | 78 | -22 | 64 | 6 | 82 | -18 | 64 | -27 | 86 | 13 | 84 | -16 | 85 | -6 | 74 | -25 | 89 | -1 |
| CAN91 | 90 | -5 | 93 | -7 | 81 | 22 | 87 | -6 | 79 | -16 | 95 | -3 | 69 | -1 | 72 | 52 | 92 | -3 | 46 | 31 | 89 | -10 | 63 | 51 | 86 | -14 | 87 | -5 | 75 | 60 |
| CAN92 | 88 | -5 | 93 | -7 | 64 | 2 | 87 | -6 | 94 | -6 | 95 | -5 | 31 | -12 | 58 | -19 | 85 | 1 | 28 | 9 | 87 | 1 | 35 | 10 | 82 | -18 | 54 | -26 | 25 | 0 |
| CAN93 | 90 | 4 | 94 | -2 | 76 | 2 | 86 | 5 | 90 | -10 | 95 | 1 | 79 | -21 | 92 | -8 | 92 | -1 | 77 | -18 | 85 | 0 | 72 | -9 | 58 | -42 | 88 | 1 | 80 | -20 |
| CAN94 | 90 | 7 | 94 | 35 | 77 | 9 | 88 | 2 | 86 | -14 | 82 | 55 | 85 | -9 | 88 | -12 | 92 | -8 | 58 | -32 | 86 | 0 | 69 | -4 | 19 | -19 | 90 | -6 | 88 | -5 |
| CAN95 | 69 | -27 | 94 | -6 | 84 | -16 | 90 | -10 | 94 | -7 | 95 | -5 | 80 | -20 | 83 | -17 | 48 | -48 | 24 | 8 | 87 | 22 | 68 | 23 | 86 | 3 | 56 | -9 | 72 | -5 |
| CAN96 | 89 | -3 | 93 | -3 | 85 | 4 | 80 | -10 | 57 | 29 | 95 | 1 | 42 | 11 | 82 | -15 | 96 | -1 | 38 | 9 | 57 | 18 | 81 | 1 | 90 | -10 | 89 | 2 | 45 | 18 |
| CAN97 | 92 | -1 | 94 | 9 | 77 | 0 | 87 | -7 | 42 | 13 | 95 | 17 | 70 | 11 | 87 | -12 | 94 | 3 | 34 | 8 | 81 | -11 | 63 | -8 | 73 | -7 | 87 | 11 | 58 | 33 |
| CAN98 | 92 | 10 | 94 | 79 | 83 | 9 | 92 | 65 | 93 | -3 | 95 | 88 | 88 | 30 | 91 | 12 | 76 | 31 | 95 | 1 | 91 | 1 | 83 | 10 | 90 | 9 | 92 | 33 | 93 | -5 |
| HB1 | 96 | -4 | 94 | -6 | 82 | 1 | 96 | -4 | 94 | 14 | 95 | -5 | 87 | 2 | 83 | -17 | 95 | -2 | 86 | 8 | 84 | -16 | 73 | 17 | 83 | -4 | 88 | 2 | 80 | 42 |
| HB10 | 93 | -4 | 93 | -7 | 83 | 13 | 89 | 6 | 23 | 0 | 95 | -4 | 32 | 9 | 76 | -4 | 94 | 1 | 24 | 18 | 83 | 7 | 61 | 15 | 78 | 5 | 88 | 9 | 63 | -25 |
| HB11 | 91 | 4 | 93 | -6 | 72 | 37 | 90 | 5 | 64 | 24 | 80 | -15 | 72 | 32 | 89 | 36 | 96 | -3 | 47 | 27 | 91 | 0 | 65 | 48 | 53 | 47 | 88 | -12 | 56 | 27 |
| HB12 | 92 | -8 | 94 | -4 | 77 | 3 | 88 | -10 | 78 | 47 | 95 | -3 | 62 | -7 | 89 | 0 | 95 | -5 | 22 | 15 | 72 | -28 | 68 | 45 | 69 | -10 | 87 | -8 | 95 | -5 |

| | | | | | | | | | | | | | | | | | | | | | | | | | | | | | | |
|---|---|---|---|---|---|---|---|---|---|---|---|---|---|---|---|---|---|---|---|---|---|---|---|---|---|---|---|---|---|---|
| **HB13** | 92 | 3 | 93 | -7 | 78 | -8 | 93 | 7 | 93 | -3 | 80 | -15 | 80 | -4 | 93 | -7 | 95 | 7 | 24 | -2 | 83 | -3 | 65 | -16 | 20 | 1 | 88 | 4 | 58 | -1 |
| **HB14** | 92 | -6 | 94 | -6 | 78 | -15 | 87 | -10 | 94 | 8 | 82 | 62 | 80 | -3 | 93 | -7 | 95 | 5 | 22 | -10 | 84 | 2 | 66 | -12 | 21 | 7 | 88 | 8 | 65 | 21 |
| **HB15** | 93 | -2 | 94 | -6 | 81 | -9 | 90 | -5 | 92 | -3 | 95 | -1 | 95 | 8 | 78 | -9 | 68 | -32 | 85 | -2 | 96 | -1 | 70 | -14 | 78 | -4 | 89 | -1 | 78 | 17 |
| **HB16** | 92 | -2 | 93 | -6 | 83 | 2 | 96 | 8 | 94 | -1 | 95 | -5 | 83 | 12 | 87 | -1 | 95 | -1 | 69 | -18 | 85 | 1 | 34 | 16 | 86 | 13 | 88 | -3 | 69 | -22 |
| **HB17** | 92 | -4 | 93 | -7 | 76 | -8 | 89 | -10 | 93 | -4 | 95 | -5 | 82 | 6 | 87 | -4 | 95 | 6 | 75 | 1 | 86 | -2 | 31 | 6 | 85 | 8 | 88 | -2 | 72 | -13 |
| **HB18** | 89 | -4 | 94 | -6 | 89 | 10 | 83 | -11 | 91 | -3 | 95 | -2 | 75 | 0 | 91 | -1 | 96 | -4 | 82 | -6 | 87 | 0 | 78 | 1 | 70 | -6 | 89 | 8 | 88 | -5 |
| **HB19** | 91 | -5 | 94 | 60 | 39 | 27 | 89 | -2 | 91 | -7 | 82 | 61 | 85 | 4 | 91 | -9 | 96 | 1 | 69 | -23 | 91 | 4 | 78 | -7 | 56 | -24 | 90 | 1 | 72 | -27 |
| **HB2** | 89 | 4 | 94 | 2 | 82 | 4 | 81 | 16 | 94 | 0 | 95 | -4 | 73 | 4 | 89 | -4 | 96 | 3 | 65 | 35 | 76 | 19 | 71 | -7 | 71 | 7 | 88 | -12 | 89 | -11 |
| **HB21** | 90 | 8 | 94 | -6 | 88 | -8 | 91 | 7 | 93 | 8 | 95 | -5 | 71 | -11 | 91 | -9 | 96 | -4 | 83 | -2 | 87 | 4 | 76 | -7 | 71 | 4 | 89 | 9 | 89 | -5 |
| **HB22** | 91 | -3 | 93 | -7 | 86 | 11 | 88 | -4 | 37 | 7 | 95 | -3 | 47 | 14 | 83 | 16 | 96 | -4 | 27 | 2 | 38 | 7 | 65 | -2 | 84 | 18 | 88 | -1 | 62 | 35 |
| **HB23** | 92 | 4 | 75 | 56 | 38 | -45 | 96 | 15 | 51 | 10 | 95 | -5 | 59 | 40 | 92 | 26 | 95 | 18 | 24 | 4 | 48 | -30 | 56 | 30 | 75 | 54 | 88 | 1 | 40 | 22 |
| **HB24** | 90 | -8 | 93 | -7 | 84 | -16 | 85 | -11 | 94 | 5 | 95 | -4 | 92 | 22 | 84 | -2 | 95 | -1 | 84 | 5 | 85 | -15 | 76 | -24 | 82 | 18 | 83 | -16 | 61 | 19 |
| **HB25** | 85 | -8 | 93 | -4 | 66 | -30 | 83 | -8 | 93 | -4 | 95 | -3 | 89 | -4 | 91 | -9 | 91 | -7 | 85 | -8 | 91 | -4 | 79 | -16 | 85 | -8 | 86 | -12 | 95 | -5 |
| **HB26** | 92 | -8 | 94 | -6 | 81 | -15 | 89 | -7 | 93 | -7 | 79 | -9 | 86 | 3 | 93 | -7 | 95 | -4 | 66 | -16 | 82 | 2 | 72 | -5 | 57 | 35 | 88 | 6 | 82 | -11 |
| **HB27** | 66 | -21 | 93 | 2 | 73 | -11 | 89 | 7 | 94 | 6 | 95 | -2 | 92 | 17 | 90 | -1 | 91 | -9 | 92 | 6 | 86 | 13 | 78 | -1 | 82 | 4 | 61 | -38 | 93 | -7 |
| **HB28** | 90 | 6 | 94 | -6 | 81 | -1 | 82 | 8 | 94 | 2 | 95 | -5 | 69 | -14 | 89 | -3 | 96 | -1 | 48 | -20 | 85 | -3 | 73 | 2 | 73 | 6 | 89 | 2 | 92 | -8 |
| **HB29** | 94 | 5 | 93 | -7 | 76 | -8 | 91 | 6 | 92 | 4 | 95 | -2 | 77 | 28 | 92 | -8 | 95 | -5 | 28 | -34 | 84 | -2 | 66 | -10 | 77 | 30 | 87 | -3 | 88 | -3 |
| **HB3** | 93 | -7 | 94 | -6 | 82 | -7 | 91 | -3 | 67 | 61 | 95 | -5 | 75 | 66 | 81 | -8 | 96 | 3 | 70 | 65 | 85 | 5 | 71 | -9 | 76 | 5 | 86 | 1 | 60 | 55 |
| **HB30** | 93 | -2 | 94 | 49 | 77 | -5 | 90 | -3 | 26 | 14 | 95 | -3 | 42 | 34 | 81 | 35 | 95 | 8 | 31 | 14 | 48 | 4 | 49 | -19 | 72 | -3 | 88 | 5 | 46 | 35 |
| **HB35** | 92 | -5 | 94 | 71 | 78 | 13 | 91 | 4 | 49 | -9 | 95 | -5 | 48 | -2 | 56 | 3 | 96 | -4 | 14 | 0 | 46 | 25 | 71 | 5 | 86 | 17 | 89 | 1 | 37 | 4 |
| **HB5** | 92 | 2 | 94 | -6 | 87 | 40 | 92 | 4 | 95 | 7 | 80 | -19 | 89 | -2 | 91 | -9 | 95 | -5 | 29 | 9 | 86 | -1 | 79 | 10 | 43 | 38 | 89 | 6 | 91 | -1 |
| **HB6** | 91 | -3 | 93 | 1 | 77 | 1 | 87 | -6 | 55 | -39 | 95 | -1 | 89 | 1 | 89 | -11 | 96 | 0 | 52 | 35 | 90 | -10 | 77 | -1 | 83 | 3 | 86 | -14 | 83 | -9 |
| **HB7** | 88 | -9 | 93 | -7 | 69 | -1 | 88 | -6 | 90 | -4 | 95 | -5 | 90 | -10 | 91 | -9 | 95 | -1 | 77 | -18 | 87 | -6 | 80 | 1 | 92 | -8 | 87 | 4 | 97 | 3 |
| **HB8** | 94 | -3 | 93 | -7 | 82 | -9 | 92 | 4 | 30 | -63 | 95 | -5 | 48 | -25 | 82 | -2 | 96 | -4 | 35 | -54 | 85 | 3 | 76 | 11 | 71 | -3 | 86 | -2 | 29 | -32 |
| **HB9** | 93 | -5 | 94 | -6 | 83 | 2 | 92 | -2 | 93 | 0 | 95 | -3 | 94 | -4 | 93 | 6 | 95 | 4 | 83 | 57 | 83 | -11 | 79 | 4 | 89 | -1 | 88 | 2 | 94 | 2 |
| **HU1** | 92 | 3 | 93 | -7 | 77 | -9 | 89 | 4 | 46 | -19 | 95 | -5 | 48 | -15 | 92 | 9 | 93 | -1 | 21 | -20 | 61 | 25 | 62 | -7 | 77 | -23 | 89 | 1 | 51 | -43 |
| **HU11** | 92 | 5 | 93 | 4 | 78 | 19 | 82 | -4 | 37 | 5 | 81 | 0 | 30 | -5 | 91 | 2 | 95 | 3 | 46 | 17 | 79 | -3 | 55 | 28 | 29 | 4 | 93 | 8 | 37 | 8 |
| **HU12** | 91 | -7 | 93 | -7 | 68 | -28 | 90 | -1 | 94 | 0 | 95 | -5 | 77 | -16 | 91 | -2 | 96 | 0 | 77 | -12 | 88 | 10 | 79 | 5 | 44 | 18 | 91 | 8 | 52 | 21 |
| **HU13** | 91 | 4 | 93 | 1 | 75 | -12 | 87 | -3 | 93 | 0 | 74 | 59 | 89 | -5 | 82 | 59 | 96 | -1 | 50 | 46 | 91 | 2 | 60 | 38 | 70 | -22 | 89 | -6 | 51 | 37 |
| **HU14** | 93 | 3 | 94 | 54 | 68 | 7 | 85 | -7 | 40 | 14 | 80 | -20 | 36 | 24 | 91 | -4 | 95 | 8 | 47 | 29 | 84 | -2 | 42 | 15 | 27 | 13 | 75 | -1 | 59 | 33 |
| **HU17** | 91 | 3 | 94 | -6 | 83 | 0 | 83 | -1 | 63 | 16 | 95 | -4 | 69 | -1 | 93 | -2 | 96 | 0 | 37 | 21 | 85 | -4 | 72 | -10 | 71 | -11 | 88 | -1 | 47 | 28 |
| **HU18** | 92 | -3 | 54 | -42 | 60 | 38 | 83 | -17 | 54 | 21 | 95 | 11 | 44 | -12 | 88 | -8 | 95 | 10 | 28 | 23 | 68 | 38 | 46 | -6 | 72 | -26 | 88 | 13 | 43 | 32 |
| **HU19** | 91 | 3 | 93 | -4 | 82 | 10 | 91 | 13 | 93 | 1 | 95 | 3 | 83 | -8 | 92 | 5 | 95 | -5 | 85 | -11 | 93 | 15 | 80 | 10 | 84 | 16 | 91 | 4 | 95 | -5 |
| **HU2** | 90 | -4 | 94 | 5 | 83 | 17 | 90 | -6 | 90 | -10 | 95 | 7 | 90 | 2 | 93 | 7 | 96 | -4 | 70 | -23 | 84 | -9 | 80 | 9 | 78 | -22 | 89 | -3 | 65 | -25 |
| **HU21** | 92 | 12 | 93 | -7 | 83 | -1 | 88 | 13 | 77 | -18 | 80 | -20 | 81 | -8 | 93 | -7 | 95 | -1 | 43 | 28 | 84 | -2 | 62 | -6 | 19 | -4 | 88 | 9 | 45 | -22 |
| **HU22** | 89 | 3 | 94 | 12 | 77 | 3 | 84 | -5 | 89 | 9 | 81 | 29 | 78 | -6 | 91 | 1 | 93 | -6 | 49 | -20 | 63 | -23 | 58 | -17 | 24 | 12 | 91 | 4 | 82 | -12 |
| **HU23** | 92 | -6 | 94 | -6 | 80 | 13 | 85 | -13 | 93 | -3 | 81 | 66 | 88 | 10 | 93 | -1 | 95 | 3 | 31 | 24 | 84 | 5 | 74 | 37 | 19 | -4 | 88 | -2 | 73 | -17 |
| **HU25** | 89 | -4 | 93 | -7 | 71 | -8 | 88 | 11 | 58 | -42 | 95 | -5 | 71 | -10 | 87 | -9 | 92 | -8 | 42 | -49 | 81 | 13 | 72 | 3 | 56 | 22 | 91 | -1 | 75 | 41 |
| **HU26** | 88 | -8 | 93 | -7 | 85 | 0 | 89 | -5 | 56 | 11 | 95 | -2 | 66 | -22 | 79 | -16 | 96 | 0 | 23 | 2 | 75 | -15 | 83 | 8 | 94 | 27 | 88 | -5 | 73 | 30 |
| **HU27** | 89 | 0 | 94 | 3 | 68 | 14 | 89 | 9 | 29 | -2 | 95 | 3 | 54 | -4 | 58 | -34 | 95 | 0 | 26 | 2 | 65 | 34 | 47 | -9 | 75 | -6 | 88 | 4 | 47 | 7 |
| **HU28** | 89 | -2 | 93 | -7 | 61 | -10 | 88 | -6 | 93 | -4 | 95 | 10 | 91 | 8 | 92 | 5 | 96 | 8 | 92 | 6 | 95 | -5 | 83 | 21 | 88 | 5 | 89 | 2 | 91 | -1 |
| **HU3** | 92 | 2 | 60 | 31 | 71 | 41 | 91 | -1 | 93 | -7 | 95 | -5 | 86 | -14 | 92 | 48 | 95 | 5 | 71 | -19 | 91 | -4 | 65 | -7 | 68 | 50 | 88 | 4 | 85 | -12 |
| **HU30** | 89 | -9 | 94 | 1 | 78 | 0 | 85 | -5 | 93 | -2 | 95 | -5 | 86 | 10 | 93 | 2 | 95 | -5 | 62 | 24 | 82 | 7 | 64 | -13 | 84 | 3 | 83 | -4 | 87 | -11 |
| **HU31** | 91 | 4 | 94 | -6 | 76 | 3 | 88 | 7 | 50 | -36 | 95 | -2 | 86 | 8 | 91 | -9 | 95 | 7 | 25 | 4 | 86 | 4 | 53 | -18 | 69 | -13 | 91 | 3 | 64 | -21 |
| **HU32** | 92 | -8 | 94 | -6 | 80 | 7 | 89 | 22 | 94 | 27 | 95 | 1 | 83 | 20 | 93 | -7 | 95 | -5 | 25 | -44 | 86 | 13 | 51 | -18 | 36 | 28 | 88 | 1 | 70 | -20 |
| **HU35** | 92 | -5 | 94 | 50 | 57 | 10 | 91 | -4 | 93 | -2 | 95 | -5 | 78 | -7 | 82 | 13 | 95 | -3 | 44 | 19 | 91 | 5 | 35 | 5 | 84 | -5 | 88 | 3 | 56 | 39 |
| **HU37** | 91 | 0 | 93 | -7 | 82 | -7 | 89 | -2 | 94 | 4 | 95 | -5 | 91 | 9 | 79 | -9 | 95 | 4 | 96 | -1 | 91 | 3 | 71 | -11 | 84 | 13 | 89 | 1 | 77 | 49 |
| **HU39** | 80 | 22 | 94 | -2 | 82 | 13 | 87 | -2 | 58 | 35 | 95 | 0 | 51 | 30 | 79 | 47 | 95 | 60 | 24 | 17 | 58 | -24 | 62 | -23 | 77 | 15 | 92 | 47 | 36 | 9 |
| **HU40** | 96 | -4 | 94 | 3 | 33 | 5 | 89 | -11 | 71 | -15 | 80 | -4 | 64 | -16 | 91 | -9 | 95 | 2 | 61 | -18 | 95 | -5 | 75 | -6 | 63 | -22 | 88 | 1 | 68 | -27 |
| **HU41** | 87 | 6 | 93 | -7 | 80 | -19 | 85 | 14 | 90 | -4 | 95 | -5 | 89 | -3 | 91 | -3 | 92 | 3 | 65 | -31 | 82 | 17 | 70 | -19 | 84 | 0 | 90 | 9 | 92 | -8 |
| **HU42** | 91 | -9 | 93 | -7 | 79 | 34 | 89 | -11 | 83 | 59 | 95 | -5 | 72 | 34 | 76 | 46 | 95 | -5 | 60 | 31 | 71 | -20 | 64 | 38 | 84 | 2 | 86 | 64 | 54 | 23 |
| **HU43** | 92 | 2 | 93 | -7 | 81 | -13 | 90 | 2 | 70 | 24 | 95 | -5 | 55 | -9 | 81 | -2 | 95 | 3 | 29 | -8 | 89 | -4 | 60 | 10 | 80 | -6 | 88 | -4 | 60 | -3 |
| **HU44** | 89 | -8 | 93 | -2 | 77 | 4 | 85 | -13 | 87 | -6 | 81 | 39 | 86 | -1 | 91 | -6 | 92 | -8 | 56 | -21 | 84 | -3 | 75 | 1 | 22 | 6 | 91 | 5 | 89 | -7 |

| | | | | | | | | | | | | | | | | | | | | | | | | | | | | | | |
|---|---|---|---|---|---|---|---|---|---|---|---|---|---|---|---|---|---|---|---|---|---|---|---|---|---|---|---|---|---|---|
| **HU45** | 91 | 8 | 94 | -6 | 89 | 4 | 85 | 6 | 94 | -3 | 95 | -5 | 72 | -8 | 93 | -2 | 96 | 4 | 56 | -42 | 85 | 12 | 76 | 6 | 71 | 1 | 88 | 3 | 90 | 8 |
| **HU46** | 95 | -5 | 94 | 1 | 79 | 8 | 87 | -9 | 67 | -28 | 95 | 4 | 72 | -16 | 80 | -20 | 95 | -5 | 39 | -11 | 90 | -1 | 58 | -3 | 72 | -5 | 88 | -5 | 61 | -27 |
| **HU47** | 92 | -2 | 94 | 39 | 86 | 5 | 88 | -6 | 93 | 65 | 81 | 34 | 83 | 43 | 92 | 56 | 95 | 5 | 41 | 23 | 89 | -2 | 66 | 54 | 29 | -14 | 88 | 8 | 70 | 61 |
| **HU5** | 92 | -5 | 94 | 3 | 80 | 9 | 90 | -6 | 80 | 42 | 80 | 1 | 84 | -2 | 92 | 50 | 95 | 7 | 37 | 10 | 85 | 0 | 55 | 25 | 33 | 2 | 88 | 12 | 77 | 60 |
| **HU50** | 96 | -4 | 94 | -6 | 83 | 7 | 95 | -5 | 94 | 0 | 95 | -4 | 86 | -2 | 84 | -13 | 95 | -5 | 78 | -14 | 84 | -10 | 67 | 0 | 85 | 0 | 88 | -2 | 68 | 4 |
| **HU52** | 91 | 36 | 94 | 5 | 85 | -9 | 80 | -11 | 94 | -2 | 95 | -3 | 89 | 0 | 78 | -22 | 96 | -3 | 73 | 16 | 87 | -6 | 83 | -13 | 64 | -13 | 89 | 11 | 80 | -18 |
| **HU53** | 75 | -23 | 94 | -6 | 79 | -19 | 93 | -1 | 23 | 6 | 80 | -18 | 36 | 6 | 74 | -22 | 75 | -15 | 22 | 10 | 90 | -1 | 39 | 2 | 22 | 6 | 68 | -20 | 40 | 4 |
| **HU54** | 92 | -1 | 94 | 11 | 77 | -4 | 87 | 0 | 51 | 1 | 95 | -5 | 51 | -4 | 90 | -10 | 92 | -6 | 29 | 11 | 59 | -24 | 56 | -20 | 77 | 10 | 89 | 1 | 88 | 2 |
| **HU55** | 90 | 4 | 93 | -7 | 85 | 2 | 87 | 2 | 74 | -3 | 95 | -5 | 87 | 0 | 81 | -19 | 95 | 6 | 49 | -15 | 74 | 3 | 73 | 64 | 77 | 11 | 88 | 1 | 41 | -15 |
| **HU57** | 92 | 3 | 93 | -7 | 83 | -3 | 88 | 1 | 94 | 1 | 95 | -5 | 82 | -9 | 85 | -7 | 96 | 3 | 72 | -25 | 84 | -7 | 70 | 18 | 88 | -6 | 84 | 2 | 94 | -6 |
| **HU58** | 92 | -7 | 94 | 6 | 84 | 14 | 88 | -12 | 68 | -22 | 60 | 38 | 76 | -12 | 75 | -19 | 95 | -5 | 47 | 17 | 81 | -15 | 57 | -12 | 41 | 16 | 88 | 2 | 67 | -28 |
| **HU59** | 74 | -26 | 93 | -7 | 81 | -6 | 94 | -6 | 23 | 2 | 80 | -20 | 40 | -11 | 90 | -3 | 95 | 3 | 22 | 1 | 90 | -1 | 36 | -7 | 20 | -1 | 88 | 8 | 43 | -6 |
| **HU6** | 91 | 2 | 93 | -7 | 77 | 4 | 92 | 3 | 94 | 3 | 95 | -1 | 92 | 7 | 86 | 0 | 95 | 5 | 76 | -20 | 84 | 2 | 81 | 10 | 76 | -16 | 89 | 4 | 61 | 19 |
| **HU60** | 91 | -1 | 94 | 9 | 78 | 16 | 91 | 1 | 91 | 0 | 95 | 6 | 72 | -1 | 90 | -2 | 95 | 4 | 20 | 8 | 83 | -3 | 64 | 5 | 62 | -10 | 90 | -3 | 90 | -4 |
| **HU61** | 90 | -6 | 94 | -6 | 82 | 2 | 87 | -13 | 54 | 30 | 95 | -2 | 52 | 31 | 75 | 30 | 95 | 5 | 23 | 6 | 71 | 26 | 53 | -16 | 79 | 1 | 88 | -6 | 47 | 30 |
| **HU62** | 90 | -8 | 93 | -7 | 82 | 11 | 89 | -4 | 66 | -27 | 95 | -4 | 54 | -37 | 93 | -7 | 96 | -4 | 56 | -41 | 85 | 1 | 71 | -5 | 79 | 5 | 88 | 1 | 68 | -25 |
| **HU63** | 88 | -12 | 93 | -3 | 79 | -10 | 89 | -2 | 39 | -1 | 95 | -5 | 45 | 16 | 93 | 53 | 96 | -4 | 28 | -1 | 73 | -13 | 80 | -3 | 72 | -11 | 87 | -7 | 66 | -25 |
| **HU64** | 93 | -5 | 94 | -3 | 72 | -25 | 90 | -10 | 94 | -1 | 95 | 6 | 89 | 3 | 91 | -1 | 95 | -5 | 58 | -34 | 95 | -5 | 79 | -13 | 88 | 10 | 89 | 5 | 83 | -14 |
| **HU65** | 90 | 56 | 94 | 28 | 77 | 13 | 88 | -5 | 82 | -8 | 95 | -5 | 72 | 8 | 67 | 36 | 94 | 58 | 84 | 64 | 89 | 1 | 62 | 37 | 61 | 24 | 89 | 56 | 65 | 41 |
| **HU66** | 92 | 0 | 94 | -3 | 78 | 9 | 91 | -2 | 93 | -7 | 95 | -5 | 77 | -11 | 92 | 5 | 93 | -1 | 65 | -28 | 83 | -3 | 35 | -6 | 82 | -18 | 88 | 6 | 86 | -3 |
| **HU67** | 94 | 1 | 94 | -5 | 80 | -20 | 86 | -6 | 21 | -9 | 95 | -3 | 25 | -12 | 71 | -16 | 94 | -4 | 18 | -7 | 76 | -14 | 44 | -56 | 73 | 4 | 88 | -5 | 59 | -41 |
| **HU68** | 65 | -21 | 94 | -6 | 65 | -10 | 86 | 3 | 94 | -1 | 95 | 1 | 74 | -11 | 90 | -2 | 95 | -5 | 86 | -3 | 76 | 52 | 74 | 2 | 90 | 24 | 68 | -19 | 77 | 38 |
| **HU69** | 90 | 1 | 93 | -7 | 86 | 9 | 87 | -3 | 85 | 55 | 95 | 2 | 77 | 48 | 87 | 50 | 96 | 6 | 24 | -4 | 85 | -1 | 78 | 36 | 67 | 4 | 91 | 0 | 67 | 36 |
| **HU7** | 93 | -1 | 94 | -6 | 57 | 35 | 90 | -3 | 94 | 5 | 95 | -5 | 76 | -8 | 91 | 26 | 95 | -5 | 66 | -30 | 92 | 6 | 77 | 2 | 84 | 34 | 85 | -1 | 86 | -11 |
| **HU70** | 92 | 1 | 94 | -5 | 78 | 13 | 91 | -5 | 21 | 0 | 95 | 5 | 55 | 17 | 73 | -27 | 95 | -5 | 21 | 9 | 87 | 4 | 52 | 19 | 46 | 29 | 88 | -2 | 42 | 9 |
| **HU71** | 89 | -4 | 94 | -6 | 82 | -3 | 91 | 0 | 67 | -24 | 95 | 1 | 85 | 1 | 89 | -11 | 92 | 3 | 28 | -57 | 88 | 4 | 66 | -6 | 84 | 0 | 89 | 3 | 62 | -37 |
| **HU72** | 91 | -2 | 93 | 2 | 79 | 5 | 90 | -3 | 95 | 23 | 95 | -3 | 84 | 29 | 90 | 16 | 95 | -3 | 95 | 16 | 91 | -4 | 80 | 2 | 80 | 27 | 90 | -4 | 93 | -5 |
| **HU73** | 92 | -1 | 93 | -7 | 84 | 6 | 87 | -2 | 69 | -22 | 80 | 11 | 80 | 20 | 93 | 7 | 95 | 4 | 25 | 1 | 78 | 5 | 73 | 13 | 22 | -3 | 88 | 0 | 70 | 50 |
| **HU74** | 84 | -8 | 93 | -7 | 66 | 1 | 91 | 4 | 94 | 5 | 95 | -2 | 89 | 1 | 91 | 0 | 95 | -5 | 76 | -18 | 92 | 4 | 64 | 23 | 86 | 4 | 89 | 3 | 84 | -16 |
| **HU75** | 91 | 0 | 94 | -6 | 87 | -12 | 88 | -2 | 47 | 17 | 95 | -5 | 61 | 32 | 83 | 16 | 96 | 0 | 40 | 12 | 72 | 37 | 74 | 26 | 81 | 1 | 88 | 2 | 67 | 36 |
| **HU76** | 92 | -8 | 94 | -2 | 77 | 7 | 93 | 3 | 47 | -40 | 95 | 2 | 63 | -11 | 92 | 9 | 95 | -5 | 26 | -58 | 83 | 7 | 60 | -14 | 74 | 8 | 88 | 1 | 94 | -6 |
| **HU77** | 92 | 9 | 94 | -6 | 81 | -14 | 87 | 9 | 63 | 8 | 95 | -5 | 70 | -3 | 90 | -3 | 96 | -3 | 36 | 16 | 87 | -1 | 71 | -5 | 78 | 14 | 88 | 6 | 39 | 15 |
| **HU78** | 92 | 18 | 94 | -5 | 77 | -5 | 90 | 15 | 49 | 30 | 81 | -19 | 36 | 17 | 91 | -9 | 95 | -1 | 50 | 45 | 84 | -2 | 57 | 27 | 33 | 17 | 89 | 6 | 44 | 26 |
| **HU79** | 92 | 9 | 76 | 58 | 61 | 47 | 92 | 16 | 94 | 22 | 95 | -5 | 90 | 6 | 91 | -9 | 96 | 0 | 62 | -7 | 86 | 11 | 80 | 1 | 94 | 9 | 85 | -3 | 77 | -19 |
| **HU80** | 89 | -5 | 94 | 1 | 77 | -3 | 84 | -12 | 87 | -7 | 82 | 54 | 87 | 6 | 91 | 0 | 92 | -8 | 52 | 21 | 86 | 7 | 76 | 20 | 21 | -1 | 91 | -3 | 85 | -8 |
| **HU81** | 89 | 0 | 93 | -7 | 56 | 37 | 88 | -5 | 94 | -4 | 95 | -5 | 91 | 7 | 91 | 2 | 96 | 4 | 80 | -18 | 86 | -2 | 80 | 1 | 90 | 7 | 88 | 1 | 91 | -4 |
| **HU82** | 72 | -18 | 94 | 5 | 77 | 5 | 89 | 2 | 87 | -4 | 95 | 5 | 69 | -13 | 81 | -8 | 92 | -8 | 72 | -24 | 86 | -7 | 53 | -22 | 76 | 3 | 88 | 1 | 71 | -29 |
| **HU83** | 69 | 58 | 94 | -6 | 85 | -5 | 90 | -1 | 94 | 2 | 95 | -5 | 85 | 3 | 93 | 4 | 95 | 4 | 93 | -1 | 86 | 7 | 77 | -1 | 93 | 13 | 68 | 60 | 90 | 1 |
| **HU84** | 94 | 2 | 93 | -7 | 85 | 16 | 94 | -5 | 24 | 6 | 81 | -16 | 38 | 1 | 77 | -7 | 93 | -7 | 20 | 14 | 91 | 7 | 43 | 21 | 21 | 1 | 88 | -7 | 42 | 10 |
| **HU85** | 92 | -2 | 94 | 70 | 77 | -4 | 88 | -4 | 50 | -6 | 95 | -5 | 44 | -15 | 55 | -6 | 96 | -4 | 13 | -1 | 43 | 22 | 67 | -3 | 75 | -17 | 89 | -2 | 40 | 16 |
| **HU9** | 88 | -10 | 93 | -7 | 82 | -7 | 87 | -13 | 74 | -12 | 95 | -5 | 63 | -21 | 66 | -34 | 92 | -7 | 54 | -35 | 86 | -8 | 68 | -4 | 83 | -1 | 91 | 3 | 61 | -34 |
| **HU93** | 91 | -4 | 94 | -6 | 74 | -16 | 90 | -6 | 94 | 0 | 95 | -2 | 75 | 1 | 91 | 3 | 97 | 8 | 93 | 4 | 91 | -6 | 80 | -13 | 92 | 17 | 82 | -18 | 91 | -4 |
| **HU94** | 89 | -4 | 93 | -7 | 85 | -8 | 88 | -10 | 74 | 41 | 95 | -3 | 76 | 18 | 93 | 7 | 96 | -4 | 55 | -16 | 84 | -11 | 70 | -14 | 78 | 2 | 88 | -5 | 70 | 33 |
| **HU95** | 91 | -6 | 93 | -7 | 76 | -10 | 90 | -7 | 92 | 5 | 95 | -5 | 69 | -9 | 88 | -10 | 95 | -4 | 30 | 11 | 81 | -1 | 63 | 2 | 73 | 1 | 91 | 7 | 88 | -12 |
| **HU97** | 92 | 0 | 94 | 1 | 84 | 13 | 90 | 5 | 93 | -3 | 95 | 2 | 90 | 10 | 93 | -7 | 96 | -3 | 72 | -22 | 88 | -5 | 74 | -3 | 92 | 9 | 40 | -46 | 67 | -33 |

| | F1 | | FAT | | GEO | | GWF | | HAM3 | | HAM53 | | LEG | | LUC04 | | N | | NEA2 | | OIL | | P | | RV2 | | TB54 | | TB69 | | UFA5s | |
|---|---|---|---|---|---|---|---|---|---|---|---|---|---|---|---|---|---|---|---|---|---|---|---|---|---|---|---|---|---|---|---|---|
| **Isolate** | Pred. | Diff. | Pred. | Diff. | Pred. | Diff. | Pred. | Diff. | Pred. | Diff. | Pred. | Diff | Pred. | Diff. | Pred. | Diff. | Pred. | Diff. | Pred. | Diff. | Pred. | Diff. | Pred. | Diff | Pred. | Diff. | Pred. | Diff. | Pred. | Diff. | Pred. | Diff. |
| **CAN1** | 92 | 8 | 91 | -6 | 82 | -18 | 94 | 1 | 93 | -1 | 84 | 3 | 83 | -12 | 91 | -5 | 91 | -9 | 85 | -12 | 85 | -15 | 82 | -18 | 90 | 2 | 78 | 9 | 84 | -10 | 85 | 1 |
| **CAN10** | 87 | -7 | 91 | -9 | 84 | -15 | 95 | 16 | 93 | -7 | 92 | 6 | 90 | 3 | 91 | -3 | 91 | -9 | 76 | -3 | 85 | -14 | 83 | 54 | 94 | 2 | 93 | -7 | 84 | -13 | 86 | -6 |
| **CAN100** | 92 | -7 | 91 | 3 | 56 | 6 | 93 | 3 | 87 | 2 | 61 | -38 | 79 | 13 | 91 | -5 | 91 | 5 | 87 | -12 | 85 | -7 | 85 | 18 | 66 | -22 | 91 | 9 | 71 | -16 | 68 | 9 |

| | | | | | | | | | | | | | | | | | | | | | | | | | | | | | | | | |
|---|---|---|---|---|---|---|---|---|---|---|---|---|---|---|---|---|---|---|---|---|---|---|---|---|---|---|---|---|---|---|---|---|
| **CAN101** | 92 | -5 | 28 | 9 | 69 | -17 | 92 | 15 | 93 | -3 | 80 | 0 | 68 | -16 | 91 | 0 | 37 | 19 | 83 | 5 | 86 | 7 | 24 | 14 | 93 | 8 | 71 | -19 | 88 | 8 | 85 | 3 |
| **CAN102** | 92 | 30 | 91 | 1 | 92 | -8 | 79 | -7 | 93 | -7 | 88 | -12 | 85 | -11 | 91 | -7 | 91 | -8 | 93 | 11 | 85 | -15 | 83 | -3 | 91 | -7 | 83 | -8 | 66 | 21 | 85 | -12 |
| **CAN103** | 85 | 14 | 91 | 4 | 88 | -1 | 94 | -6 | 93 | -7 | 83 | 19 | 91 | 8 | 91 | -1 | 91 | 1 | 34 | 4 | 85 | -12 | 85 | -1 | 63 | 30 | 86 | -14 | 81 | -19 | 80 | 14 |
| **CAN104** | 92 | -7 | 91 | -3 | 89 | -9 | 87 | 12 | 93 | -7 | 95 | -3 | 90 | 2 | 91 | -8 | 91 | -5 | 41 | 26 | 85 | -15 | 85 | -9 | 94 | -3 | 96 | -4 | 86 | -10 | 85 | -14 |
| **CAN105** | 92 | 4 | 78 | -7 | 85 | 9 | 85 | 5 | 93 | -4 | 39 | 13 | 68 | 4 | 91 | -3 | 91 | 4 | 39 | 17 | 85 | -4 | 59 | 14 | 56 | 39 | 78 | -9 | 56 | -9 | 80 | -5 |
| **CAN106** | 79 | -9 | 91 | -9 | 88 | 21 | 93 | 3 | 88 | -1 | 82 | 4 | 81 | 25 | 91 | -4 | 89 | 3 | 43 | 24 | 86 | -2 | 76 | 37 | 46 | 19 | 88 | 3 | 78 | 1 | 43 | 11 |
| **CAN107** | 92 | 5 | 91 | 2 | 77 | -2 | 95 | -5 | 93 | -7 | 87 | 11 | 93 | 13 | 91 | -6 | 91 | 3 | 80 | -15 | 86 | -6 | 86 | 0 | 65 | -28 | 99 | -1 | 88 | -12 | 89 | 11 |
| **CAN108** | 92 | -8 | 91 | -4 | 86 | 15 | 95 | 0 | 93 | 14 | 83 | 7 | 87 | -8 | 91 | -7 | 91 | -6 | 79 | -15 | 86 | -5 | 82 | -17 | 94 | 6 | 71 | -6 | 85 | 17 | 85 | 9 |
| **CAN109** | 84 | -6 | 91 | -9 | 87 | 14 | 94 | -6 | 93 | -4 | 88 | 7 | 93 | -7 | 91 | 3 | 82 | -16 | 81 | -2 | 86 | 14 | 77 | -23 | 89 | 4 | 82 | -18 | 80 | -20 | 76 | 5 |
| **CAN11** | 92 | 1 | 91 | -9 | 80 | -20 | 94 | -6 | 93 | 10 | 81 | -1 | 93 | -7 | 91 | -4 | 91 | -7 | 88 | -8 | 86 | -10 | 80 | -21 | 93 | 0 | 94 | 2 | 84 | 1 | 85 | -3 |
| **CAN110** | 72 | -28 | 91 | 1 | 91 | -9 | 94 | 8 | 93 | -7 | 84 | -16 | 95 | 9 | 91 | -5 | 91 | -9 | 72 | 29 | 85 | 0 | 85 | -1 | 92 | 9 | 99 | -1 | 88 | 7 | 85 | -9 |
| **CAN111** | 85 | 3 | 91 | -9 | 71 | -15 | 94 | -6 | 93 | -2 | 82 | 2 | 89 | -7 | 91 | -6 | 83 | -6 | 52 | 9 | 85 | -12 | 80 | -21 | 92 | 4 | 94 | -4 | 67 | -32 | 85 | -1 |
| **CAN112** | 92 | -7 | 91 | -9 | 88 | -12 | 80 | -19 | 93 | -6 | 85 | -12 | 94 | -7 | 91 | -3 | 91 | -7 | 86 | -8 | 85 | -9 | 83 | -12 | 73 | -21 | 91 | -4 | 67 | 19 | 80 | -19 |
| **CAN113** | 92 | 4 | 91 | -9 | 84 | -2 | 92 | -8 | 93 | -7 | 67 | 3 | 33 | 1 | 91 | 0 | 38 | 18 | 26 | 7 | 86 | -5 | 26 | -8 | 85 | 0 | 84 | -16 | 74 | -26 | 86 | 11 |
| **CAN114** | 92 | 14 | 91 | -5 | 73 | 37 | 93 | 3 | 93 | 1 | 70 | 7 | 59 | 4 | 91 | 0 | 37 | -7 | 29 | -4 | 86 | 5 | 27 | -2 | 78 | 50 | 82 | -4 | 76 | 44 | 86 | 20 |
| **CAN115** | 70 | -22 | 91 | -3 | 69 | -22 | 62 | 17 | 93 | -7 | 84 | -1 | 65 | -18 | 91 | 0 | 78 | -17 | 24 | -66 | 52 | 2 | 75 | -5 | 67 | -25 | 88 | -12 | 85 | -8 | 55 | -34 |
| **CAN116** | 77 | -9 | 91 | -9 | 94 | 4 | 94 | -2 | 93 | -7 | 81 | -10 | 91 | -9 | 91 | -9 | 91 | -9 | 67 | 8 | 85 | -15 | 86 | -1 | 94 | -7 | 97 | -3 | 84 | -13 | 85 | 0 |
| **CAN117** | 53 | 45 | 91 | 61 | 85 | 54 | 94 | 25 | 93 | 0 | 84 | 9 | 90 | 10 | 91 | 74 | 92 | 35 | 84 | 75 | 86 | 72 | 84 | 43 | 84 | 74 | 86 | 62 | 89 | 12 | 85 | 76 |
| **CAN118** | 92 | -8 | 91 | 0 | 82 | -18 | 80 | -20 | 93 | -7 | 90 | -10 | 86 | 8 | 91 | 0 | 91 | -9 | 83 | -11 | 85 | -15 | 84 | 3 | 89 | 2 | 84 | -16 | 80 | -17 | 85 | -10 |
| **CAN119** | 62 | 17 | 91 | 27 | 81 | 1 | 84 | 27 | 93 | 10 | 84 | 23 | 91 | 38 | 74 | 48 | 61 | -19 | 32 | 5 | 86 | -10 | 79 | 26 | 86 | 53 | 70 | 41 | 58 | 7 | 82 | 55 |
| **CAN12** | 92 | 1 | 91 | -9 | 87 | -13 | 92 | -8 | 93 | -6 | 82 | 2 | 77 | -3 | 91 | -8 | 91 | -9 | 89 | 2 | 86 | -14 | 83 | 2 | 92 | 0 | 82 | -18 | 88 | 5 | 85 | -1 |
| **CAN120** | 92 | 2 | 31 | 22 | 56 | -11 | 94 | 5 | 93 | -6 | 39 | 19 | 38 | 29 | 91 | -3 | 28 | -8 | 58 | -31 | 86 | -7 | 28 | 22 | 81 | -19 | 89 | 0 | 90 | 25 | 71 | -18 |
| **CAN121** | 92 | 2 | 91 | 7 | 54 | 27 | 93 | 0 | 89 | 53 | 64 | -25 | 75 | -8 | 91 | -4 | 91 | 8 | 54 | -28 | 86 | 3 | 62 | -18 | 82 | -9 | 77 | -14 | 62 | 11 | 82 | -11 |
| **CAN122** | 92 | -1 | 91 | -6 | 88 | 12 | 96 | 1 | 93 | -7 | 86 | 3 | 94 | -6 | 91 | -9 | 91 | 5 | 74 | 10 | 86 | 1 | 83 | -4 | 87 | -8 | 97 | -3 | 85 | -2 | 85 | -2 |
| **CAN123** | 92 | 1 | 91 | 7 | 85 | 72 | 94 | 10 | 93 | -2 | 80 | 1 | 83 | 11 | 91 | -4 | 91 | 9 | 94 | 11 | 86 | 1 | 84 | 14 | 92 | 0 | 96 | 12 | 86 | 18 | 86 | 4 |
| **CAN124** | 84 | -16 | 91 | -9 | 68 | 5 | 79 | 7 | 93 | 24 | 92 | 14 | 95 | -5 | 91 | -2 | 91 | 9 | 65 | 4 | 86 | 16 | 87 | 2 | 85 | -1 | 71 | 3 | 61 | -2 | 81 | 7 |
| **CAN125** | 85 | 20 | 91 | -2 | 84 | -11 | 85 | -6 | 93 | 9 | 81 | -10 | 92 | 4 | 91 | 2 | 91 | 2 | 57 | -34 | 86 | 0 | 80 | 14 | 72 | -7 | 88 | -12 | 88 | -2 | 64 | -16 |
| **CAN126** | 92 | -2 | 91 | -2 | 50 | -45 | 94 | 5 | 87 | -4 | 64 | -31 | 92 | -5 | 91 | 4 | 91 | -4 | 86 | -3 | 86 | 76 | 83 | -9 | 90 | 2 | 91 | 1 | 62 | 34 | 82 | -6 |
| **CAN127** | 56 | 40 | 91 | 0 | 41 | 30 | 93 | 6 | 93 | 9 | 22 | 15 | 70 | 27 | 91 | -3 | 90 | 8 | 47 | 16 | 86 | 5 | 56 | 16 | 58 | 9 | 85 | 28 | 77 | 12 | 78 | 4 |
| **CAN128** | 92 | 3 | 91 | -9 | 81 | -3 | 92 | -5 | 93 | -7 | 86 | -2 | 84 | -16 | 91 | -2 | 91 | 4 | 86 | -5 | 85 | -15 | 87 | -6 | 87 | -12 | 87 | -8 | 85 | 53 | 60 | -35 |
| **CAN129** | 92 | 21 | 91 | 34 | 46 | 8 | 94 | 13 | 93 | -3 | 87 | 5 | 66 | 24 | 91 | -5 | 91 | 12 | 86 | -8 | 85 | -10 | 86 | 44 | 93 | 4 | 85 | -9 | 85 | 12 | 85 | -6 |
| **CAN13** | 92 | 6 | 91 | -9 | 57 | -25 | 94 | -6 | 93 | -5 | 83 | 7 | 88 | -12 | 91 | 1 | 92 | 15 | 61 | 1 | 86 | 10 | 84 | -16 | 75 | 47 | 94 | -6 | 89 | -3 | 78 | 40 |
| **CAN131** | 92 | -6 | 91 | 17 | 72 | -23 | 96 | -4 | 93 | -7 | 86 | -13 | 84 | 9 | 91 | -2 | 62 | 24 | 86 | -4 | 85 | -7 | 37 | 23 | 91 | -4 | 90 | 10 | 79 | 1 | 83 | -12 |
| **CAN132** | 92 | -8 | 91 | 13 | 82 | -12 | 92 | 21 | 93 | 10 | 90 | -1 | 91 | 16 | 91 | -7 | 91 | -8 | 80 | -12 | 85 | -7 | 83 | 10 | 92 | -1 | 93 | 29 | 83 | 20 | 87 | -5 |
| **CAN133** | 92 | -8 | 91 | -9 | 86 | -3 | 92 | -8 | 87 | -2 | 65 | -7 | 76 | -3 | 91 | 8 | 88 | -12 | 81 | -6 | 86 | 1 | 72 | -19 | 79 | 10 | 81 | -19 | 77 | -7 | 77 | 7 |
| **CAN134** | 51 | 26 | 91 | -9 | 36 | 12 | 93 | -2 | 93 | -1 | 21 | 11 | 63 | 1 | 91 | 5 | 91 | 5 | 44 | 8 | 86 | 7 | 55 | 7 | 60 | 15 | 69 | -16 | 70 | -10 | 78 | 14 |
| **CAN135** | 92 | 5 | 91 | 6 | 92 | 5 | 21 | 2 | 93 | 6 | 90 | 7 | 78 | 1 | 91 | 0 | 91 | 1 | 76 | -17 | 86 | -8 | 83 | 4 | 92 | 0 | 87 | -7 | 82 | -6 | 85 | 3 |
| **CAN136** | 71 | -26 | 91 | -4 | 89 | 7 | 93 | 3 | 93 | 9 | 87 | -3 | 94 | -6 | 91 | -9 | 91 | 0 | 82 | -8 | 85 | -2 | 89 | 4 | 92 | -6 | 88 | 9 | 85 | 3 | 85 | -8 |
| **CAN137** | 92 | -8 | 91 | 17 | 93 | 8 | 88 | 3 | 93 | -3 | 83 | -17 | 82 | 25 | 91 | -9 | 91 | -3 | 52 | 0 | 86 | -5 | 87 | 31 | 58 | -8 | 77 | -5 | 71 | -7 | 80 | -17 |
| **CAN138** | 92 | -5 | 91 | 13 | 93 | 9 | 22 | 12 | 93 | 7 | 89 | -8 | 79 | 4 | 91 | -5 | 92 | -3 | 75 | -20 | 85 | -10 | 82 | 4 | 91 | -6 | 90 | 19 | 84 | 14 | 85 | -15 |
| **CAN139** | 84 | -11 | 91 | 17 | 79 | 16 | 80 | 18 | 93 | 2 | 89 | -11 | 72 | 17 | 91 | -5 | 90 | 9 | 46 | 13 | 86 | 3 | 87 | 29 | 63 | 43 | 86 | 11 | 63 | 54 | 86 | 65 |
| **CAN14** | 92 | -6 | 91 | -5 | 89 | -11 | 94 | -6 | 93 | -7 | 88 | 17 | 90 | 9 | 91 | -9 | 91 | -9 | 81 | -19 | 86 | 46 | 86 | 4 | 82 | -9 | 92 | -8 | 88 | -5 | 87 | 9 |
| **CAN140** | 84 | -7 | 91 | 20 | 78 | 28 | 93 | 1 | 93 | 3 | 81 | 2 | 78 | 17 | 91 | -2 | 92 | 15 | 25 | 15 | 86 | 18 | 75 | -10 | 33 | 15 | 93 | 2 | 90 | -2 | 33 | 10 |
| **CAN141** | 92 | -3 | 77 | -4 | 87 | -4 | 68 | 9 | 93 | 7 | 58 | -37 | 82 | 10 | 91 | -6 | 91 | -4 | 75 | -10 | 86 | -5 | 83 | 8 | 88 | -5 | 70 | -6 | 49 | -19 | 84 | -16 |
| **CAN142** | 92 | 6 | 91 | -2 | 85 | -15 | 83 | -12 | 93 | 21 | 83 | -15 | 75 | -25 | 91 | 3 | 91 | -9 | 56 | 12 | 86 | 7 | 77 | -21 | 69 | 23 | 81 | 10 | 73 | 4 | 84 | 2 |
| **CAN143** | 85 | -4 | 91 | -3 | 85 | -1 | 92 | -8 | 93 | -7 | 90 | 0 | 81 | -10 | 91 | -1 | 92 | 3 | 74 | -18 | 86 | -11 | 86 | 1 | 75 | -10 | 98 | -2 | 79 | -21 | 86 | -3 |
| **CAN144** | 58 | -28 | 91 | 9 | 85 | -7 | 84 | 0 | 93 | -4 | 83 | 1 | 96 | 4 | 91 | 3 | 91 | 3 | 83 | -14 | 86 | -5 | 84 | 0 | 89 | 5 | 83 | -7 | 85 | 7 | 86 | 5 |
| **CAN145** | 92 | 5 | 91 | -4 | 90 | 22 | 48 | -8 | 93 | -5 | 90 | 4 | 77 | -6 | 91 | 1 | 92 | 13 | 76 | -10 | 86 | 4 | 83 | 4 | 93 | 7 | 85 | -15 | 82 | -5 | 84 | -1 |
| **CAN146** | 84 | -4 | 91 | -9 | 82 | -4 | 94 | -1 | 93 | -7 | 82 | 1 | 89 | -8 | 91 | -1 | 91 | -2 | 52 | -38 | 85 | -10 | 85 | -1 | 59 | -24 | 94 | -7 | 74 | -10 | 71 | -17 |
| **CAN147** | 92 | -5 | 91 | 4 | 89 | -6 | 95 | 11 | 93 | 0 | 80 | -18 | 81 | 3 | 91 | -5 | 91 | -8 | 58 | -4 | 86 | 46 | 87 | 5 | 87 | -6 | 80 | -4 | 83 | 11 | 58 | 17 |
| **CAN148** | 92 | -6 | 91 | -3 | 94 | 9 | 80 | -2 | 93 | 11 | 24 | 6 | 36 | 15 | 91 | -1 | 91 | 15 | 34 | 16 | 86 | 3 | 39 | 16 | 47 | 24 | 79 | 1 | 55 | 22 | 62 | 29 |
| **CAN149** | 92 | -8 | 91 | 2 | 88 | 1 | 92 | -8 | 93 | -7 | 36 | 12 | 40 | 16 | 91 | -7 | 91 | 45 | 50 | 18 | 86 | -6 | 39 | 18 | 50 | 25 | 85 | -15 | 84 | 19 | 56 | -4 |
| **CAN15** | 52 | 32 | 91 | -9 | 81 | -19 | 94 | -6 | 93 | -7 | 89 | 7 | 93 | 2 | 91 | -7 | 91 | -9 | 54 | -46 | 85 | -15 | 83 | -15 | 93 | -1 | 96 | -4 | 88 | -4 | 70 | -10 |

| | | | | | | | | | | | | | | | | | | | | | | | | | | | | | | | | |
|---|---|---|---|---|---|---|---|---|---|---|---|---|---|---|---|---|---|---|---|---|---|---|---|---|---|---|---|---|---|---|---|---|
| **CAN150** | 92 | 5 | 91 | -3 | 94 | -2 | 74 | -19 | 93 | -4 | 89 | -3 | 84 | -15 | 91 | -5 | 91 | -6 | 82 | -6 | 86 | -5 | 83 | -11 | 92 | 0 | 81 | -14 | 67 | 33 | 87 | -11 |
| **CAN151** | 83 | 7 | 91 | 2 | 76 | -12 | 94 | 3 | 87 | -13 | 83 | 14 | 88 | 7 | 91 | 3 | 91 | 3 | 73 | -17 | 86 | 0 | 86 | 6 | 71 | -7 | 96 | 3 | 83 | -7 | 86 | 15 |
| **CAN152** | 92 | -8 | 91 | -9 | 84 | 2 | 81 | 14 | 93 | -7 | 73 | -27 | 84 | -16 | 91 | 11 | 91 | -3 | 35 | -8 | 86 | -5 | 79 | -8 | 86 | 5 | 87 | -13 | 59 | 26 | 85 | -3 |
| **CAN153** | 92 | 2 | 31 | -53 | 74 | 22 | 94 | -6 | 93 | -4 | 79 | -9 | 36 | -44 | 91 | 2 | 43 | -51 | 59 | -35 | 85 | -11 | 18 | -56 | 81 | -7 | 70 | 26 | 87 | 0 | 79 | -7 |
| **CAN154** | 92 | 5 | 91 | -9 | 84 | -5 | 95 | -5 | 93 | -3 | 80 | -7 | 94 | -6 | 91 | -1 | 91 | 0 | 78 | -17 | 85 | -8 | 85 | -9 | 87 | -4 | 98 | 7 | 85 | -9 | 85 | -5 |
| **CAN155** | 92 | -1 | 91 | 3 | 83 | 0 | 76 | -15 | 93 | -1 | 78 | -6 | 86 | 7 | 91 | -4 | 91 | -9 | 42 | 16 | 86 | -4 | 81 | 4 | 83 | -10 | 87 | -13 | 47 | -17 | 85 | -6 |
| **CAN156** | 92 | 6 | 85 | 77 | 79 | -9 | 89 | 7 | 93 | 13 | 90 | 10 | 81 | 59 | 91 | 5 | 67 | 51 | 90 | 5 | 86 | 3 | 74 | 69 | 94 | 17 | 76 | 36 | 88 | 17 | 83 | 9 |
| **CAN157** | 92 | 12 | 91 | 13 | 92 | 21 | 80 | 10 | 93 | 7 | 79 | 10 | 88 | 23 | 91 | 2 | 91 | -4 | 36 | -2 | 86 | -6 | 84 | 23 | 87 | 6 | 95 | 18 | 55 | 14 | 86 | 14 |
| **CAN158** | 92 | -8 | 91 | -4 | 90 | 28 | 85 | -14 | 93 | -3 | 90 | -2 | 85 | -12 | 91 | -4 | 92 | 24 | 90 | 29 | 86 | 22 | 83 | -8 | 92 | -1 | 85 | -12 | 74 | -20 | 87 | 0 |
| **CAN159** | 84 | 1 | 91 | 5 | 81 | -5 | 93 | 5 | 93 | -7 | 78 | 10 | 81 | 7 | 91 | 3 | 91 | 6 | 83 | -5 | 86 | -11 | 83 | 14 | 80 | 1 | 90 | -1 | 69 | -12 | 75 | 11 |
| **CAN16** | 84 | -10 | 91 | -4 | 78 | -22 | 93 | 7 | 93 | 6 | 81 | -10 | 82 | 3 | 91 | 8 | 91 | -9 | 38 | 10 | 86 | -2 | 86 | 1 | 54 | 15 | 93 | -1 | 91 | 9 | 49 | 5 |
| **CAN160** | 79 | 14 | 91 | -7 | 87 | 4 | 78 | -12 | 93 | 11 | 81 | -19 | 95 | -5 | 91 | 43 | 91 | -2 | 46 | -21 | 86 | -4 | 87 | 4 | 82 | -8 | 75 | 9 | 73 | 0 | 80 | -4 |
| **CAN161** | 92 | -6 | 91 | 5 | 91 | 2 | 83 | 0 | 93 | 8 | 83 | -12 | 87 | 8 | 91 | -3 | 91 | -2 | 84 | -6 | 85 | -5 | 83 | -1 | 92 | -2 | 86 | 3 | 78 | -4 | 86 | -2 |
| **CAN162** | 86 | 30 | 91 | 2 | 84 | -16 | 90 | 1 | 93 | -1 | 89 | -8 | 68 | 4 | 91 | -3 | 91 | -9 | 50 | 8 | 86 | -10 | 81 | -5 | 64 | 18 | 74 | 21 | 44 | 7 | 85 | -8 |
| **CAN163** | 92 | -2 | 91 | -4 | 82 | -9 | 87 | -13 | 93 | -7 | 90 | -2 | 87 | 0 | 91 | -2 | 91 | -3 | 81 | -10 | 86 | -5 | 83 | -2 | 92 | 0 | 85 | -13 | 72 | -21 | 87 | -3 |
| **CAN164** | 92 | -1 | 91 | -7 | 89 | 3 | 85 | -5 | 93 | 5 | 65 | -13 | 40 | -9 | 91 | 0 | 37 | 7 | 22 | -13 | 86 | -1 | 27 | -5 | 86 | -6 | 87 | 0 | 70 | -6 | 76 | -14 |
| **CAN165** | 92 | -5 | 91 | -9 | 86 | 19 | 81 | -8 | 93 | -3 | 85 | -3 | 86 | -14 | 91 | -1 | 92 | 25 | 58 | 29 | 86 | 17 | 86 | -4 | 66 | 0 | 91 | -6 | 83 | -17 | 85 | -8 |
| **CAN166** | 92 | -6 | 91 | 0 | 95 | -5 | 83 | -1 | 93 | 3 | 89 | -4 | 94 | -6 | 91 | -8 | 91 | -9 | 88 | 14 | 86 | 8 | 82 | 0 | 91 | -9 | 90 | 3 | 47 | -33 | 85 | -5 |
| **CAN167** | 92 | 2 | 38 | 37 | 72 | -5 | 94 | 6 | 93 | -7 | 65 | 50 | 41 | 33 | 91 | 0 | 39 | 29 | 74 | 64 | 86 | 74 | 23 | 8 | 79 | 64 | 90 | 11 | 89 | 7 | 76 | 62 |
| **CAN168** | 92 | -8 | 91 | -4 | 82 | -8 | 92 | 5 | 93 | -7 | 79 | -21 | 93 | -7 | 91 | 6 | 91 | -1 | 88 | -2 | 85 | -8 | 84 | -11 | 83 | 1 | 98 | -2 | 39 | 21 | 84 | -9 |
| **CAN169** | 92 | -1 | 91 | 17 | 93 | 3 | 83 | 38 | 93 | 4 | 89 | -1 | 79 | 15 | 91 | 0 | 91 | 2 | 82 | -5 | 85 | -2 | 83 | 8 | 92 | 3 | 91 | 16 | 54 | -14 | 87 | 4 |
| **CAN17** | 92 | -2 | 47 | 31 | 67 | 35 | 94 | -1 | 93 | 16 | 83 | -9 | 61 | 25 | 91 | -2 | 51 | 28 | 89 | -11 | 86 | -14 | 36 | 19 | 90 | 0 | 84 | 50 | 87 | 24 | 85 | 3 |
| **CAN170** | 92 | 0 | 91 | 0 | 88 | -3 | 83 | -15 | 93 | -7 | 82 | -11 | 76 | -16 | 91 | 1 | 91 | -1 | 50 | 17 | 86 | -7 | 87 | 1 | 73 | -17 | 86 | -4 | 69 | -16 | 77 | -23 |
| **CAN171** | 92 | -2 | 91 | -4 | 84 | -6 | 85 | 1 | 93 | 4 | 87 | 1 | 86 | 0 | 91 | -4 | 91 | -9 | 68 | -32 | 85 | -15 | 85 | -12 | 87 | -8 | 86 | 0 | 84 | 2 | 85 | -9 |
| **CAN172** | 92 | 8 | 91 | 2 | 89 | 11 | 90 | 15 | 93 | 11 | 28 | 4 | 44 | 19 | 91 | -3 | 72 | 45 | 31 | 16 | 86 | 6 | 28 | 9 | 78 | -4 | 82 | 9 | 74 | 22 | 77 | 7 |
| **CAN173** | 88 | 7 | 47 | -40 | 76 | -17 | 90 | -5 | 93 | -7 | 78 | 6 | 33 | 13 | 91 | 0 | 72 | -22 | 93 | -5 | 86 | -12 | 26 | -3 | 89 | 3 | 63 | -37 | 88 | -10 | 53 | -22 |
| **CAN174** | 92 | -8 | 91 | 3 | 87 | -13 | 92 | 7 | 93 | 6 | 66 | -5 | 36 | -2 | 91 | -2 | 57 | 34 | 23 | -5 | 86 | 9 | 29 | 6 | 89 | -2 | 85 | 8 | 71 | -2 | 83 | 0 |
| **CAN175** | 92 | -8 | 91 | 9 | 88 | 1 | 85 | 4 | 93 | -5 | 89 | 19 | 89 | 26 | 91 | -2 | 91 | 4 | 43 | -3 | 85 | -7 | 89 | 9 | 77 | 14 | 89 | 5 | 76 | 5 | 76 | 0 |
| **CAN176** | 92 | -8 | 91 | 0 | 81 | -19 | 89 | 14 | 93 | -7 | 83 | -17 | 93 | 15 | 91 | 5 | 91 | 8 | 56 | 15 | 86 | -13 | 88 | 15 | 65 | -2 | 92 | -8 | 84 | -12 | 85 | -12 |
| **CAN177** | 92 | 7 | 91 | -1 | 85 | -11 | 79 | -17 | 93 | -7 | 89 | 8 | 96 | 7 | 91 | -9 | 91 | -5 | 84 | -12 | 86 | -10 | 81 | -4 | 85 | -9 | 91 | -7 | 80 | -19 | 85 | -2 |
| **CAN178** | 92 | 7 | 91 | 0 | 90 | -2 | 93 | 10 | 88 | 3 | 84 | 0 | 55 | -44 | 91 | -2 | 91 | -2 | 83 | -9 | 85 | -8 | 84 | 6 | 89 | -10 | 95 | 19 | 77 | 6 | 82 | -12 |
| **CAN179** | 70 | -19 | 91 | 2 | 69 | -15 | 96 | -1 | 93 | -7 | 83 | 5 | 92 | 9 | 91 | -7 | 91 | 1 | 76 | -18 | 86 | -6 | 87 | 12 | 75 | -9 | 99 | -1 | 88 | -12 | 85 | 1 |
| **CAN18** | 92 | -5 | 91 | -1 | 66 | 21 | 93 | 3 | 88 | 23 | 51 | 22 | 71 | 5 | 91 | 2 | 91 | 8 | 69 | -31 | 86 | -15 | 27 | -11 | 48 | 18 | 80 | -7 | 81 | -7 | 85 | -6 |
| **CAN180** | 84 | -6 | 91 | 3 | 60 | -28 | 93 | -3 | 93 | -2 | 81 | 5 | 72 | -25 | 91 | -8 | 91 | -5 | 25 | 14 | 86 | 36 | 74 | -13 | 30 | 8 | 95 | 3 | 90 | -3 | 32 | 7 |
| **CAN181** | 92 | 2 | 91 | -3 | 83 | 4 | 94 | -6 | 93 | -7 | 82 | 3 | 85 | 3 | 91 | -9 | 91 | 0 | 90 | -1 | 85 | -2 | 81 | 3 | 80 | -17 | 83 | -17 | 89 | 4 | 37 | -48 |
| **CAN182** | 92 | 1 | 91 | 15 | 36 | -23 | 93 | 13 | 70 | 31 | 84 | -3 | 91 | 9 | 91 | -1 | 91 | -5 | 84 | 4 | 86 | 6 | 93 | 15 | 89 | 3 | 94 | -2 | 50 | 13 | 84 | -9 |
| **CAN183** | 92 | -8 | 91 | 1 | 42 | 2 | 92 | 1 | 66 | 12 | 79 | -21 | 90 | 6 | 91 | -9 | 91 | -3 | 83 | -2 | 86 | -14 | 85 | -15 | 87 | -13 | 95 | 4 | 49 | 9 | 83 | -17 |
| **CAN184** | 92 | -7 | 91 | -9 | 84 | -1 | 90 | -6 | 93 | -7 | 88 | 6 | 94 | -6 | 91 | -2 | 91 | 2 | 88 | -6 | 86 | -9 | 83 | -17 | 74 | -23 | 98 | -2 | 32 | 13 | 84 | -8 |
| **CAN185** | 92 | 16 | 91 | 2 | 79 | -5 | 74 | -14 | 93 | -7 | 81 | 8 | 80 | -5 | 91 | -9 | 91 | 0 | 45 | 1 | 86 | 27 | 84 | 8 | 87 | -6 | 93 | -2 | 89 | -11 | 86 | 3 |
| **CAN186** | 85 | -12 | 91 | -3 | 86 | -12 | 77 | -15 | 83 | -2 | 83 | -4 | 84 | -16 | 91 | -1 | 91 | -9 | 53 | -47 | 85 | -15 | 86 | -7 | 66 | -24 | 85 | 6 | 72 | 0 | 78 | -21 |
| **CAN187** | 92 | -6 | 91 | -3 | 63 | 27 | 58 | 25 | 93 | 0 | 30 | 26 | 37 | 10 | 91 | -4 | 91 | -9 | 27 | 5 | 86 | 70 | 71 | -9 | 45 | 1 | 76 | 26 | 26 | 14 | 54 | 10 |
| **CAN188** | 92 | 4 | 91 | -8 | 59 | 20 | 56 | 16 | 93 | -6 | 18 | -14 | 34 | 2 | 91 | 0 | 91 | -9 | 29 | 11 | 86 | 70 | 73 | 0 | 42 | -7 | 64 | -11 | 26 | 15 | 56 | 16 |
| **CAN189** | 92 | 8 | 91 | 0 | 65 | 0 | 93 | 3 | 88 | -6 | 69 | -1 | 85 | -2 | 91 | -3 | 91 | 2 | 84 | 0 | 86 | 6 | 79 | -10 | 90 | 2 | 89 | -5 | 89 | -5 | 83 | 14 |
| **CAN19** | 92 | 4 | 91 | -6 | 97 | -3 | 94 | -2 | 93 | 6 | 53 | 27 | 76 | 28 | 91 | -2 | 91 | 35 | 75 | 44 | 85 | -15 | 43 | 23 | 82 | 2 | 79 | -8 | 84 | 29 | 85 | -2 |
| **CAN190** | 92 | -8 | 91 | 4 | 87 | -13 | 94 | -5 | 93 | 4 | 69 | 38 | 59 | 19 | 91 | -5 | 38 | 16 | 26 | 6 | 86 | 13 | 27 | 6 | 85 | 15 | 85 | 8 | 72 | 3 | 85 | 1 |
| **CAN191** | 84 | -9 | 91 | 8 | 85 | -9 | 94 | -4 | 93 | -7 | 86 | -1 | 81 | 5 | 74 | -16 | 91 | -7 | 69 | -21 | 86 | 22 | 83 | -5 | 88 | 2 | 87 | 1 | 88 | -9 | 85 | -1 |
| **CAN192** | 92 | -2 | 91 | -5 | 62 | 39 | 92 | -6 | 87 | 15 | 66 | -22 | 73 | -22 | 91 | 1 | 91 | 10 | 87 | 30 | 86 | 13 | 61 | 33 | 85 | -2 | 82 | -8 | 84 | -16 | 82 | -4 |
| **CAN193** | 79 | -5 | 91 | 2 | 83 | -5 | 94 | -6 | 88 | -12 | 82 | 0 | 87 | 3 | 91 | -6 | 91 | 0 | 74 | -21 | 85 | -7 | 85 | -2 | 70 | -9 | 94 | -6 | 81 | -19 | 85 | 6 |
| **CAN194** | 85 | 9 | 91 | -9 | 61 | -7 | 93 | -3 | 88 | 28 | 81 | -6 | 95 | -4 | 78 | 51 | 91 | -9 | 48 | -14 | 86 | -14 | 86 | -14 | 68 | 27 | 72 | 28 | 87 | -6 | 58 | 26 |
| **CAN195** | 84 | -9 | 91 | 11 | 87 | 9 | 78 | 6 | 93 | -5 | 87 | 12 | 80 | 15 | 91 | -2 | 91 | -1 | 64 | 21 | 86 | -7 | 77 | 4 | 76 | 45 | 82 | 7 | 86 | 7 | 80 | -2 |
| **CAN196** | 84 | -1 | 91 | -9 | 84 | 11 | 88 | 5 | 93 | 8 | 83 | 10 | 94 | -6 | 91 | 4 | 91 | -8 | 73 | -17 | 86 | -1 | 77 | -8 | 69 | -15 | 91 | 8 | 84 | 9 | 78 | 5 |
| **CAN197** | 81 | 53 | 91 | 11 | 84 | 5 | 94 | -6 | 87 | -13 | 84 | 6 | 81 | -18 | 76 | 33 | 91 | -9 | 86 | 38 | 86 | -7 | 76 | -13 | 91 | -5 | 87 | 7 | 67 | 40 | 85 | 9 |
| **CAN199** | 92 | 15 | 91 | -9 | 90 | -10 | 89 | -1 | 93 | 11 | 81 | 14 | 91 | -4 | 91 | 4 | 91 | -9 | 50 | -46 | 85 | -14 | 85 | -15 | 79 | 7 | 92 | 8 | 84 | -1 | 82 | 19 |

| | | | | | | | | | | | | | | | | | | | | | | | | | | | | | | | | |
|---|---|---|---|---|---|---|---|---|---|---|---|---|---|---|---|---|---|---|---|---|---|---|---|---|---|---|---|---|---|---|---|---|
| **CAN2** | 92 | 4 | 91 | -3 | 88 | -12 | 84 | -8 | 93 | 0 | 78 | -9 | 86 | -9 | 91 | 1 | 92 | 16 | 32 | -61 | 85 | -15 | 82 | -14 | 64 | -2 | 94 | 0 | 69 | -15 | 76 | -7 |
| **CAN20** | 92 | 2 | 91 | -3 | 58 | 28 | 95 | 5 | 87 | -5 | 33 | 19 | 72 | -24 | 91 | 0 | 88 | -12 | 79 | -21 | 86 | 28 | 82 | -13 | 91 | 12 | 91 | 4 | 81 | 16 | 85 | 10 |
| **CAN200** | 85 | 24 | 91 | 6 | 63 | 24 | 94 | 4 | 93 | -7 | 89 | -11 | 79 | 7 | 91 | 2 | 91 | -9 | 66 | -22 | 85 | -15 | 79 | 3 | 78 | -10 | 95 | 5 | 83 | -1 | 79 | -15 |
| **CAN201** | 92 | -2 | 91 | -9 | 60 | -20 | 92 | -7 | 93 | 0 | 78 | -11 | 78 | -22 | 91 | -9 | 67 | -17 | 77 | -11 | 85 | -13 | 55 | -39 | 86 | -14 | 89 | 2 | 56 | -31 | 74 | -21 |
| **CAN202** | 68 | -32 | 91 | -2 | 85 | -15 | 85 | 2 | 93 | 2 | 87 | -9 | 80 | -20 | 91 | -7 | 91 | -9 | 86 | -8 | 86 | -12 | 78 | -21 | 71 | -27 | 86 | -8 | 73 | 18 | 84 | -13 |
| **CAN203** | 92 | 0 | 91 | -9 | 90 | -5 | 94 | 15 | 93 | 8 | 87 | 2 | 93 | -7 | 91 | 3 | 91 | -9 | 90 | -3 | 86 | -11 | 83 | 0 | 89 | 2 | 91 | 13 | 86 | 9 | 85 | 2 |
| **CAN204** | 92 | -2 | 91 | 2 | 69 | -4 | 92 | -6 | 93 | -7 | 70 | -1 | 88 | 7 | 91 | 2 | 91 | -9 | 72 | -25 | 85 | -11 | 85 | 0 | 70 | -6 | 87 | 44 | 75 | -18 | 70 | 1 |
| **CAN205** | 92 | 5 | 91 | -9 | 85 | 3 | 96 | 2 | 93 | -7 | 81 | 7 | 90 | -2 | 91 | 0 | 91 | -9 | 77 | -3 | 86 | -6 | 86 | 3 | 93 | -1 | 96 | 1 | 86 | 14 | 85 | 1 |
| **CAN21** | 92 | 4 | 79 | 67 | 60 | 37 | 92 | -8 | 93 | -7 | 88 | 13 | 78 | 14 | 91 | 2 | 91 | -9 | 93 | 7 | 85 | -15 | 71 | 41 | 93 | 4 | 62 | 30 | 88 | 2 | 85 | 13 |
| **CAN23** | 92 | -4 | 91 | -7 | 68 | 19 | 93 | -2 | 88 | 12 | 65 | -24 | 86 | -10 | 91 | -7 | 91 | 3 | 92 | 14 | 86 | -10 | 86 | -5 | 88 | -11 | 89 | -4 | 80 | -20 | 83 | -5 |
| **CAN24** | 92 | 0 | 91 | 0 | 88 | -12 | 79 | -5 | 93 | 24 | 84 | -1 | 89 | -9 | 91 | -1 | 91 | -9 | 87 | -13 | 85 | -15 | 83 | -17 | 83 | 47 | 93 | 23 | 81 | 9 | 88 | 8 |
| **CAN25** | 92 | -8 | 91 | -6 | 72 | -28 | 94 | -3 | 93 | -4 | 94 | -6 | 88 | -10 | 91 | -9 | 91 | -9 | 73 | 43 | 85 | -15 | 84 | -7 | 94 | -4 | 92 | -5 | 83 | -17 | 85 | -15 |
| **CAN26** | 92 | 2 | 91 | -9 | 79 | -13 | 94 | -2 | 93 | 12 | 83 | -4 | 84 | -16 | 91 | -1 | 91 | 1 | 88 | -12 | 86 | 1 | 79 | -21 | 94 | 3 | 85 | 6 | 86 | 6 | 53 | -35 |
| **CAN27** | 74 | 37 | 91 | -9 | 71 | -15 | 94 | 6 | 93 | -3 | 82 | 1 | 87 | -9 | 91 | -3 | 85 | -1 | 60 | 52 | 86 | -6 | 78 | -17 | 89 | -4 | 91 | 1 | 75 | 58 | 85 | 2 |
| **CAN28** | 92 | -1 | 91 | -4 | 88 | 8 | 95 | -5 | 93 | -7 | 86 | 1 | 95 | -5 | 91 | -6 | 91 | 13 | 74 | 9 | 86 | 5 | 82 | -10 | 86 | -8 | 97 | -3 | 86 | 2 | 85 | -13 |
| **CAN29** | 92 | 2 | 91 | -6 | 96 | -4 | 85 | 1 | 93 | 1 | 66 | -5 | 41 | 6 | 91 | -1 | 54 | 10 | 27 | 5 | 85 | -15 | 28 | -2 | 90 | 3 | 86 | -5 | 64 | -16 | 83 | 11 |
| **CAN3** | 92 | -2 | 91 | -6 | 80 | -17 | 93 | -7 | 81 | -10 | 37 | -51 | 86 | -14 | 91 | 0 | 91 | -2 | 77 | -21 | 86 | -11 | 83 | -9 | 91 | 12 | 89 | 17 | 60 | -16 | 85 | -4 |
| **CAN30** | 84 | -9 | 91 | 2 | 82 | -7 | 78 | 1 | 93 | 19 | 82 | 6 | 96 | 8 | 91 | -5 | 91 | 1 | 37 | 0 | 86 | -6 | 87 | -2 | 86 | -7 | 79 | 10 | 79 | 13 | 86 | 9 |
| **CAN31** | 92 | -8 | 91 | -9 | 84 | 17 | 83 | -9 | 93 | -4 | 84 | 3 | 72 | -18 | 91 | 0 | 91 | 9 | 78 | -1 | 85 | -3 | 87 | -5 | 60 | -26 | 94 | -4 | 69 | -17 | 85 | 8 |
| **CAN32** | 84 | -7 | 91 | -9 | 82 | 9 | 76 | -22 | 93 | 0 | 83 | 15 | 95 | -5 | 91 | 12 | 92 | 12 | 42 | 20 | 86 | -9 | 86 | -14 | 90 | 15 | 79 | -17 | 75 | -25 | 85 | 17 |
| **CAN33** | 92 | 9 | 91 | 3 | 90 | 26 | 93 | 3 | 93 | -2 | 75 | -13 | 39 | 15 | 91 | 55 | 38 | 17 | 26 | 9 | 86 | 3 | 45 | 22 | 89 | -7 | 86 | -4 | 71 | -2 | 82 | -13 |
| **CAN34** | 85 | -2 | 91 | -8 | 79 | -4 | 80 | 1 | 93 | 22 | 82 | -6 | 95 | -5 | 91 | -4 | 91 | -2 | 42 | 12 | 86 | -5 | 86 | -14 | 87 | -6 | 75 | 5 | 75 | 1 | 85 | -4 |
| **CAN35** | 92 | 0 | 91 | 32 | 86 | -5 | 93 | 33 | 88 | -7 | 69 | 45 | 73 | -12 | 91 | -6 | 91 | -2 | 84 | -16 | 86 | 21 | 35 | 25 | 65 | 29 | 88 | 28 | 69 | -18 | 72 | 21 |
| **CAN36** | 92 | 0 | 91 | 0 | 95 | 16 | 85 | -6 | 93 | 2 | 87 | 6 | 88 | -2 | 91 | -6 | 91 | -4 | 89 | 7 | 86 | 8 | 87 | -7 | 93 | 0 | 91 | 8 | 81 | 5 | 85 | 1 |
| **CAN37** | 92 | -2 | 91 | -9 | 70 | -30 | 94 | 3 | 88 | 9 | 85 | -7 | 74 | 32 | 91 | -2 | 91 | -9 | 87 | -13 | 86 | -14 | 82 | -15 | 93 | 4 | 90 | 15 | 85 | 10 | 85 | -3 |
| **CAN38** | 92 | -7 | 91 | 14 | 93 | -7 | 82 | 12 | 93 | 9 | 88 | -9 | 86 | 13 | 91 | -3 | 91 | -6 | 86 | -11 | 86 | -10 | 83 | -12 | 93 | -1 | 89 | 13 | 55 | -28 | 85 | -12 |
| **CAN39** | 92 | -8 | 91 | -9 | 92 | -8 | 81 | 1 | 93 | -7 | 89 | 5 | 84 | -12 | 91 | -7 | 91 | -9 | 93 | 9 | 86 | 6 | 83 | -12 | 92 | -7 | 92 | -8 | 72 | -26 | 85 | 0 |
| **CAN4** | 92 | 12 | 91 | 1 | 89 | -11 | 90 | -4 | 87 | -10 | 67 | 40 | 48 | 8 | 91 | -4 | 91 | -1 | 55 | -45 | 86 | 27 | 73 | 3 | 52 | 36 | 91 | -6 | 56 | -7 | 63 | 38 |
| **CAN40** | 92 | 2 | 91 | -3 | 91 | 13 | 58 | -17 | 93 | -5 | 82 | 2 | 78 | -17 | 91 | 0 | 92 | 9 | 60 | 30 | 86 | 59 | 85 | -15 | 92 | 6 | 86 | -13 | 81 | -19 | 85 | 6 |
| **CAN41** | 92 | -8 | 75 | 54 | 65 | 43 | 96 | -4 | 93 | -7 | 76 | 64 | 65 | 50 | 91 | 9 | 65 | 57 | 93 | 12 | 86 | 14 | 71 | 59 | 90 | 35 | 92 | -8 | 88 | -12 | 89 | 35 |
| **CAN43** | 92 | 11 | 91 | -4 | 87 | 11 | 67 | -28 | 93 | 5 | 81 | 5 | 94 | -6 | 91 | 1 | 91 | -1 | 68 | -32 | 85 | -6 | 80 | -20 | 55 | -29 | 86 | -1 | 85 | 8 | 79 | -1 |
| **CAN44** | 84 | -8 | 91 | 5 | 83 | 5 | 85 | -4 | 93 | -6 | 82 | 3 | 97 | 23 | 91 | 1 | 91 | 1 | 50 | -29 | 86 | -1 | 88 | 18 | 82 | -8 | 66 | -22 | 74 | -5 | 80 | 0 |
| **CAN45** | 92 | 2 | 91 | 0 | 98 | 7 | 92 | -2 | 93 | 2 | 32 | -1 | 41 | 0 | 91 | 2 | 38 | 13 | 31 | 8 | 85 | -9 | 28 | 7 | 83 | 0 | 77 | -6 | 70 | -6 | 82 | 1 |
| **CAN46** | 25 | 12 | 91 | -7 | 85 | -15 | 93 | -7 | 93 | -7 | 81 | 15 | 73 | -18 | 91 | -9 | 91 | -3 | 68 | -18 | 86 | -5 | 83 | -8 | 92 | 5 | 96 | -3 | 83 | -15 | 87 | 18 |
| **CAN47** | 92 | -5 | 91 | 2 | 54 | 41 | 93 | 8 | 88 | 10 | 30 | 5 | 88 | 1 | 91 | 2 | 91 | 12 | 89 | 10 | 86 | 70 | 87 | 1 | 88 | -3 | 90 | 15 | 80 | 13 | 84 | -9 |
| **CAN48** | 92 | -7 | 91 | -9 | 96 | 28 | 82 | -18 | 93 | -7 | 84 | 11 | 86 | 0 | 91 | -9 | 91 | 1 | 42 | 26 | 86 | -1 | 83 | -13 | 67 | 23 | 87 | -13 | 72 | -24 | 80 | 11 |
| **CAN49** | 92 | -3 | 91 | -4 | 86 | 29 | 90 | -6 | 93 | -3 | 52 | -11 | 42 | -17 | 91 | 7 | 38 | 16 | 35 | 17 | 86 | 24 | 38 | -8 | 83 | 4 | 85 | -9 | 72 | -23 | 77 | 3 |
| **CAN5** | 92 | -5 | 91 | -9 | 89 | -11 | 94 | -6 | 87 | -13 | 41 | 29 | 90 | 43 | 91 | -4 | 91 | 2 | 88 | 0 | 85 | -15 | 62 | 34 | 80 | -7 | 91 | -9 | 79 | -11 | 85 | 5 |
| **CAN50** | 92 | -8 | 91 | -9 | 82 | -15 | 95 | -5 | 93 | -7 | 84 | -16 | 95 | -3 | 91 | 3 | 92 | 18 | 69 | -9 | 86 | -12 | 83 | 1 | 89 | 8 | 95 | -5 | 84 | -16 | 85 | -9 |
| **CAN51** | 92 | -1 | 91 | -6 | 90 | 15 | 95 | -1 | 93 | 24 | 64 | -21 | 57 | -32 | 91 | -9 | 33 | -38 | 24 | 1 | 86 | 19 | 28 | 9 | 81 | -14 | 82 | 17 | 73 | 10 | 85 | -6 |
| **CAN52** | 92 | -7 | 91 | -6 | 91 | -6 | 39 | -28 | 93 | -3 | 89 | -8 | 75 | -21 | 91 | -7 | 91 | -9 | 74 | -26 | 86 | 25 | 81 | -19 | 91 | -6 | 87 | -6 | 81 | -14 | 85 | -13 |
| **CAN53** | 92 | -6 | 91 | -5 | 72 | -15 | 80 | -20 | 93 | 3 | 89 | -3 | 90 | -10 | 91 | -9 | 91 | -3 | 77 | -13 | 86 | -2 | 77 | -18 | 91 | -5 | 90 | 3 | 60 | -11 | 88 | 2 |
| **CAN54** | 92 | -1 | 91 | -1 | 93 | -7 | 82 | 12 | 93 | -3 | 89 | 1 | 86 | 6 | 91 | -7 | 91 | -9 | 87 | -7 | 86 | -14 | 83 | 11 | 92 | -5 | 95 | 4 | 79 | 5 | 85 | -3 |
| **CAN55** | 55 | -28 | 91 | 3 | 60 | 30 | 85 | -2 | 93 | 12 | 85 | -15 | 84 | -16 | 91 | -6 | 91 | -1 | 66 | -17 | 86 | 60 | 74 | 26 | 92 | -4 | 89 | 20 | 51 | -5 | 85 | -10 |
| **CAN56** | 92 | -8 | 91 | -1 | 74 | -8 | 91 | -9 | 93 | -7 | 84 | -11 | 84 | 1 | 91 | -9 | 91 | -2 | 88 | -7 | 86 | -9 | 84 | 5 | 71 | -29 | 93 | -5 | 83 | -17 | 83 | -17 |
| **CAN57** | 84 | -7 | 91 | 0 | 82 | 6 | 85 | 4 | 93 | 16 | 80 | -6 | 92 | -6 | 91 | -6 | 91 | 7 | 62 | -16 | 86 | 1 | 83 | -14 | 54 | -8 | 75 | 0 | 77 | 1 | 58 | -31 |
| **CAN58** | 84 | -4 | 91 | 0 | 84 | -16 | 94 | -6 | 93 | -7 | 76 | 7 | 92 | 17 | 91 | -2 | 87 | -13 | 52 | -33 | 86 | -14 | 85 | 8 | 85 | -4 | 90 | -9 | 86 | -2 | 88 | 26 |
| **CAN59** | 92 | 4 | 91 | -5 | 83 | -6 | 88 | -4 | 93 | -5 | 84 | -1 | 69 | -23 | 91 | 0 | 86 | -2 | 39 | 29 | 86 | -4 | 66 | -30 | 66 | 53 | 86 | -7 | 90 | 14 | 81 | -15 |
| **CAN6** | 92 | -8 | 91 | -9 | 86 | -14 | 94 | 0 | 93 | -4 | 84 | 13 | 84 | -15 | 91 | 0 | 91 | -2 | 85 | 5 | 86 | -9 | 83 | -12 | 94 | -1 | 94 | -1 | 80 | -14 | 86 | 10 |
| **CAN60** | 92 | -4 | 91 | 4 | 96 | -4 | 93 | 12 | 93 | 1 | 70 | 39 | 39 | 10 | 91 | 1 | 58 | 30 | 35 | 15 | 85 | -15 | 30 | 7 | 81 | -9 | 87 | 4 | 82 | 12 | 82 | -16 |
| **CAN61** | 92 | 9 | 91 | 16 | 77 | 0 | 89 | 16 | 93 | 13 | 80 | 4 | 94 | 14 | 91 | -2 | 91 | -4 | 60 | 47 | 86 | -4 | 83 | 4 | 62 | 14 | 82 | 25 | 77 | 25 | 78 | 0 |
| **CAN62** | 92 | 2 | 91 | -3 | 80 | 49 | 82 | -13 | 93 | -7 | 83 | -1 | 94 | 14 | 91 | 4 | 92 | 29 | 87 | 31 | 86 | 34 | 82 | 4 | 92 | 0 | 85 | -11 | 66 | 56 | 89 | 15 |
| **CAN63** | 85 | 9 | 91 | -8 | 89 | -11 | 78 | -13 | 93 | 12 | 83 | 11 | 95 | -5 | 91 | 0 | 91 | -9 | 36 | -5 | 86 | -14 | 86 | -14 | 89 | 9 | 73 | -8 | 74 | -6 | 85 | 14 |

| | | | | | | | | | | | | | | | | | | | | | | | | | | | | | | | | |
|---|---|---|---|---|---|---|---|---|---|---|---|---|---|---|---|---|---|---|---|---|---|---|---|---|---|---|---|---|---|---|---|---|
| **CAN64** | 92 | -3 | 91 | -2 | 75 | -18 | 93 | -4 | 93 | -7 | 81 | -7 | 94 | 13 | 91 | -2 | 91 | -1 | 52 | 18 | 86 | -14 | 86 | 4 | 52 | 9 | 94 | -2 | 69 | -27 | 74 | 31 |
| **CAN65** | 84 | -7 | 91 | 1 | 85 | 3 | 94 | -6 | 93 | -2 | 88 | 2 | 81 | -8 | 91 | 36 | 87 | -13 | 61 | 20 | 86 | 37 | 84 | 7 | 85 | -3 | 81 | -4 | 76 | -7 | 86 | 1 |
| **CAN66** | 23 | 4 | 91 | 3 | 85 | -14 | 95 | 11 | 93 | -7 | 80 | 4 | 76 | -2 | 91 | -3 | 91 | -9 | 74 | 0 | 86 | -14 | 84 | 13 | 90 | 1 | 98 | 6 | 84 | 0 | 86 | 7 |
| **CAN67** | 92 | 28 | 91 | 7 | 69 | -23 | 93 | -1 | 93 | -1 | 76 | -23 | 82 | -5 | 91 | -1 | 90 | 1 | 87 | -4 | 86 | -11 | 73 | -19 | 86 | -5 | 93 | 3 | 67 | -4 | 77 | -16 |
| **CAN68** | 92 | 26 | 91 | 4 | 97 | 3 | 95 | 10 | 93 | 2 | 73 | -18 | 61 | -20 | 91 | -7 | 46 | -48 | 21 | -78 | 86 | 60 | 34 | -45 | 81 | -16 | 83 | 0 | 79 | -2 | 85 | -5 |
| **CAN69** | 92 | -3 | 91 | -1 | 94 | -6 | 85 | 3 | 69 | -10 | 86 | -7 | 90 | -1 | 91 | -1 | 91 | -9 | 92 | -4 | 85 | -15 | 82 | -12 | 89 | 1 | 77 | 3 | 83 | 6 | 85 | 4 |
| **CAN7** | 92 | -8 | 91 | -9 | 81 | -19 | 94 | -6 | 93 | -2 | 84 | 34 | 90 | -1 | 91 | -6 | 91 | -9 | 87 | -13 | 85 | -15 | 82 | -18 | 92 | 0 | 92 | -8 | 87 | 1 | 85 | 6 |
| **CAN70** | 92 | -8 | 91 | 6 | 89 | -11 | 18 | -7 | 93 | -1 | 88 | -12 | 79 | 10 | 91 | 1 | 91 | -6 | 78 | -1 | 86 | -14 | 83 | 16 | 92 | 5 | 89 | 8 | 84 | 13 | 85 | -10 |
| **CAN71** | 92 | 9 | 91 | 1 | 80 | -20 | 85 | -4 | 93 | 1 | 70 | 44 | 39 | -2 | 91 | 1 | 51 | -1 | 30 | 0 | 85 | -15 | 29 | 7 | 91 | 9 | 94 | 8 | 77 | 16 | 83 | 3 |
| **CAN72** | 92 | 5 | 91 | 7 | 84 | -16 | 82 | 19 | 93 | 0 | 75 | -6 | 93 | 26 | 91 | -2 | 91 | -9 | 76 | -8 | 85 | -15 | 83 | 23 | 65 | -26 | 92 | 14 | 74 | 5 | 72 | -18 |
| **CAN73** | 92 | -2 | 91 | -8 | 91 | -9 | 93 | 9 | 93 | 13 | 34 | 4 | 40 | -5 | 91 | 0 | 36 | 0 | 26 | -6 | 85 | -15 | 27 | 0 | 84 | 14 | 77 | 1 | 71 | -1 | 82 | -3 |
| **CAN74** | 85 | 14 | 91 | -9 | 85 | 7 | 93 | -7 | 93 | -6 | 82 | 6 | 80 | -3 | 91 | -3 | 91 | -9 | 41 | 16 | 86 | -15 | 85 | -7 | 47 | 4 | 89 | -10 | 89 | -7 | 35 | 1 |
| **CAN75** | 92 | 0 | 91 | -8 | 52 | 8 | 92 | -5 | 59 | -29 | 76 | -12 | 85 | -15 | 91 | -2 | 91 | -9 | 79 | -4 | 86 | -14 | 77 | -23 | 88 | -4 | 92 | -4 | 78 | -13 | 85 | -3 |
| **CAN76** | 92 | -8 | 91 | 13 | 89 | -11 | 23 | 18 | 93 | 0 | 90 | 5 | 81 | 17 | 91 | -8 | 91 | -9 | 90 | 74 | 86 | -14 | 84 | 17 | 91 | -4 | 90 | 13 | 84 | 13 | 85 | -4 |
| **CAN77** | 92 | 16 | 80 | 8 | 97 | -3 | 87 | 27 | 93 | 6 | 50 | -1 | 65 | -3 | 91 | -5 | 91 | 26 | 37 | 5 | 86 | -14 | 61 | 2 | 47 | 5 | 85 | 18 | 60 | 10 | 80 | -5 |
| **CAN78** | 92 | 2 | 91 | -7 | 65 | -35 | 94 | -6 | 88 | 7 | 66 | -21 | 82 | -8 | 91 | 0 | 91 | -9 | 80 | -17 | 85 | -15 | 84 | -4 | 92 | 14 | 87 | -12 | 75 | -22 | 83 | 8 |
| **CAN79** | 92 | -1 | 91 | 5 | 87 | -13 | 92 | -6 | 93 | 2 | 66 | -11 | 34 | 3 | 91 | 0 | 37 | 9 | 23 | -3 | 85 | -15 | 27 | 1 | 84 | -6 | 87 | 2 | 84 | 3 | 82 | -7 |
| **CAN8** | 92 | -6 | 91 | -9 | 98 | 30 | 84 | -16 | 87 | -13 | 52 | 30 | 73 | -15 | 91 | 8 | 91 | 6 | 85 | -15 | 86 | 5 | 34 | -7 | 61 | 38 | 89 | -11 | 82 | -12 | 73 | 32 |
| **CAN80** | 84 | -9 | 91 | 2 | 90 | -11 | 96 | 5 | 93 | -7 | 79 | -11 | 83 | 21 | 91 | -6 | 91 | -9 | 58 | 20 | 86 | -14 | 86 | 12 | 40 | -49 | 89 | -2 | 78 | 42 | 71 | -11 |
| **CAN81** | 92 | -3 | 91 | 3 | 95 | -5 | 84 | 16 | 93 | 16 | 89 | 0 | 84 | -2 | 91 | -9 | 83 | -17 | 83 | -8 | 86 | -14 | 80 | -5 | 92 | -5 | 96 | 24 | 91 | 26 | 84 | -11 |
| **CAN82** | 92 | 7 | 91 | -3 | 50 | -6 | 92 | -8 | 74 | 23 | 54 | -15 | 94 | 4 | 91 | -2 | 91 | -9 | 80 | -6 | 86 | -14 | 50 | -35 | 72 | -13 | 84 | -16 | 77 | -23 | 65 | -7 |
| **CAN83** | 92 | -7 | 91 | 13 | 48 | 12 | 92 | -5 | 72 | 42 | 83 | -16 | 88 | 1 | 91 | -7 | 91 | 0 | 79 | -17 | 85 | -8 | 86 | -9 | 88 | -6 | 93 | -5 | 47 | 1 | 84 | -12 |
| **CAN84** | 92 | -7 | 91 | -2 | 95 | -5 | 92 | -6 | 67 | -30 | 86 | -5 | 90 | -1 | 91 | -1 | 91 | -9 | 92 | -2 | 86 | -14 | 82 | -13 | 89 | -3 | 71 | -21 | 82 | -10 | 85 | 0 |
| **CAN85** | 92 | -1 | 91 | -6 | 37 | -1 | 94 | -5 | 93 | -6 | 84 | 14 | 53 | -42 | 91 | 1 | 91 | -9 | 91 | -1 | 85 | -15 | 82 | -15 | 90 | 9 | 97 | 5 | 50 | 12 | 85 | 20 |
| **CAN86** | 92 | -6 | 91 | -9 | 93 | 6 | 87 | 3 | 93 | -2 | 95 | 0 | 89 | -10 | 91 | -5 | 91 | 2 | 34 | 25 | 85 | -4 | 85 | -13 | 94 | -1 | 93 | -2 | 87 | -5 | 86 | -7 |
| **CAN88** | 92 | 0 | 91 | 8 | 94 | -6 | 86 | 14 | 74 | 25 | 87 | -2 | 91 | 6 | 91 | 0 | 91 | -9 | 93 | -2 | 85 | -15 | 87 | 18 | 88 | -4 | 82 | 21 | 87 | 30 | 85 | -6 |
| **CAN89** | 92 | 2 | 91 | -1 | 95 | -5 | 79 | -7 | 93 | -5 | 89 | 6 | 83 | -4 | 91 | -6 | 82 | -18 | 84 | -3 | 85 | -15 | 80 | -5 | 93 | -1 | 87 | -11 | 80 | -20 | 85 | -5 |
| **CAN9** | 92 | 10 | 91 | -4 | 78 | -9 | 92 | -8 | 93 | -7 | 82 | -1 | 82 | -18 | 91 | -5 | 92 | 8 | 78 | 1 | 85 | -10 | 84 | -15 | 58 | -27 | 91 | -9 | 82 | -11 | 83 | 3 |
| **CAN90** | 92 | 10 | 91 | -9 | 55 | 21 | 93 | -1 | 77 | 50 | 71 | 5 | 91 | -9 | 91 | 0 | 91 | -9 | 86 | -5 | 85 | -15 | 69 | -31 | 81 | 0 | 88 | 10 | 79 | 4 | 86 | 17 |
| **CAN91** | 84 | -7 | 91 | -5 | 67 | 47 | 93 | 9 | 93 | -1 | 85 | -1 | 88 | 0 | 91 | 0 | 91 | 0 | 60 | 44 | 85 | -15 | 78 | -11 | 55 | 27 | 92 | 6 | 89 | 0 | 58 | 41 |
| **CAN92** | 92 | -8 | 80 | 15 | 96 | -4 | 84 | -5 | 93 | -6 | 46 | -27 | 68 | 6 | 91 | -1 | 91 | -9 | 35 | -3 | 86 | -14 | 52 | -24 | 35 | -47 | 79 | -8 | 58 | 0 | 82 | 4 |
| **CAN93** | 92 | -5 | 91 | -9 | 73 | -27 | 70 | 24 | 93 | 14 | 86 | 2 | 95 | 2 | 91 | 6 | 91 | -9 | 75 | -14 | 86 | -14 | 76 | -24 | 84 | 1 | 77 | -1 | 79 | 8 | 74 | -3 |
| **CAN94** | 85 | 12 | 91 | -4 | 88 | -12 | 80 | 12 | 93 | 19 | 82 | -15 | 95 | -5 | 91 | 3 | 91 | -9 | 47 | -41 | 85 | -15 | 87 | 4 | 89 | 7 | 77 | 11 | 71 | 11 | 84 | -5 |
| **CAN95** | 92 | -2 | 91 | -9 | 96 | -4 | 92 | -8 | 93 | -7 | 79 | 9 | 32 | -25 | 91 | -7 | 91 | -9 | 48 | -10 | 86 | -7 | 65 | -5 | 88 | -10 | 86 | -14 | 74 | 7 | 84 | 22 |
| **CAN96** | 84 | -2 | 91 | 7 | 90 | -10 | 94 | 2 | 93 | 0 | 84 | 2 | 91 | 2 | 91 | -5 | 91 | -9 | 40 | 13 | 86 | -14 | 86 | 6 | 66 | 38 | 94 | 13 | 79 | 4 | 82 | 9 |
| **CAN97** | 92 | -3 | 91 | 2 | 86 | -14 | 86 | 24 | 88 | 9 | 81 | -7 | 97 | 7 | 91 | -2 | 91 | -9 | 54 | 27 | 86 | -15 | 82 | 12 | 83 | -7 | 90 | 16 | 78 | 4 | 84 | 0 |
| **CAN98** | 92 | -8 | 91 | 21 | 85 | 19 | 95 | 2 | 94 | 43 | 84 | -3 | 81 | 29 | 91 | 26 | 91 | 5 | 94 | 3 | 86 | 70 | 86 | 42 | 91 | -2 | 87 | 23 | 86 | -12 | 85 | 1 |
| **HB1** | 92 | -8 | 91 | 6 | 85 | -15 | 60 | 29 | 93 | -7 | 84 | -7 | 92 | 8 | 91 | -7 | 91 | -9 | 62 | 9 | 45 | 15 | 83 | 9 | 94 | -6 | 96 | 6 | 84 | 4 | 85 | -13 |
| **HB10** | 75 | 54 | 91 | -9 | 68 | 1 | 93 | -5 | 93 | -7 | 86 | 9 | 92 | -8 | 91 | 1 | 90 | 11 | 22 | 15 | 86 | 0 | 84 | 2 | 86 | 4 | 97 | -3 | 87 | 15 | 80 | 13 |
| **HB11** | 84 | -16 | 91 | 0 | 49 | 10 | 92 | -1 | 93 | 31 | 95 | -5 | 86 | 4 | 91 | -8 | 91 | -1 | 50 | 16 | 86 | -9 | 84 | -1 | 53 | 26 | 92 | 7 | 51 | 25 | 85 | -3 |
| **HB12** | 92 | -8 | 91 | 3 | 61 | -15 | 93 | -7 | 93 | -7 | 89 | -11 | 94 | -6 | 73 | -27 | 92 | 0 | 60 | 42 | 86 | -11 | 85 | -8 | 91 | -9 | 87 | 3 | 85 | 2 | 84 | -16 |
| **HB13** | 21 | -1 | 91 | 1 | 86 | 14 | 94 | -6 | 93 | -7 | 83 | -10 | 72 | -2 | 91 | -4 | 92 | -9 | 55 | 32 | 86 | -3 | 83 | -3 | 91 | -3 | 97 | -3 | 84 | -8 | 87 | -3 |
| **HB14** | 21 | 2 | 91 | 4 | 80 | -6 | 94 | -6 | 93 | -7 | 84 | 1 | 72 | -1 | 91 | -6 | 91 | -9 | 39 | -14 | 86 | 4 | 83 | 5 | 91 | -6 | 97 | -3 | 84 | -7 | 87 | -6 |
| **HB15** | 84 | -12 | 62 | -37 | 86 | 8 | 96 | -3 | 93 | -7 | 92 | 4 | 66 | -34 | 91 | -9 | 66 | -13 | 73 | -12 | 85 | -1 | 83 | -9 | 93 | -3 | 99 | -1 | 85 | -15 | 85 | -8 |
| **HB16** | 92 | 3 | 91 | -1 | 69 | 3 | 94 | -5 | 93 | -7 | 79 | -7 | 88 | -12 | 91 | -6 | 92 | 10 | 84 | -7 | 85 | 1 | 82 | -13 | 72 | 7 | 96 | 0 | 84 | -5 | 81 | -3 |
| **HB17** | 92 | 4 | 91 | 0 | 67 | -3 | 94 | -3 | 93 | -7 | 80 | 4 | 94 | 21 | 91 | 5 | 91 | 5 | 78 | -16 | 85 | -4 | 83 | 1 | 56 | 27 | 93 | -7 | 83 | -4 | 81 | 3 |
| **HB18** | 92 | -4 | 91 | -9 | 70 | -3 | 95 | -5 | 93 | -7 | 80 | -3 | 84 | -16 | 91 | -4 | 91 | 10 | 82 | -2 | 85 | -1 | 85 | 1 | 89 | -6 | 99 | -1 | 85 | -1 | 84 | -2 |
| **HB19** | 84 | -12 | 91 | 3 | 55 | 23 | 94 | 5 | 93 | -3 | 91 | 0 | 86 | 5 | 76 | 44 | 91 | -9 | 76 | -16 | 86 | -7 | 84 | -4 | 94 | -1 | 52 | 22 | 89 | 5 | 85 | -4 |
| **HB2** | 92 | -5 | 91 | 3 | 73 | 5 | 95 | -3 | 93 | -3 | 81 | 15 | 84 | -5 | 91 | -1 | 91 | 5 | 84 | -6 | 85 | -1 | 85 | -11 | 84 | 11 | 99 | 17 | 85 | 13 | 86 | 27 |
| **HB21** | 92 | 0 | 91 | -5 | 71 | 2 | 95 | -3 | 93 | -7 | 79 | -11 | 89 | -1 | 91 | -9 | 91 | 0 | 83 | 2 | 85 | 0 | 85 | 3 | 90 | 2 | 99 | 0 | 84 | -10 | 84 | -5 |
| **HB22** | 92 | 6 | 91 | -4 | 88 | 7 | 83 | -17 | 93 | -7 | 77 | 0 | 91 | -4 | 91 | -3 | 91 | 6 | 46 | 17 | 86 | 21 | 85 | -1 | 52 | 17 | 95 | -5 | 73 | 50 | 62 | -9 |
| **HB23** | 81 | 59 | 91 | -1 | 65 | 4 | 93 | -3 | 93 | -7 | 80 | 14 | 74 | -2 | 71 | 57 | 92 | 24 | 35 | 21 | 86 | 18 | 83 | -2 | 79 | -3 | 45 | 7 | 79 | -11 | 86 | 17 |
| **HB24** | 92 | -4 | 91 | 6 | 85 | 20 | 64 | 12 | 87 | -13 | 78 | -19 | 85 | -6 | 91 | -6 | 92 | 25 | 45 | 23 | 86 | 19 | 80 | -15 | 91 | -5 | 88 | -12 | 86 | -14 | 85 | -15 |

| | | | | | | | | | | | | | | | | | | | | | | | | | | | | | | | | |
|---|---|---|---|---|---|---|---|---|---|---|---|---|---|---|---|---|---|---|---|---|---|---|---|---|---|---|---|---|---|---|---|---|
| **HB25** | 59 | -41 | 91 | -4 | 80 | -20 | 94 | -2 | 93 | -1 | 83 | -17 | 84 | -14 | 91 | -4 | 91 | -8 | 89 | -3 | 86 | -10 | 83 | -15 | 91 | -3 | 79 | -14 | 86 | -8 | 85 | -10 |
| **HB26** | 85 | -8 | 91 | -4 | 87 | 5 | 94 | -6 | 93 | -7 | 88 | 16 | 78 | -9 | 73 | -25 | 91 | -6 | 67 | -28 | 85 | -8 | 84 | 3 | 89 | -8 | 92 | -8 | 84 | -16 | 87 | 18 |
| **HB27** | 92 | -8 | 91 | -7 | 56 | -11 | 96 | 6 | 93 | -1 | 84 | 13 | 86 | -9 | 91 | -3 | 76 | -5 | 90 | 7 | 85 | 2 | 85 | -5 | 82 | -1 | 92 | 9 | 61 | 52 | 67 | -4 |
| **HB28** | 92 | -7 | 91 | -4 | 70 | -6 | 96 | 1 | 93 | -7 | 79 | -21 | 88 | 12 | 91 | 16 | 91 | -7 | 83 | -13 | 86 | -2 | 86 | 12 | 83 | 10 | 92 | -8 | 84 | -1 | 85 | -4 |
| **HB29** | 92 | 3 | 91 | -9 | 79 | 29 | 91 | -9 | 93 | -7 | 80 | -9 | 87 | -13 | 73 | -22 | 91 | 26 | 67 | 9 | 86 | 28 | 83 | -5 | 73 | -19 | 87 | -13 | 86 | 9 | 80 | -10 |
| **HB3** | 92 | -8 | 91 | -2 | 69 | 2 | 85 | -15 | 93 | -7 | 84 | 12 | 95 | 15 | 91 | 3 | 80 | -2 | 55 | 51 | 86 | 0 | 79 | -4 | 89 | 6 | 98 | -2 | 89 | -3 | 76 | 14 |
| **HB30** | 92 | -1 | 91 | 7 | 75 | -5 | 95 | 4 | 88 | -9 | 82 | -1 | 80 | 10 | 91 | 66 | 88 | -2 | 34 | 17 | 86 | 8 | 84 | 7 | 82 | 1 | 75 | 24 | 82 | 3 | 74 | 58 |
| **HB35** | 80 | 23 | 91 | 1 | 45 | 23 | 94 | -6 | 93 | -1 | 84 | 3 | 95 | -4 | 61 | 44 | 91 | 15 | 26 | 9 | 86 | 8 | 88 | 1 | 66 | 10 | 56 | 9 | 89 | 8 | 51 | -14 |
| **HB5** | 84 | 57 | 91 | -9 | 91 | 19 | 92 | 6 | 93 | -4 | 90 | 4 | 95 | -5 | 91 | -5 | 92 | -5 | 78 | -13 | 86 | 9 | 85 | 5 | 94 | 5 | 85 | -3 | 71 | 53 | 85 | -4 |
| **HB6** | 63 | -35 | 91 | 0 | 74 | -6 | 86 | -1 | 93 | 2 | 89 | -7 | 84 | 26 | 91 | -2 | 92 | 8 | 70 | -11 | 85 | -3 | 86 | -11 | 82 | -12 | 83 | -3 | 79 | 2 | 77 | -23 |
| **HB7** | 92 | 2 | 91 | -3 | 74 | -13 | 92 | -8 | 93 | 3 | 82 | -5 | 86 | 0 | 71 | -22 | 91 | -9 | 87 | -5 | 86 | -5 | 80 | 0 | 93 | -3 | 86 | -14 | 88 | -11 | 87 | 1 |
| **HB8** | 92 | 16 | 91 | -2 | 68 | -2 | 89 | 59 | 93 | -7 | 82 | -4 | 91 | -9 | 91 | 9 | 81 | 5 | 30 | -29 | 45 | 22 | 77 | -8 | 88 | 1 | 98 | -2 | 89 | -2 | 69 | -9 |
| **HB9** | 92 | 6 | 91 | 2 | 82 | 2 | 94 | 3 | 93 | -7 | 84 | 6 | 92 | 19 | 91 | 2 | 91 | 9 | 84 | -16 | 85 | 2 | 83 | 2 | 88 | 2 | 89 | -4 | 89 | 7 | 85 | 9 |
| **HU1** | 92 | 0 | 91 | 7 | 62 | -11 | 93 | -7 | 93 | -7 | 79 | -8 | 79 | 2 | 72 | -28 | 92 | 14 | 44 | -14 | 86 | -14 | 85 | 9 | 76 | -16 | 82 | -18 | 79 | -9 | 85 | -5 |
| **HU11** | 66 | -26 | 91 | 10 | 72 | 1 | 90 | 17 | 93 | 17 | 92 | 2 | 91 | 13 | 72 | -22 | 92 | 16 | 33 | 19 | 86 | 9 | 85 | 9 | 84 | -6 | 80 | 18 | 64 | 9 | 79 | -7 |
| **HU12** | 92 | -6 | 91 | 1 | 59 | -33 | 94 | -6 | 93 | -7 | 85 | 3 | 85 | 2 | 91 | -9 | 92 | -1 | 72 | -18 | 86 | -9 | 84 | 6 | 93 | -5 | 92 | -8 | 88 | -12 | 85 | 1 |
| **HU13** | 92 | 1 | 91 | 4 | 68 | -11 | 92 | -1 | 75 | 3 | 78 | -10 | 90 | 2 | 91 | -5 | 92 | -1 | 54 | 36 | 85 | -7 | 88 | 0 | 68 | 56 | 94 | 17 | 88 | 63 | 55 | 30 |
| **HU14** | 80 | 31 | 91 | 22 | 76 | -6 | 85 | 12 | 93 | 2 | 84 | -6 | 90 | 38 | 62 | 42 | 80 | -11 | 42 | 23 | 42 | -51 | 79 | 19 | 92 | -2 | 85 | 9 | 55 | -7 | 85 | -8 |
| **HU17** | 92 | 0 | 91 | -1 | 68 | -15 | 95 | -5 | 93 | -7 | 86 | -1 | 94 | -3 | 91 | -1 | 91 | 2 | 37 | 28 | 85 | -3 | 85 | -6 | 87 | -3 | 97 | -3 | 84 | -13 | 85 | -1 |
| **HU18** | 52 | -48 | 91 | 15 | 68 | 19 | 95 | 21 | 93 | 5 | 79 | -16 | 81 | 19 | 47 | -45 | 91 | -1 | 31 | 9 | 85 | -8 | 84 | 21 | 85 | 29 | 47 | 19 | 84 | 22 | 85 | -8 |
| **HU19** | 92 | -6 | 91 | -3 | 85 | 15 | 96 | 12 | 93 | 3 | 74 | -9 | 82 | -18 | 91 | -1 | 91 | -9 | 89 | -9 | 86 | -5 | 85 | 1 | 93 | -1 | 94 | 6 | 89 | 16 | 85 | 8 |
| **HU2** | 92 | 0 | 91 | -5 | 73 | -2 | 94 | 4 | 93 | 7 | 86 | -1 | 92 | -6 | 91 | -6 | 91 | 8 | 89 | -3 | 85 | -12 | 84 | 2 | 78 | -15 | 91 | 19 | 89 | 28 | 85 | -10 |
| **HU21** | 23 | 2 | 91 | 4 | 88 | 12 | 96 | 4 | 93 | -7 | 84 | -16 | 82 | 5 | 91 | 8 | 91 | -1 | 54 | -23 | 86 | -7 | 84 | 11 | 90 | 10 | 94 | -4 | 84 | -5 | 84 | -13 |
| **HU22** | 74 | -19 | 91 | 3 | 83 | -11 | 79 | 11 | 93 | 10 | 91 | -1 | 95 | -1 | 91 | -2 | 91 | 0 | 62 | -16 | 86 | 10 | 87 | 1 | 80 | -10 | 73 | -3 | 71 | 4 | 75 | -12 |
| **HU23** | 20 | -4 | 91 | 14 | 87 | 3 | 97 | 18 | 93 | -7 | 80 | -1 | 76 | 21 | 91 | -1 | 91 | 2 | 57 | -16 | 86 | -1 | 83 | 11 | 92 | -1 | 97 | 5 | 87 | 20 | 86 | 6 |
| **HU25** | 92 | -5 | 91 | -2 | 88 | 12 | 79 | -21 | 93 | -7 | 83 | 15 | 80 | -20 | 91 | 2 | 91 | 1 | 61 | -31 | 86 | -13 | 80 | -6 | 68 | -1 | 77 | -17 | 48 | -27 | 81 | 17 |
| **HU26** | 92 | -2 | 91 | 0 | 85 | 5 | 95 | 24 | 93 | -5 | 81 | -9 | 75 | 39 | 91 | -1 | 92 | 5 | 53 | 10 | 86 | -2 | 85 | 4 | 66 | 39 | 96 | 4 | 64 | -19 | 85 | -4 |
| **HU27** | 92 | 4 | 91 | 5 | 72 | -13 | 73 | 5 | 93 | 8 | 83 | 10 | 72 | -9 | 91 | 1 | 91 | -2 | 25 | -7 | 86 | -1 | 84 | 12 | 77 | -4 | 64 | -3 | 81 | 24 | 55 | 1 |
| **HU28** | 83 | -6 | 91 | 20 | 84 | 6 | 84 | 15 | 93 | 5 | 84 | -6 | 83 | 13 | 91 | 2 | 91 | 14 | 92 | 16 | 86 | 4 | 84 | 11 | 77 | -11 | 85 | 14 | 90 | 18 | 87 | -5 |
| **HU3** | 91 | 72 | 91 | 8 | 70 | 42 | 81 | 45 | 93 | 22 | 84 | 45 | 91 | 13 | 79 | 54 | 91 | 3 | 72 | -21 | 86 | -14 | 87 | 17 | 94 | 1 | 86 | 55 | 88 | 7 | 85 | -13 |
| **HU30** | 82 | -18 | 91 | 9 | 77 | 7 | 96 | 11 | 93 | -6 | 80 | 5 | 91 | -3 | 91 | -5 | 91 | -5 | 72 | -24 | 86 | -12 | 83 | -2 | 91 | -2 | 70 | -23 | 87 | 3 | 86 | 11 |
| **HU31** | 92 | 3 | 91 | 11 | 80 | 5 | 79 | -3 | 93 | -4 | 82 | -4 | 81 | 16 | 91 | -2 | 91 | -9 | 39 | -1 | 86 | 7 | 84 | 9 | 91 | 1 | 85 | -1 | 60 | -17 | 85 | -1 |
| **HU32** | 22 | 7 | 91 | -9 | 76 | -15 | 95 | -1 | 93 | -7 | 80 | 6 | 94 | -6 | 91 | 3 | 91 | 4 | 56 | -1 | 86 | 25 | 83 | 0 | 90 | -7 | 92 | -8 | 86 | 9 | 86 | 14 |
| **HU35** | 92 | 6 | 91 | -1 | 62 | -2 | 77 | -19 | 93 | -7 | 88 | 8 | 92 | 8 | 91 | 65 | 91 | 3 | 70 | 20 | 86 | -6 | 83 | 16 | 67 | 34 | 53 | 20 | 85 | -12 | 74 | 27 |
| **HU37** | 92 | 1 | 91 | -4 | 46 | -28 | 94 | -3 | 93 | -7 | 82 | 2 | 89 | 10 | 91 | -7 | 71 | -14 | 90 | 0 | 86 | 0 | 83 | -2 | 78 | 46 | 95 | -5 | 84 | -6 | 86 | -4 |
| **HU39** | 84 | -5 | 91 | -3 | 60 | 26 | 94 | -2 | 93 | -2 | 91 | 7 | 76 | 65 | 91 | 1 | 91 | -3 | 34 | 19 | 86 | 3 | 83 | 37 | 66 | 46 | 83 | 8 | 90 | -1 | 69 | 42 |
| **HU40** | 84 | -13 | 91 | -8 | 64 | 47 | 84 | -11 | 93 | -7 | 86 | -7 | 85 | -15 | 77 | 32 | 91 | -9 | 78 | 2 | 85 | 9 | 84 | -4 | 74 | -26 | 52 | 31 | 84 | -7 | 78 | -22 |
| **HU41** | 84 | -1 | 91 | 2 | 88 | 11 | 85 | -15 | 93 | -7 | 83 | 18 | 80 | -3 | 91 | 16 | 91 | 4 | 79 | -20 | 86 | -2 | 81 | -2 | 85 | 18 | 94 | -6 | 81 | -19 | 74 | 5 |
| **HU42** | 92 | 6 | 91 | -3 | 87 | 10 | 94 | -4 | 93 | -7 | 85 | 1 | 89 | -11 | 56 | -40 | 91 | 2 | 62 | 37 | 86 | -6 | 73 | -19 | 70 | -22 | 93 | -7 | 83 | -16 | 66 | -25 |
| **HU43** | 84 | -2 | 91 | -6 | 57 | -22 | 93 | -5 | 93 | -7 | 89 | 2 | 94 | 12 | 91 | -3 | 91 | 1 | 39 | 3 | 86 | -9 | 82 | -5 | 71 | 1 | 91 | -9 | 82 | -11 | 68 | -6 |
| **HU44** | 84 | -8 | 91 | -8 | 88 | 3 | 77 | -9 | 93 | 6 | 82 | -8 | 93 | -7 | 91 | -2 | 91 | -6 | 56 | 33 | 86 | -7 | 87 | 2 | 87 | -5 | 70 | -13 | 76 | 5 | 85 | -2 |
| **HU45** | 92 | 7 | 91 | 4 | 72 | -3 | 95 | -5 | 93 | -7 | 80 | -1 | 88 | 14 | 91 | 1 | 91 | -8 | 75 | -21 | 86 | 0 | 85 | 5 | 86 | 1 | 96 | -4 | 84 | -2 | 63 | -18 |
| **HU46** | 88 | -2 | 91 | 0 | 79 | -3 | 95 | 7 | 88 | -6 | 87 | 9 | 93 | -1 | 91 | -2 | 91 | -9 | 53 | -31 | 85 | -9 | 83 | 2 | 86 | -8 | 78 | 1 | 87 | 18 | 62 | -17 |
| **HU47** | 87 | 52 | 91 | 5 | 75 | -15 | 90 | -6 | 93 | -7 | 82 | 4 | 86 | 7 | 91 | 41 | 91 | -9 | 45 | 18 | 86 | 2 | 84 | 9 | 93 | 2 | 80 | 40 | 83 | -11 | 74 | 56 |
| **HU5** | 85 | 12 | 91 | 17 | 86 | 1 | 95 | 0 | 93 | -3 | 86 | 0 | 83 | 6 | 91 | 0 | 91 | -2 | 76 | 53 | 85 | -12 | 84 | 11 | 94 | -3 | 70 | -11 | 74 | 6 | 76 | 41 |
| **HU50** | 92 | -6 | 91 | -9 | 94 | 14 | 41 | -21 | 93 | -7 | 84 | 7 | 86 | -14 | 91 | -9 | 91 | -4 | 61 | 5 | 47 | 25 | 82 | -10 | 93 | -7 | 91 | -9 | 80 | -13 | 85 | 2 |
| **HU52** | 92 | -6 | 91 | -6 | 80 | -18 | 88 | -6 | 93 | 6 | 78 | 20 | 90 | -10 | 91 | -4 | 91 | -5 | 62 | -30 | 86 | -10 | 85 | 6 | 62 | -29 | 88 | 2 | 76 | -13 | 77 | -17 |
| **HU53** | 86 | -4 | 91 | 6 | 76 | -5 | 93 | -7 | 93 | -7 | 85 | 4 | 78 | -3 | 91 | -5 | 67 | -33 | 16 | 5 | 86 | -13 | 67 | -14 | 88 | 33 | 91 | -9 | 85 | -15 | 75 | -11 |
| **HU54** | 92 | 10 | 91 | 1 | 71 | 0 | 93 | -2 | 93 | -7 | 80 | 11 | 74 | -26 | 78 | 29 | 91 | -9 | 54 | 11 | 86 | 10 | 83 | -7 | 78 | -9 | 93 | 5 | 83 | -2 | 86 | 14 |
| **HU55** | 92 | 10 | 91 | 1 | 89 | -6 | 78 | -17 | 88 | -12 | 81 | 9 | 90 | 15 | 91 | -1 | 91 | -5 | 52 | -6 | 86 | 12 | 83 | 1 | 53 | -39 | 93 | -7 | 74 | -6 | 78 | 4 |
| **HU57** | 92 | -8 | 91 | 2 | 93 | 7 | 94 | -5 | 87 | -13 | 83 | -17 | 81 | 2 | 91 | 1 | 91 | 0 | 84 | -16 | 85 | -5 | 76 | -2 | 87 | 0 | 95 | -1 | 84 | -3 | 84 | -12 |
| **HU58** | 70 | 56 | 91 | -7 | 67 | -16 | 95 | 14 | 93 | 19 | 87 | 0 | 81 | -19 | 91 | -3 | 91 | 1 | 64 | -28 | 85 | -7 | 83 | -1 | 74 | -20 | 84 | 7 | 67 | 28 | 73 | -18 |
| **HU59** | 86 | -6 | 91 | 9 | 76 | -2 | 93 | -7 | 93 | -7 | 84 | 0 | 97 | 25 | 91 | -9 | 88 | -2 | 18 | -1 | 86 | -6 | 84 | 7 | 78 | -17 | 93 | -3 | 84 | -8 | 84 | -5 |

| | | | | | | | | | | | | | | | | | | | | | | | | | | | | | | | | |
|---|---|---|---|---|---|---|---|---|---|---|---|---|---|---|---|---|---|---|---|---|---|---|---|---|---|---|---|---|---|---|---|---|
| **HU6** | 92 | 6 | 91 | -4 | 82 | 5 | 71 | -8 | 93 | -3 | 81 | 5 | 85 | 2 | 91 | -6 | 91 | 6 | 44 | 11 | 86 | -2 | 84 | -4 | 92 | 2 | 95 | -5 | 86 | -2 | 85 | 4 |
| **HU60** | 92 | 0 | 91 | 14 | 72 | -2 | 93 | 15 | 93 | 15 | 85 | -5 | 86 | 14 | 80 | 60 | 91 | 2 | 72 | -21 | 85 | -4 | 83 | 2 | 83 | -7 | 80 | 19 | 83 | 24 | 81 | -8 |
| **HU61** | 84 | -12 | 91 | 4 | 62 | 34 | 93 | 2 | 93 | -5 | 85 | 8 | 84 | 11 | 91 | -2 | 91 | 2 | 45 | 23 | 86 | -2 | 84 | 0 | 58 | -26 | 93 | -1 | 77 | -6 | 57 | 24 |
| **HU62** | 92 | 2 | 91 | -7 | 67 | -5 | 96 | -4 | 93 | -7 | 84 | 3 | 91 | -9 | 91 | -8 | 91 | -9 | 67 | -24 | 86 | -4 | 85 | -4 | 87 | -5 | 93 | -7 | 89 | 0 | 85 | 4 |
| **HU63** | 92 | -2 | 91 | -8 | 80 | 3 | 94 | -6 | 93 | 3 | 85 | 8 | 89 | -11 | 91 | -6 | 91 | -9 | 45 | 9 | 86 | 14 | 85 | -6 | 57 | -39 | 82 | 10 | 86 | 11 | 80 | 6 |
| **HU64** | 92 | -4 | 91 | -9 | 75 | -7 | 94 | -2 | 93 | 1 | 80 | -13 | 88 | -12 | 91 | -3 | 91 | 5 | 53 | -42 | 86 | -1 | 85 | -11 | 93 | -2 | 87 | -13 | 88 | -12 | 85 | -11 |
| **HU65** | 84 | -7 | 91 | 19 | 73 | 8 | 56 | 2 | 93 | -7 | 84 | 0 | 76 | 22 | 91 | 0 | 87 | 55 | 46 | 29 | 62 | 59 | 87 | 54 | 77 | 53 | 87 | 3 | 76 | 11 | 61 | 33 |
| **HU66** | 92 | 1 | 91 | 10 | 73 | -2 | 94 | 7 | 93 | -5 | 84 | 7 | 91 | 11 | 91 | -1 | 91 | 8 | 79 | -14 | 85 | -12 | 83 | 7 | 57 | -36 | 94 | 8 | 87 | 15 | 85 | 1 |
| **HU67** | 88 | -3 | 91 | -8 | 69 | 4 | 93 | -3 | 93 | -4 | 84 | -15 | 91 | -9 | 91 | -4 | 89 | 4 | 16 | -8 | 86 | -2 | 82 | -18 | 83 | -10 | 95 | -5 | 85 | -15 | 73 | -21 |
| **HU68** | 92 | -8 | 91 | -9 | 76 | -7 | 96 | -4 | 93 | -7 | 86 | -2 | 86 | -14 | 91 | -5 | 68 | -23 | 85 | -9 | 86 | -6 | 86 | 3 | 75 | 38 | 90 | -10 | 85 | 4 | 76 | 33 |
| **HU69** | 92 | 6 | 91 | 5 | 89 | 21 | 87 | 14 | 93 | 1 | 82 | -4 | 88 | 15 | 91 | -1 | 91 | 0 | 36 | 14 | 85 | 8 | 87 | 14 | 73 | 42 | 89 | -1 | 85 | 11 | 68 | 34 |
| **HU7** | 92 | -3 | 91 | -8 | 62 | 33 | 94 | 4 | 93 | -7 | 82 | -5 | 88 | -1 | 91 | -2 | 91 | -5 | 78 | -20 | 85 | 3 | 79 | -6 | 90 | -5 | 95 | -5 | 61 | 45 | 84 | -5 |
| **HU70** | 88 | 2 | 91 | -4 | 77 | 1 | 88 | -2 | 93 | -4 | 76 | 4 | 89 | -11 | 91 | -4 | 91 | -9 | 27 | 19 | 85 | -4 | 83 | -2 | 76 | -1 | 88 | -5 | 85 | 11 | 76 | 2 |
| **HU71** | 84 | -16 | 91 | 11 | 84 | -16 | 71 | -14 | 93 | -2 | 86 | 6 | 78 | -2 | 91 | -2 | 91 | -9 | 59 | -21 | 86 | 5 | 77 | 1 | 52 | -37 | 80 | -3 | 88 | 4 | 81 | -1 |
| **HU72** | 92 | -6 | 91 | 0 | 54 | 1 | 94 | 14 | 93 | 4 | 78 | -14 | 91 | 6 | 91 | -3 | 79 | 20 | 92 | 30 | 86 | 27 | 81 | -2 | 89 | -4 | 95 | 15 | 82 | 7 | 85 | -4 |
| **HU73** | 58 | 19 | 91 | 4 | 89 | 29 | 94 | 1 | 93 | -7 | 84 | 14 | 79 | 11 | 91 | 4 | 92 | 17 | 55 | 34 | 86 | 7 | 84 | 4 | 93 | 7 | 96 | -1 | 82 | 4 | 84 | 12 |
| **HU74** | 92 | -7 | 91 | 4 | 72 | -5 | 87 | -2 | 93 | -7 | 84 | 6 | 90 | 9 | 91 | 1 | 91 | 5 | 81 | -7 | 86 | -6 | 87 | 9 | 89 | 0 | 92 | -3 | 75 | 4 | 85 | 10 |
| **HU75** | 85 | 22 | 91 | -2 | 84 | 5 | 94 | -6 | 87 | -13 | 83 | 14 | 88 | 3 | 91 | 5 | 92 | 5 | 46 | 23 | 86 | -3 | 85 | 3 | 65 | 39 | 96 | -4 | 81 | 48 | 77 | 63 |
| **HU76** | 92 | 1 | 91 | -9 | 61 | -5 | 93 | 2 | 93 | -7 | 85 | 8 | 94 | -6 | 73 | -18 | 91 | 9 | 59 | -26 | 86 | 2 | 86 | -3 | 87 | 3 | 89 | -1 | 82 | 4 | 83 | 13 |
| **HU77** | 92 | -8 | 91 | -3 | 72 | 8 | 84 | -16 | 93 | -7 | 84 | -16 | 96 | 12 | 91 | 4 | 92 | -2 | 36 | 30 | 85 | 2 | 85 | 8 | 79 | 0 | 97 | -3 | 86 | -14 | 83 | -13 |
| **HU78** | 59 | -14 | 91 | 8 | 85 | 3 | 84 | -7 | 87 | -12 | 90 | -10 | 88 | -6 | 72 | -6 | 91 | -9 | 40 | 30 | 85 | -2 | 85 | 5 | 90 | 17 | 84 | -11 | 80 | 6 | 85 | 1 |
| **HU79** | 92 | 0 | 91 | 1 | 72 | 46 | 96 | -4 | 93 | 10 | 81 | -5 | 86 | 8 | 69 | 56 | 91 | -9 | 79 | 13 | 86 | 20 | 78 | -1 | 94 | -3 | 90 | -10 | 57 | 53 | 85 | 3 |
| **HU80** | 79 | 15 | 91 | -9 | 88 | 6 | 89 | -11 | 93 | -2 | 82 | 1 | 95 | -5 | 91 | 9 | 91 | 9 | 66 | 19 | 86 | -2 | 87 | -2 | 81 | -7 | 73 | -8 | 64 | -7 | 79 | -3 |
| **HU81** | 92 | 2 | 91 | 4 | 48 | 28 | 94 | -4 | 93 | -7 | 84 | 2 | 92 | 18 | 91 | -5 | 91 | 3 | 91 | -9 | 85 | -10 | 86 | 3 | 93 | -1 | 97 | 6 | 89 | 3 | 85 | 0 |
| **HU82** | 83 | -17 | 91 | -9 | 74 | -3 | 82 | 7 | 93 | 6 | 88 | -12 | 80 | -20 | 60 | -28 | 81 | 1 | 32 | -7 | 58 | 50 | 71 | -16 | 71 | -15 | 84 | 4 | 52 | -19 | 57 | -33 |
| **HU83** | 92 | 1 | 82 | 73 | 82 | 9 | 96 | -4 | 93 | -7 | 84 | 5 | 86 | 8 | 91 | -4 | 65 | 48 | 88 | -5 | 86 | -7 | 67 | 59 | 88 | -8 | 95 | -5 | 84 | -9 | 85 | 10 |
| **HU84** | 86 | -4 | 91 | -9 | 77 | 3 | 93 | -7 | 93 | -7 | 82 | 7 | 89 | -11 | 91 | -2 | 89 | 5 | 16 | 10 | 86 | -2 | 83 | -2 | 80 | -6 | 93 | -5 | 86 | 16 | 78 | 5 |
| **HU85** | 78 | 19 | 91 | -7 | 42 | 12 | 93 | -7 | 94 | 33 | 84 | 2 | 95 | -5 | 62 | 47 | 91 | -1 | 26 | 8 | 86 | -10 | 87 | -2 | 58 | -17 | 56 | 6 | 87 | -3 | 50 | -16 |
| **HU9** | 85 | 23 | 91 | -3 | 81 | 0 | 93 | -6 | 93 | -6 | 85 | -6 | 77 | -11 | 91 | -5 | 91 | -9 | 49 | -30 | 85 | 4 | 80 | -4 | 73 | -27 | 93 | -3 | 81 | -14 | 72 | -22 |
| **HU93** | 92 | -3 | 91 | -3 | 77 | 9 | 94 | -6 | 93 | -6 | 83 | -3 | 82 | 13 | 91 | -7 | 72 | -8 | 94 | 8 | 86 | 2 | 84 | -14 | 87 | -10 | 96 | -2 | 82 | -11 | 85 | -8 |
| **HU94** | 92 | 5 | 91 | -9 | 79 | 10 | 95 | -5 | 93 | -7 | 83 | 5 | 94 | -6 | 91 | -4 | 91 | 10 | 66 | 33 | 85 | -1 | 84 | -6 | 94 | 1 | 99 | -1 | 83 | -17 | 86 | 8 |
| **HU95** | 92 | -8 | 91 | -5 | 78 | 7 | 92 | -8 | 93 | -7 | 85 | 15 | 79 | 1 | 73 | -22 | 91 | -5 | 76 | -13 | 86 | 7 | 83 | 1 | 83 | -3 | 81 | -19 | 79 | -9 | 83 | 14 |
| **HU97** | 87 | -14 | 91 | -7 | 83 | -17 | 94 | 4 | 88 | 0 | 90 | -10 | 92 | -8 | 91 | 1 | 91 | -6 | 77 | 0 | 86 | 7 | 83 | -5 | 89 | 5 | 94 | 16 | 81 | 7 | 84 | -16 |

**Table S4**: Observed scores for the set of 50 cocktail testing set of strains. Phage are grouped and colour coded according to their activity groupings (Figure 2). Grey highlighted phage names are the phage used in the General Cocktail. The red coloured scores show the phage chosen for the Bespoke Cocktail for each strain. The final column shows the total number of phage that have an activity score of less than 60 for each strain (highlighted in green). Yellow highlighted boxes show score of between 61 and 80 which were selected for a bespoke cocktail if no phage scoring less than 60 in a grouping were available.

| Isolate | CHAP1 | D11 AU | BAT1 | ALD2 | D4 | NEA2 | E4 | F1 | GWF | FAT | TB54 | LUC04 | A01 | ALDi98 | ALD1 | HAM3 | TB69 | AB19 | Geo | OIL | D3 | BO1 | UFA5s | RV2 | HAM53 | A | N | P | E | LEG | BOW | Total active phage |
|---|---|---|---|---|---|---|---|---|---|---|---|---|---|---|---|---|---|---|---|---|---|---|---|---|---|---|---|---|---|---|---|---|
| CAN08 | 91 | 99 | 84 | 100 | 85 | 100 | 100 | 98 | 100 | 100 | 100 | 83 | 100 | 100 | 81 | 100 | 93 | 98 | 67 | 81 | 90 | 71 | 41 | 23 | 22 | 57 | 85 | 41 | 49 | 89 | 92 | 6 |
| CAN11 | 100 | 100 | 94 | 94 | 83 | 96 | 94 | 91 | 101 | 100 | 92 | 95 | 89 | 89 | 87 | 83 | 83 | 84 | 100 | 95 | 98 | 100 | 88 | 92 | 82 | 92 | 98 | 100 | 94 | 100 | 98 | 0 |
| CAN111 | 23 | 93 | 66 | 34 | 78 | 43 | 28 | 83 | 100 | 100 | 98 | 97 | 96 | 98 | 87 | 95 | 98 | 100 | 87 | 98 | 87 | 87 | 86 | 88 | 80 | 92 | 89 | 100 | 100 | 96 | 99 | 4 |
| CAN116 | 95 | 99 | 73 | 75 | 27 | 59 | 91 | 86 | 96 | 100 | 100 | 100 | 100 | 96 | 100 | 100 | 97 | 81 | 90 | 100 | 97 | 94 | 85 | 100 | 91 | 100 | 100 | 87 | 90 | 101 | 100 | 2 |
| CAN118 | 97 | 100 | 100 | 92 | 76 | 94 | 100 | 100 | 100 | 90 | 100 | 91 | 100 | 100 | 86 | 100 | 97 | 94 | 100 | 100 | 100 | 100 | 95 | 87 | 100 | 99 | 100 | 81 | 87 | 77 | 92 | 0 |
| CAN119 | 31 | 21 | 43 | 71 | 24 | 27 | 39 | 45 | 57 | 64 | 29 | 26 | 38 | 81 | 69 | 84 | 51 | 56 | 80 | 96 | 57 | 43 | 27 | 33 | 61 | 89 | 81 | 53 | 75 | 53 | 81 | 19 |
| CAN127 | 21 | 89 | 87 | 38 | 56 | 30 | 37 | 16 | 86 | 91 | 57 | 94 | 92 | 89 | 85 | 84 | 64 | 40 | 12 | 80 | 84 | 41 | 74 | 50 | 7 | 49 | 82 | 40 | 41 | 43 | 42 | 17 |
| CAN130 | 100 | 87 | 90 | 100 | 72 | 94 | 93 | 91 | 94 | 91 | 84 | 97 | 95 | 90 | 93 | 93 | 73 | 74 | 79 | 95 | 97 | 88 | 89 | 93 | 85 | 95 | 88 | 89 | 88 | 100 | 99 | 0 |
| CAN137 | 39 | 89 | 88 | 72 | 53 | 51 | 94 | 100 | 85 | 74 | 82 | 100 | 100 | 96 | 89 | 97 | 78 | 72 | 85 | 91 | 88 | 98 | 97 | 66 | 100 | 90 | 94 | 56 | 71 | 57 | 90 | 5 |
| CAN144 | 91 | 95 | 100 | 92 | 83 | 97 | 96 | 86 | 84 | 82 | 90 | 89 | 100 | 96 | 81 | 97 | 78 | 81 | 91 | 90 | 83 | 89 | 81 | 83 | 82 | 87 | 88 | 84 | 89 | 93 | 98 | 0 |
| CAN154 | 96 | 85 | 97 | 94 | 90 | 96 | 89 | 87 | 101 | 100 | 91 | 93 | 99 | 96 | 97 | 96 | 93 | 101 | 89 | 93 | 97 | 89 | 90 | 92 | 87 | 97 | 91 | 93 | 97 | 100 | 100 | 0 |
| CAN156 | 74 | 93 | 100 | 96 | 83 | 85 | 77 | 87 | 82 | 8 | 41 | 86 | 87 | 88 | 79 | 80 | 71 | 27 | 88 | 82 | 72 | 83 | 74 | 77 | 80 | 8 | 15 | 5 | 5 | 22 | 28 | 9 |
| CAN159 | 90 | 95 | 100 | 100 | 64 | 88 | 100 | 83 | 88 | 86 | 90 | 88 | 100 | 97 | 76 | 100 | 81 | 81 | 86 | 96 | 74 | 91 | 64 | 79 | 69 | 91 | 86 | 69 | 78 | 74 | 94 | 0 |
| CAN163 | 18 | 86 | 92 | 92 | 83 | 91 | 95 | 94 | 101 | 95 | 98 | 94 | 100 | 100 | 91 | 100 | 94 | 88 | 91 | 91 | 87 | 97 | 91 | 92 | 92 | 91 | 95 | 85 | 91 | 88 | 99 | 1 |
| CAN165 | 29 | 50 | 31 | 72 | 99 | 29 | 77 | 97 | 89 | 100 | 97 | 92 | 95 | 96 | 89 | 96 | 100 | 99 | 67 | 68 | 99 | 94 | 94 | 66 | 87 | 78 | 67 | 90 | 93 | 100 | 97 | 4 |
| CAN166 | 17 | 85 | 83 | 79 | 84 | 75 | 93 | 99 | 84 | 91 | 87 | 99 | 97 | 94 | 99 | 90 | 80 | 86 | 100 | 77 | 98 | 100 | 90 | 100 | 93 | 95 | 100 | 81 | 85 | 100 | 100 | 1 |
| CAN167 | 14 | 86 | 9 | 22 | 7 | 11 | 12 | 90 | 88 | 1 | 79 | 91 | 100 | 99 | 79 | 100 | 83 | 69 | 77 | 12 | 71 | 35 | 14 | 15 | 15 | 5 | 10 | 15 | 13 | 8 | 13 | 18 |
| CAN173 | 100 | 85 | 91 | 100 | 86 | 98 | 86 | 81 | 95 | 87 | 100 | 91 | 100 | 100 | 82 | 100 | 98 | 96 | 92 | 98 | 87 | 100 | 75 | 86 | 73 | 39 | 94 | 29 | 90 | 20 | 30 | 4 |
| CAN174 | 20 | 81 | 23 | 77 | 24 | 28 | 25 | 100 | 86 | 88 | 77 | 93 | 91 | 95 | 85 | 87 | 74 | 83 | 100 | 76 | 83 | 69 | 82 | 92 | 71 | 27 | 23 | 23 | 20 | 38 | 24 | 11 |
| CAN182 | 45 | 89 | 90 | 80 | 72 | 80 | 97 | 91 | 80 | 77 | 96 | 92 | 100 | 86 | 91 | 39 | 37 | 43 | 59 | 79 | 96 | 95 | 93 | 86 | 87 | 94 | 96 | 78 | 89 | 82 | 93 | 5 |
| CAN186 | 100 | 88 | 100 | 100 | 91 | 100 | 96 | 96 | 92 | 94 | 80 | 92 | 86 | 99 | 97 | 85 | 73 | 85 | 98 | 100 | 90 | 100 | 99 | 90 | 87 | 96 | 100 | 94 | 94 | 100 | 100 | 0 |
| CAN188 | 9 | 12 | 38 | 92 | 45 | 18 | 28 | 88 | 40 | 99 | 75 | 92 | 100 | 100 | 84 | 99 | 11 | 52 | 39 | 16 | 80 | 22 | 40 | 49 | 32 | 85 | 100 | 73 | 74 | 32 | 44 | 17 |
| CAN199 | 99 | 64 | 94 | 100 | 95 | 96 | 99 | 77 | 90 | 100 | 84 | 87 | 86 | 92 | 68 | 83 | 85 | 86 | 100 | 99 | 68 | 100 | 63 | 72 | 66 | 83 | 100 | 100 | 100 | 96 | 99 | 0 |
| CAN20 | 100 | 74 | 98 | 100 | 92 | 100 | 71 | 91 | 89 | 94 | 88 | 91 | 98 | 95 | 84 | 92 | 66 | 46 | 30 | 57 | 89 | 54 | 76 | 79 | 14 | 83 | 100 | 94 | 88 | 97 | 92 | 5 |
| CAN203 | 92 | 98 | 92 | 97 | 80 | 93 | 93 | 92 | 79 | 100 | 78 | 88 | 95 | 88 | 87 | 85 | 77 | 77 | 95 | 96 | 87 | 100 | 83 | 87 | 85 | 87 | 100 | 84 | 91 | 100 | 100 | 0 |
| CAN24 | 32 | 100 | 40 | 29 | 85 | 100 | 100 | 92 | 84 | 91 | 70 | 92 | 71 | 88 | 86 | 69 | 72 | 76 | 100 | 100 | 83 | 100 | 80 | 36 | 85 | 97 | 100 | 100 | 98 | 97 | 99 | 4 |
| CAN49 | 17 | 53 | 17 | 77 | 50 | 18 | 26 | 95 | 96 | 95 | 94 | 84 | 98 | 99 | 79 | 96 | 95 | 99 | 58 | 62 | 93 | 74 | 74 | 80 | 63 | 24 | 22 | 46 | 45 | 58 | 45 | 13 |
| CAN73 | 30 | 100 | 43 | 54 | 20 | 33 | 18 | 94 | 84 | 98 | 76 | 91 | 84 | 81 | 91 | 80 | 72 | 87 | 100 | 100 | 89 | 89 | 86 | 70 | 30 | 26 | 37 | 27 | 28 | 45 | 21 | 13 |

| | | | | | | | | | | | | | | | | | | | | | | | | | | | | | | | | |
|---|---|---|---|---|---|---|---|---|---|---|---|---|---|---|---|---|---|---|---|---|---|---|---|---|---|---|---|---|---|---|---|---|
| **CAN74** | **24** | 100 | 34 | 30 | 45 | 25 | 36 | **71** | 99 | 100 | 99 | 95 | 98 | 101 | 88 | 99 | 96 | 89 | **78** | 100 | **29** | 48 | 35 | 42 | **77** | 94 | 100 | 92 | 95 | 83 | 100 | 10 |
| **CAN76** | **97** | 100 | 31 | 89 | 61 | **16** | 98 | 100 | **5** | 78 | 76 | 99 | 100 | 93 | 100 | 93 | **71** | 71 | 100 | 100 | 98 | 81 | 89 | 96 | 85 | 98 | 100 | 67 | 86 | **64** | 94 | 3 |
| **CAN80** | **22** | 100 | 100 | 100 | **29** | 39 | 88 | 93 | 91 | 89 | 91 | 97 | 100 | 101 | 97 | 100 | **35** | 86 | 100 | 100 | 97 | **68** | 82 | 89 | 90 | 98 | 100 | 73 | 86 | **62** | 96 | 4 |
| **CAN85** | 90 | 100 | 100 | 100 | 80 | 92 | 98 | 93 | 98 | 97 | 92 | 90 | 99 | 100 | 91 | 99 | 38 | 19 | 38 | 100 | 88 | 100 | 65 | 81 | 71 | 100 | 100 | 97 | 96 | 95 | 100 | 3 |
| **CAN90** | 92 | 92 | 100 | 100 | 100 | 91 | 91 | 82 | 93 | 100 | 78 | 91 | 86 | 95 | 81 | **28** | 76 | **57** | **34** | 100 | 73 | **58** | 69 | 82 | **65** | 90 | 100 | 100 | 99 | 100 | 100 | 4 |
| **CAN91** | **15** | 100 | 71 | 96 | **12** | 15 | 15 | 91 | 85 | 96 | 86 | 91 | 100 | 97 | 93 | 95 | 89 | 59 | **21** | 100 | 99 | **20** | **17** | 28 | 86 | 95 | 92 | 88 | 92 | 88 | 95 | 9 |
| **CAN96** | **29** | 100 | 31 | 29 | 80 | **27** | 27 | 86 | 92 | 84 | 81 | 96 | 96 | 94 | 90 | 94 | **75** | 82 | 100 | 100 | **39** | 97 | 73 | **28** | 82 | 92 | 100 | 80 | 86 | 89 | 97 | 7 |
| **HB01** | 78 | 87 | 85 | 80 | 56 | 52 | **38** | 100 | **31** | 85 | 90 | 98 | 100 | 100 | 100 | 100 | 81 | 82 | 100 | **29** | 100 | 100 | 98 | 100 | 91 | 100 | 100 | **74** | 86 | 84 | 97 | 5 |
| **HB08** | 90 | 75 | 73 | 93 | 65 | 59 | 61 | 76 | 30 | 93 | 100 | 82 | 100 | 100 | 89 | 100 | 90 | 92 | 70 | 23 | 82 | 84 | 78 | 87 | 86 | 97 | 76 | 85 | 87 | 100 | 100 | 3 |
| **HU03** | 90 | **18** | 100 | 100 | 72 | 93 | 97 | **20** | 36 | 83 | 30 | 25 | 29 | 101 | 91 | 71 | 81 | 29 | **28** | 100 | 95 | **44** | 98 | 93 | **39** | 90 | 88 | 70 | 84 | 79 | 90 | 10 |
| **HU11** | 28 | **25** | 35 | 32 | 26 | **14** | 30 | 92 | 73 | 81 | **62** | 94 | 89 | 80 | 86 | 76 | **55** | 59 | 71 | 76 | 82 | 90 | 86 | 89 | 90 | 87 | **75** | 77 | 86 | 78 | 92 | 9 |
| **HU12** | 88 | 26 | 92 | 94 | 74 | 90 | 30 | 98 | 100 | 90 | 100 | 100 | 100 | 100 | 91 | 100 | 100 | 96 | 91 | 95 | 79 | 93 | 85 | 99 | 81 | 98 | 93 | 79 | 82 | 83 | 96 | 2 |
| **HU17** | **16** | 82 | 69 | **46** | 82 | **9** | **18** | 92 | 100 | 92 | 100 | 92 | 100 | 99 | 84 | 100 | 97 | 83 | 82 | 89 | 89 | 95 | 86 | 90 | 87 | 88 | 90 | 91 | 89 | 97 | 96 | 4 |
| **HU22** | 70 | 13 | 84 | 80 | 75 | 77 | 94 | 93 | 68 | 88 | 76 | 94 | 81 | 52 | 89 | 84 | 67 | 74 | 94 | 76 | 86 | 90 | 88 | 90 | 92 | 87 | 91 | 85 | 87 | 96 | 98 | 2 |
| **HU25** | 91 | 33 | 81 | 100 | 69 | 92 | 34 | 97 | 100 | 93 | 94 | 90 | 100 | 100 | 77 | 100 | 75 | 79 | 76 | 98 | 68 | 97 | 65 | 69 | 68 | 93 | 91 | 86 | 91 | 100 | 100 | 2 |
| **HU31** | 21 | 82 | 78 | 86 | 71 | 41 | 85 | 89 | 82 | 80 | 86 | 93 | 100 | 96 | 81 | 97 | 77 | 73 | 75 | 79 | 82 | 100 | 86 | 90 | 86 | 87 | 100 | 75 | 88 | 66 | 88 | 2 |
| **HU32** | 70 | 8 | 63 | 68 | 70 | 57 | 90 | 16 | 96 | 100 | 100 | 89 | 100 | 93 | 67 | 100 | 77 | 73 | 91 | 61 | 74 | 100 | 72 | 97 | 74 | 100 | 87 | 83 | 87 | 100 | 100 | 3 |
| **HU36** | **58** | 53 | 57 | 87 | 65 | **50** | 95 | 92 | 100 | 93 | 100 | 95 | 100 | 83 | 93 | 100 | 100 | 95 | **53** | 77 | 87 | 88 | 86 | 94 | 80 | 94 | 70 | 82 | **66** | 84 | 92 | 5 |
| **HU58** | **30** | **25** | 88 | 90 | 69 | 92 | 95 | **14** | 81 | 98 | 77 | 95 | 87 | 22 | 100 | 74 | **38** | 70 | 83 | 93 | 96 | 94 | 91 | 94 | 87 | 99 | 91 | 84 | 86 | 100 | 100 | 5 |
| **HU65** | 20 | 36 | 64 | 90 | 25 | **16** | 24 | 91 | **54** | 72 | 84 | 91 | 66 | 100 | 93 | 100 | 65 | 64 | 65 | **3** | 88 | 31 | 28 | **24** | 84 | 33 | **31** | 33 | 33 | 54 | 36 | 16 |
| **HU84** | **5** | **19** | 37 | **18** | 22 | **6** | 32 | 90 | 100 | 100 | 99 | 93 | 100 | 97 | 99 | 100 | **69** | 69 | 74 | 87 | 84 | 84 | 73 | 87 | 76 | 92 | 84 | 85 | 95 | 100 | 100 | 7 |
| **HU85** | **14** | 93 | 59 | 56 | 70 | 18 | 24 | 59 | 100 | 98 | 50 | **15** | **24** | 100 | 92 | 60 | 90 | 81 | **30** | 95 | **22** | 61 | 67 | 74 | 82 | 94 | 93 | 89 | 91 | 100 | 100 | 11 |

**Table S5**: Results of cocktail testing against the set of 50 strains. The individual scores for each phage in the General Cocktail also shown for comparison. Green highlighted boxes show scores less than 60 and yellow highlighted for scores 61 – 80 (for the individual phage scores). The bespoke cocktails for each strain were composed of the phage shown in red text in Table S4.

| Isolate | CHAP1 | F1 | TB69 | RV2 | P | Active phage in Gen cocktail | General Cocktail Score | St.Dev | Bespoke Cocktail Score | St.Dev |
|---|---|---|---|---|---|---|---|---|---|---|
| CAN08 | 91 | 98 | 93 | 23 | 41 | 2 | 36 | 0.6 | 26 | 1.5 |
| CAN11 | 100 | 91 | 83 | 92 | 100 | 0 | 96 | 4.8 | | |
| CAN111 | 23 | 83 | 98 | 88 | 100 | 1 | 46 | 2.4 | 43 | 0.4 |
| CAN116 | 95 | 86 | 97 | 100 | 87 | 0 | 71 | 8.4 | 54 | 0.8 |
| CAN118 | 97 | 100 | 97 | 87 | 81 | 0 | 93 | 4.0 | | |
| CAN119 | 31 | 45 | 51 | 33 | 53 | 5 | 37 | 2.2 | 32 | 0.1 |
| CAN127 | 21 | 16 | 64 | 50 | 40 | 4 | 33 | 2.4 | 9 | 2.0 |
| CAN130 | 100 | 91 | 73 | 93 | 89 | 0 | 95 | 14.4 | | |
| CAN137 | 39 | 100 | 78 | 66 | 56 | 2 | 56 | 1.4 | 53 | 0.7 |
| CAN144 | 91 | 86 | 78 | 83 | 84 | 0 | 92 | 10.2 | | |
| CAN154 | 96 | 87 | 93 | 92 | 93 | 0 | 95 | 1.2 | | |
| CAN156 | 74 | 87 | 71 | 77 | 5 | 1 | 11 | 2.4 | 11 | 3.1 |
| CAN159 | 90 | 83 | 81 | 79 | 69 | 0 | 90 | 7.9 | | |
| CAN163 | 18 | 94 | 94 | 92 | 85 | 1 | 29 | 0.5 | | |
| CAN165 | 29 | 97 | 100 | 66 | 90 | 1 | 50 | 2.2 | 47 | 0.8 |
| CAN166 | 17 | 99 | 80 | 100 | 81 | 1 | 29 | 0.3 | | |
| CAN167 | 14 | 90 | 83 | 15 | 15 | 3 | 9 | 2.8 | 4 | 0.3 |
| CAN173 | 100 | 81 | 98 | 86 | 29 | 1 | 26 | 0.0 | 29 | 0.1 |
| CAN174 | 20 | 100 | 74 | 92 | 23 | 2 | 24 | 0.2 | 24 | 0.4 |
| CAN182 | 45 | 91 | 37 | 86 | 78 | 2 | 62 | 1.6 | 40 | 2.8 |
| CAN186 | 100 | 96 | 73 | 90 | 94 | 0 | 99 | 4.9 | | |
| CAN188 | 9 | 88 | 11 | 49 | 73 | 3 | 4 | 0.1 | 4 | 0.8 |
| CAN199 | 99 | 77 | 85 | 72 | 100 | 0 | 91 | 13.8 | | |
| CAN20 | 100 | 91 | 66 | 79 | 94 | 0 | 42 | 0.7 | 12 | 0.9 |
| CAN203 | 92 | 92 | 77 | 87 | 84 | 0 | 95 | 0.8 | | |
| CAN24 | 32 | 92 | 72 | 36 | 100 | 2 | 37 | 0.6 | 33 | 0.7 |
| CAN49 | 17 | 95 | 95 | 80 | 46 | 2 | 31 | 1.0 | 24 | 1.1 |
| CAN73 | 30 | 94 | 72 | 70 | 27 | 2 | 31 | 0.5 | 32 | 0.8 |
| CAN74 | 24 | 71 | 96 | 42 | 92 | 2 | 36 | 1.6 | 30 | 1.3 |
| CAN76 | 97 | 100 | 71 | 96 | 67 | 0 | 93 | 0.3 | 31 | 1.4 |
| CAN80 | 22 | 93 | 35 | 89 | 73 | 2 | 31 | 0.2 | 31 | 0.6 |
| CAN85 | 90 | 93 | 38 | 81 | 97 | 1 | 76 | 2.6 | 37 | 12.1 |
| CAN90 | 92 | 82 | 76 | 82 | 100 | 0 | 85 | 0.7 | 33 | 0.2 |
| CAN91 | 15 | 91 | 89 | 28 | 88 | 2 | 17 | 0.2 | 14 | 1.9 |
| CAN96 | 29 | 86 | 75 | 28 | 80 | 2 | 34 | 1.9 | 16 | 4.0 |
| HB01 | 78 | 100 | 81 | 100 | 74 | 0 | 100 | 2.5 | 22 | 1.7 |
| HB08 | 90 | 76 | 90 | 87 | 85 | 0 | 88 | 3.3 | 36 | 0.4 |
| HU03 | 90 | 20 | 81 | 93 | 70 | 1 | 29 | 0.4 | 18 | 0.6 |
| HU11 | 28 | 92 | 55 | 89 | 77 | 2 | 18 | 0.1 | 26 | 0.5 |
| HU12 | 88 | 98 | 100 | 99 | 79 | 0 | 95 | 4.5 | | |
| HU17 | 16 | 92 | 97 | 90 | 91 | 1 | 23 | 6.9 | 15 | 0.1 |
| HU22 | 70 | 93 | 67 | 90 | 85 | 0 | 69 | 1.1 | 15 | 1.9 |
| HU25 | 91 | 97 | 75 | 69 | 86 | 0 | 95 | 15.9 | 38 | 5.2 |
| HU31 | 21 | 89 | 77 | 90 | 75 | 1 | 45 | 1.9 | 39 | 1.5 |
| HU32 | 70 | 16 | 77 | 97 | 83 | 1 | 9 | 0.6 | 7 | 1.3 |
| HU36 | 58 | 92 | 100 | 94 | 82 | 1 | 62 | 1.8 | 60 | 2.0 |
| HU58 | 30 | 14 | 38 | 94 | 84 | 3 | 13 | 0.6 | 2 | 0.3 |
| HU65 | 20 | 91 | 65 | 24 | 33 | 3 | 28 | 0.4 | 16 | 0.0 |
| HU84 | 5 | 90 | 69 | 87 | 85 | 1 | 48 | 1.0 | 21 | 4.3 |
| HU85 | 14 | 59 | 90 | 74 | 89 | 2 | 16 | 0.8 | 8 | 4.1 |

**Table S6:** Scores for predicted and observed phage activity for the validation set of 'unseen' strains (n=10). Phage names are colour coded as before to show phage activity groupings. F1 scores to indicate accuracy of the model with F1 scores over 0.6 highlighted in green. The predicted and observed scores are shown for each strain side by side. The scores in blue text denote the phage selected for each bespoke cocktail. Following the rules set for cocktail composition we also noted that the majority of predictive cocktails did contain a minimum of 3 phage from the models with higher F1 scores.

| | | HU256 | | HU258 | | HU260 | | HU263 | | HU270 | | HU98 | | HU99 | | DTU09 | | DTU14 | | DTU16 | |
|---|---|---|---|---|---|---|---|---|---|---|---|---|---|---|---|---|---|---|---|---|---|
| Phage | f1 | Predicted | Observed | Predicted | Observed | Predicted | Observed | Predicted | Observed | Predicted | Observed | Predicted | Observed | Predicted | Observed | Predicted | Observed | Predicted | Observed | Predicted | Observed |
| A01 | 0.38 | 94 | 78 | 94 | 84 | 94 | 68 | 94 | 87 | 94 | 91 | 94 | 100 | 94 | 95 | 94 | 97 | 94 | 100 | 94 | 93 |
| AB19 | 0.31 | 61 | 91 | 77 | 85 | 68 | 98 | 77 | 92 | 74 | 96 | 83 | 99 | 84 | 92 | 70 | 86 | 83 | 100 | 68 | 95 |
| ACH4 | 0.61 | 92 | 94 | 93 | 67 | 94 | 81 | 73 | 83 | 94 | 99 | 80 | 95 | 90 | 74 | 70 | 98 | 91 | 56 | 26 | 42 |
| ALD1 | NA | 89 | 89 | 87 | 73 | 87 | 68 | 86 | 86 | 87 | 87 | 89 | 100 | 88 | 83 | 86 | 94 | 88 | 100 | 83 | 93 |
| ALD2 | 0.68 | 53 | 94 | 75 | 63 | 27 | 32 | 44 | 42 | 28 | 27 | 77 | 80 | 76 | 72 | 78 | 88 | 75 | 96 | 73 | 53 |
| ALDi98 | 0.13 | 43 | 100 | 95 | 80 | 95 | 88 | 95 | 100 | 95 | 91 | 95 | 97 | 95 | 100 | 95 | 93 | 95 | 100 | 95 | 92 |
| BAT1 | 0.70 | 77 | 82 | 62 | 52 | 28 | 44 | 46 | 36 | 37 | 51 | 67 | 48 | 61 | 72 | 62 | 85 | 70 | 99 | 29 | 39 |
| BO1 | 0.30 | 71 | 81 | 84 | 67 | 76 | 66 | 68 | 31 | 70 | 100 | 93 | 74 | 76 | 95 | 55 | 88 | 91 | 100 | 63 | 88 |
| BOW | 0.72 | 94 | 99 | 94 | 76 | 93 | 95 | 90 | 100 | 93 | 97 | 92 | 96 | 95 | 71 | 79 | 91 | 76 | 99 | 33 | 37 |
| CHAP1 | 0.83 | 44 | 24 | 31 | 40 | 23 | 33 | 33 | 32 | 18 | 74 | 70 | 96 | 89 | 75 | 71 | 94 | 92 | 100 | 32 | 34 |
| D11 AU | 0.74 | 44 | 35 | 59 | 39 | 58 | 69 | 56 | 79 | 56 | 90 | 75 | 97 | 82 | 50 | 77 | 50 | 79 | 99 | 77 | 90 |
| D3 | 0.59 | 82 | 84 | 70 | 73 | 64 | 23 | 39 | 24 | 65 | 34 | 89 | 100 | 89 | 89 | 75 | 94 | 89 | 99 | 72 | 62 |
| D4 | 0.67 | 58 | 84 | 53 | 72 | 45 | 75 | 55 | 87 | 51 | 99 | 68 | 98 | 72 | 100 | 58 | 94 | 63 | 100 | 42 | 49 |
| E4 | 0.75 | 73 | 100 | 68 | 77 | 70 | 50 | 41 | 75 | 48 | 98 | 86 | 98 | 85 | 77 | 89 | 93 | 87 | 100 | 34 | 47 |
| EBB3 | 0.70 | 88 | 95 | 93 | 82 | 88 | 99 | 83 | 97 | 88 | 99 | 74 | 96 | 89 | 73 | 49 | 91 | 89 | 43 | 35 | 39 |
| F1 | 0.57 | 46 | 88 | 84 | 86 | 88 | 94 | 84 | 88 | 92 | 95 | 78 | 78 | 92 | 82 | 92 | 90 | 87 | 99 | 92 | 93 |
| FAT | 0.47 | 91 | 77 | 91 | 83 | 91 | 68 | 91 | 86 | 91 | 92 | 91 | 97 | 91 | 97 | 91 | 91 | 72 | 100 | 91 | 93 |
| Geo | 0.58 | 63 | 88 | 61 | 28 | 54 | 91 | 50 | 78 | 63 | 100 | 67 | 94 | 67 | 94 | 49 | 82 | 53 | 100 | 65 | 93 |
| GWF | 0.58 | 89 | 81 | 95 | 69 | 80 | 80 | 95 | 82 | 93 | 86 | 94 | 100 | 94 | 89 | 81 | 89 | 92 | 100 | 89 | 90 |
| HAM3 | NA | 93 | 82 | 93 | 74 | 93 | 63 | 93 | 79 | 93 | 97 | 93 | 97 | 93 | 82 | 72 | 93 | 93 | 99 | 93 | 91 |
| HAM53 | 0.71 | 86 | 91 | 88 | 50 | 80 | 69 | 88 | 89 | 86 | 100 | 84 | 99 | 85 | 82 | 33 | 99 | 83 | 100 | 28 | 49 |
| LEG | 0.67 | 91 | 98 | 93 | 73 | 93 | 100 | 75 | 97 | 91 | 99 | 91 | 98 | 80 | 73 | 78 | 94 | 73 | 100 | 48 | 49 |
| LUC04 | NA | 91 | 84 | 69 | 84 | 91 | 76 | 69 | 83 | 91 | 89 | 91 | 97 | 91 | 88 | 69 | 90 | 91 | 99 | 69 | 91 |
| NAB1 | 0.72 | 91 | 98 | 91 | 77 | 88 | 92 | 69 | 97 | 88 | 99 | 79 | 97 | 91 | 78 | 66 | 93 | 62 | 55 | 37 | 39 |
| NEA2 | 0.81 | 71 | 78 | 51 | 64 | 24 | 26 | 43 | 38 | 25 | 78 | 90 | 100 | 87 | 99 | 84 | 91 | 84 | 100 | 43 | 40 |
| OIL | 0.31 | 86 | 75 | 86 | 85 | 86 | 81 | 86 | 90 | 86 | 90 | 86 | 97 | 86 | 81 | 86 | 90 | 86 | 97 | 86 | 79 |
| POC2 | 0.71 | 85 | 96 | 86 | 78 | 83 | 92 | 72 | 97 | 83 | 99 | 80 | 88 | 83 | 74 | 54 | 94 | 83 | 43 | 39 | 40 |
| RV2 | 0.46 | 79 | 94 | 63 | 81 | 76 | 73 | 71 | 93 | 78 | 98 | 90 | 98 | 60 | 79 | 79 | 95 | 81 | 100 | 84 | 98 |
| TB54 | 0.48 | 80 | 91 | 86 | 85 | 88 | 85 | 74 | 93 | 89 | 99 | 96 | 93 | 95 | 95 | 71 | 88 | 95 | 98 | 71 | 75 |
| TB69 | 0.47 | 59 | 76 | 78 | 68 | 84 | 55 | 77 | 79 | 83 | 96 | 83 | 100 | 84 | 89 | 52 | 94 | 83 | 100 | 69 | 95 |
| UFA5s | 0.48 | 59 | 78 | 55 | 68 | 75 | 68 | 51 | 52 | 73 | 100 | 86 | 95 | 85 | 76 | 81 | 94 | 85 | 94 | 53 | 97 |

**Table S7:** Results from cocktail testing of the set of 10 'unseen' validation strains. Scores are shown from two duplicate experiments testing the general, bespoke predicted and bespoke observed cocktails in artificial urine (AU) media and canine urine (CU). The biological repeats were carried out using different batches of media. The data from experiment 1 is plotted in Figure 4D.

| | Experiment 1 | | | | | | Experiment 2 | | | | | |
|---|---|---|---|---|---|---|---|---|---|---|---|---|
| | AU | | | CU | | | AU | | | CU | | |
| Strain | General | Predicted | Observed | General | Predicted | Observed | General | Predicted | Observed | General | Predicted | Observed |
| HU256 | 42.2 | 22.0 | 21.1 | 59.2 | 21.6 | 19.8 | 13.4 | 14.8 | 11.3 | 47.1 | 15.3 | 16.6 |
| HU258 | 81.3 | 52.3 | 53.8 | 93.9 | 90.4 | 96.7 | 58.9 | 39.9 | 47.6 | 84.6 | 80.6 | 85.5 |
| HU260 | 43.1 | 20.7 | 36.8 | 65.5 | 42.6 | 34.8 | 32.7 | 16.8 | 9.9 | 41.6 | 13.0 | 16.0 |
| HU263 | 71.4 | 40.8 | 40.3 | 78.3 | 16.9 | 15.9 | 41.1 | 29.6 | 19.1 | 80.2 | 24.2 | 26.4 |
| HU270 | 99.3 | 51.8 | 33.3 | 89.8 | 30.7 | 37.4 | 105.5 | 46.0 | 22.3 | 89.4 | 24.7 | 39.7 |
| HU98 | 91.6 | 98.7 | 91.7 | 89.3 | 84.3 | 60.2 | 89.3 | 104.1 | 91.7 | 95.6 | 87.3 | 83.6 |
| HU99 | 100.4 | 100.5 | 90.7 | 102.7 | 95.3 | 95.4 | 104.6 | 102.1 | 93.6 | 97.6 | 98.6 | 99.9 |
| DTU09 | 99.0 | 98.9 | 99.6 | 96.3 | 96.9 | 97.9 | 80.7 | 84.8 | 84.6 | 97.5 | 99.9 | 103.0 |
| DTU14 | 61.7 | 61.1 | 24.8 | 100.0 | 98.2 | 113.4 | 84.7 | 92.2 | 75.2 | 100.9 | 93.6 | 88.2 |
| DTU16 | 22.1 | 23.0 | 12.1 | 50.9 | 53.3 | 48.9 | 21.6 | 21.6 | 28.6 | 39.9 | 98.1 | 80.2 |

Supplementary References

1. T. Brooks, C. W. Keevil, A simple artificial urine for the growth of urinary pathogens. *Lett. Appl. Microbiol.* **24**, 203–206 (1997).

2. N. Sarigul, F. Korkmaz, İ. Kurultak, A New Artificial Urine Protocol to Better Imitate Human Urine. *Sci. Rep.* **9**, 20159 (2019).
